# Age of the bone marrow dictates clonality of smooth muscle-derived cells in the atherosclerotic plaque

**DOI:** 10.1101/2022.01.18.476756

**Authors:** Inamul Kabir, Xinbo Zhang, Jui M. Dave, Raja Chakraborty, Rihao Qu, Rachana R. Chandran, Aglaia Ntokou, Eunate Gallardo-Vara, Binod Aryal, Noemi Rotllan, Rolando Garcia-Milian, John Hwa, Yuval Kluger, Kathleen A. Martin, Carlos Fernández-Hernando, Daniel M. Greif

## Abstract

Aging is the predominant risk factor for atherosclerosis, the leading cause of death. Rare smooth muscle cell (SMC) progenitors clonally expand giving rise to up to ∼70% of atherosclerotic plaque cells; however, the effect of age on SMC clonality is not known. Our results indicate that aged bone marrow (BM)-derived cells non-cell autonomously induce SMC polyclonality and worsen atherosclerosis. Indeed, in myeloid cells from aged mice and humans, TET2 levels are reduced which epigenetically silences integrin β3 resulting in increased tumor necrosis factor [TNF]-α signaling. In turn, TNFα signals through TNF receptor 1 on SMCs to promote proliferation and induces recruitment and expansion of multiple SMC progenitors into the atherosclerotic plaque. Notably, integrin β3 overexpression in aged BM preserves dominance of the lineage of a single SMC progenitor and attenuates plaque burden. Our results demonstrate a molecular mechanism of aged macrophage-induced SMC polyclonality and atherogenesis and suggest novel therapeutic strategies.

**Graphical abstract:** **Age of BM-derived monocytes/macrophages determines clonality of SMC lineage in the atherosclerotic plaque.** Atherogenesis is depicted in a young (**a**) or aged (**b**) host. Aged monocytes/macrophages have decreased levels of the epigenetic regulator TET2, leading to reduction of the 5-hydroxymethylcytosine (5hmC) mark on the *Itgb3* promoter. The resulting low integrin β3 levels in aged monocytes/macrophages induces high TNF-α levels, facilitating recruitment and expansion of multiple SMC progenitors (polyclonality) in the atherosclerotic plaque and worse disease burden. In contrast, the young control is characterized by mono/oligoclonal SMC expansion in a smaller plaque.

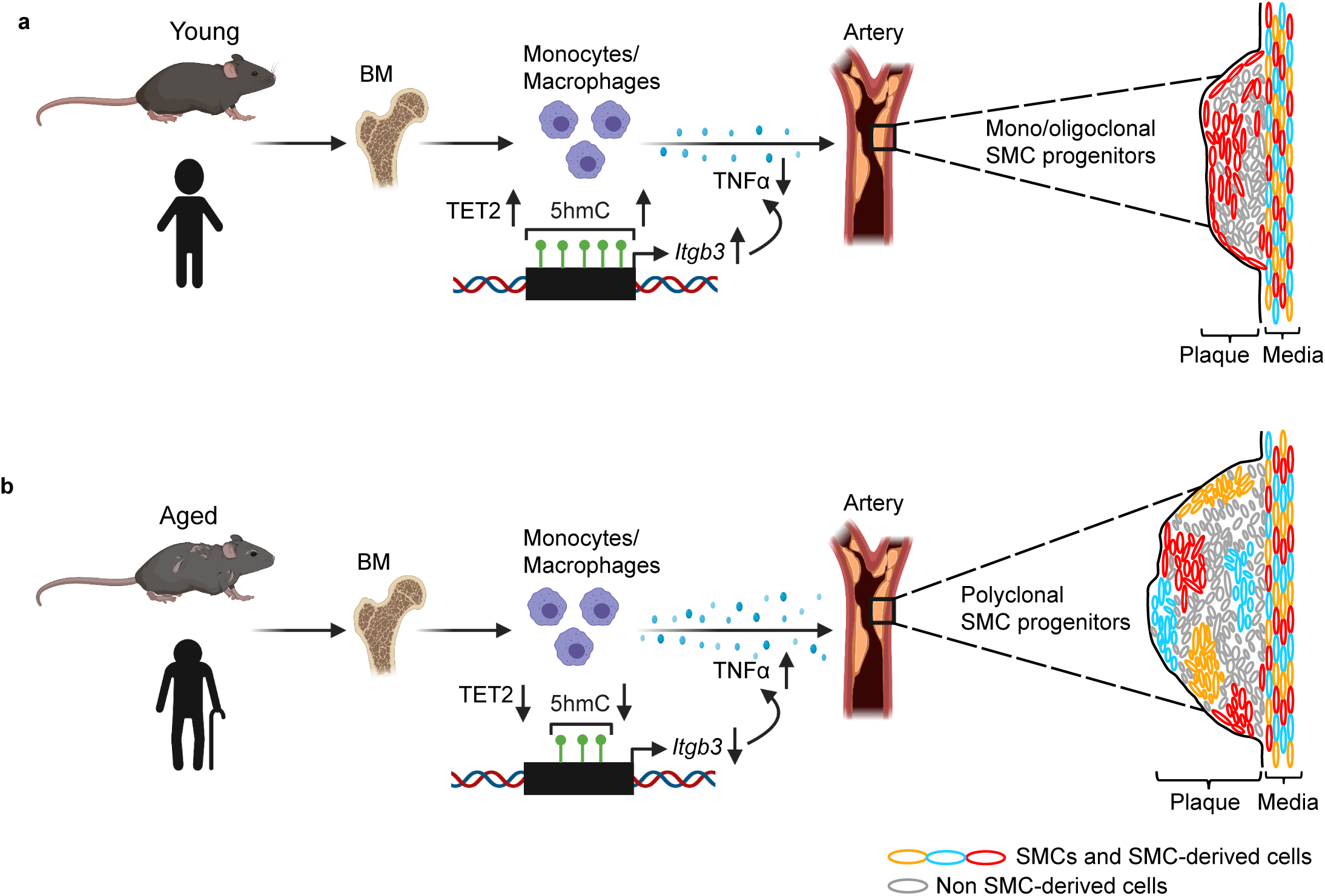

## INTRODUCTION

The revolution of single cell analysis has unveiled heterogeneity within populations of specific cell types, and clonal expansion of individual cells is increasingly appreciated to underlie a number of diseases beyond cancer, including vascular pathologies, cirrhosis, and neurodegeneration (Brunner et al., 2019; Chappell et al., 2016; Dobnikar et al., 2018; Jacobsen et al., 2017; Misra et al., 2018; Sheikh et al., 2015; Tay et al., 2017). For instance, multiple smooth muscle cell (SMC) progenitors give rise to the normal arterial wall during development, but a select few progenitors within the wall participate in atherosclerotic plaque formation (Chappell et al., 2016; Greif et al., 2012; Jacobsen et al., 2017; Misra et al., 2018). Additionally, in age-related clonal hematopoiesis of indeterminate potential (CHIP), stem cells carrying somatic mutations give rise to dominant leukocyte variants, and CHIP is associated with an increased risk of major atherosclerosis-related diseases, myocardial infarction and ischemic stroke (Jaiswal and Libby, 2020; Jaiswal et al., 2017). Recent human studies indicate a similar progressive accumulation of mutated somatic clones in epithelial cells of diverse organs with normal aging (Martincorena et al., 2018; Martincorena et al., 2015; Moore et al., 2020; Yoshida et al., 2020). Investigations have primarily focused on cell autonomous mechanisms underlying clonal expansion, and non-cell autonomous regulation is not well understood, particularly in aging. Notably, a recent controversial bioinformatic study challenges the view that increasing somatic mutations with age underlies the enhanced incidence of cancer later in life and instead argues that immune system decline is paramount (Palmer et al., 2018a, b). Herein, we investigate the phenomenon of non-cell autonomous regulation of clonality in aging by delineating the role of aged hematopoietic cells in modulating recruitment and clonal expansion of SMC progenitors in the atherosclerotic plaque.

Inflammatory cells, especially macrophages, and SMCs are key players in atherogenesis. Rare SMC progenitors from the tunica media contribute cells that populate the nascent plaque and clonally expand giving rise to ∼30-70% of the cellularity of an advanced plaque with cells forming the protective fibrous cap or transitioning into plaque destabilizing fates (Basatemur et al., 2019; Chappell et al., 2016; Jacobsen et al., 2017; Misra et al., 2018; Shankman et al., 2015; Wirka et al., 2019). Integrins are heterodimeric proteins that link the extracellular and intracellular compartments, and transplant of bone marrow (BM) null for the gene encoding integrin β3 (*Itgb3*) in atheroprone mice induces multiple SMC progenitors to enter the plaque and clonally expand, exacerbating disease burden (Misra et al., 2018; Schneider et al., 2007). On an *Apoe^(-/-)^*background, *Itgb3^(-/-)^* BM-derived macrophages have elevated levels of the cytokine tumor necrosis factor (TNF)-α (Schneider et al., 2007). Plasma TNFα levels are increased in aged mice and humans, and high TNFα levels in older humans are correlated with atherosclerosis (Bauernfeind et al., 2016; Bruunsgaard et al., 2000). Importantly, BM transplant (BMT) from aged mice into young mice worsens atherosclerosis (Du et al., 2016), but underlying mechanisms and effects on SMC recruitment and clonality are not delineated.

In CHIP, clonally expanded hematopoietic stem cells commonly harbor somatic mutations in epigenetic regulators, such as *Ten Eleven Translocation (TET)-2* (Jaiswal and Libby, 2020; Jaiswal et al., 2017). On a *Low-density lipoprotein receptor* (*Ldlr)* null background, *Tet2*-deficient BM predisposes mice to develop enhanced Western diet (WD)-induced atherosclerosis, and macrophages isolated from *Tet2^(-/-)^* mice express elevated levels of cytokines, including TNFα (Fuster et al., 2017; Jaiswal et al., 2017). Macrophages recruited to early atherosclerotic plaques proliferate locally during disease progression (Robbins et al., 2013), and in atheroprone mice, initiating WD in the aged as compared to the young increases the accumulation of macrophages in the aorta (Du et al., 2016). Importantly, in the context of CHIP in humans or in aged or myeloid cell *Tet2*-deficient mice, the clonality of myeloid cells in the atherosclerotic plaque itself is elusive. And similar to aging, the effect of BM *Tet2*-deficiency on SMC clonality in the plaque is undefined.

Herein, we report that the age of the BM is a key factor dictating the clonality of SMC-derived cells in the atherosclerotic plaque: aged BM non-cell autonomously induces recruitment and expansion of multiple SMC progenitors. Mechanistically, reduced TET2 levels in aged monocytes/macrophages epigenetically decreases *Itgb3* gene expression, which enhances TNFα-TNF receptor 1 (TNFR1) signaling. Thus, polyclonality of the SMC lineage and worse atherosclerosis ensue.

## RESULTS

### Age of mice dictates clonality of SMC- and monocyte/macrophage-derived plaque cells

In hypercholesterolemic mice, rare smooth muscle myosin heavy chain (SMMHC)^+^ cells migrate into each atherosclerotic plaque and clonally expand (Chappell et al., 2016; Jacobsen et al., 2017; Misra et al., 2018; Wang et al., 2020). However, the clonality of macrophages in plaques and how aging regulates SMC and macrophage clonality are not elucidated. (Throughout our studies, young and aged mice refer to 3 and 18 month old mice, respectively.) Young and aged mice carrying the multi-color *ROSA26R-Rainbow (Rb)* Cre reporter and either *Myh11-CreER^T2^*or *Csf1r-Mer-iCre-Mer* were induced with tamoxifen (1 mg/day for 5 or 20 days, respectively), injected with recombinant adeno-associated virus (AAV) encoding constitutively active PCSK9 ([AAV*-Pcsk9*]) and fed a WD for 16 weeks (Figure 1a). Transverse aortic root sections were stained with H&E or oil red O (ORO) or for nuclei (DAPI) and directly imaged for Rb colors (Cerulean [Cer], membrane Cherry [mCh], membrane Orange [mOr]). In comparison to young mice, plaques in aged mice are larger and have increased lipid content (Figure 1b-e). Furthermore, the percent of plaque cells that derive from SMCs (i.e., marked by any Rb color) does not change with age, but there is a marked shift towards polyclonality (Figure 1f-h). Indeed, for young mice, the most prevalent color (#1) in each plaque comprises 97±1% of marked cells, and the second and third most prevalent color (#2, 3) make up the remaining ∼3%. For aged mice, the marked cells are substantially more distributed among the Rb colors (50±5%, 28±4%, 22±6%). Importantly, the distribution of marked cells of the tunica media underlying the plaques are similar in young and aged mice (Figure 1i). In contrast to SMC tracing, in plaques of aged mice, CSF1R^+^ cells give rise to a ∼3-fold higher percentage of cells and are considerably more dominated by a single clone than in young mice (i.e., 95±3%, 3±1%, 1±1% vs. 63±9%, 25±5%, 12±3%; Figure 1j-l). Thus, aging induces polyclonality of SMC-derived cells but dominance by a single clone of the monocyte/macrophage lineage.

**Figure 1.**
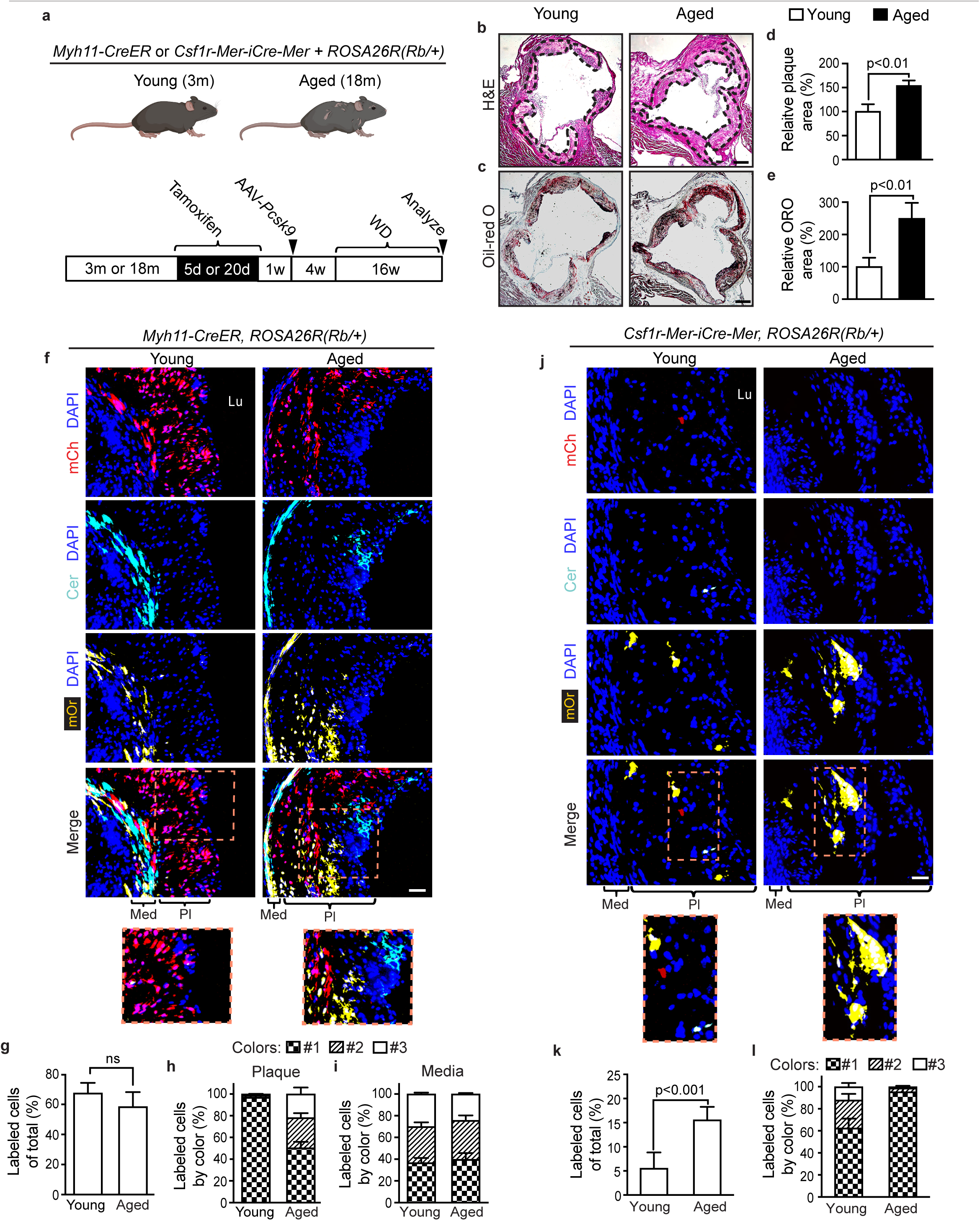
Age of mice dictates clonality of SMC- and macrophage-derived atherosclerotic plaque cells. Young (3 month) and aged (18 month) *ROSA26R^(Rb/+)^*mice also carrying either *Myh11-CreER^T2^* or *Csf1r-Mer-iCre-Mer* were induced with tamoxifen, rested, injected with AAV-*Pcsk9* and fed a Western diet (WD) for 16 weeks. Transverse aortic root sections were analyzed. **a,** Schematic of experimental plan is shown. **b-e,** For *Myh11-CreER^T2^*, *ROSA26R^(Rb/+)^* mice, sections were stained with H&E **(b;** dashed lines demarcating plaque), or with Oil Red O (ORO; **c)**, and the area of the plaque **(d)** or lipid **(e)** were quantified, respectively. n=5-7 mice per group, triplicate measurements for each mouse. **f-l,** Sections were stained for nuclei (DAPI) and directly imaged for Rb colors (mCherry [mCh], mOrange [mOr], Cerulean [Cer]), and labeled cells were quantified. In **f, j,** representative sections are shown with boxed regions displayed as close-ups below. The percent of DAPI^+^ plaque cells that were marked by any of the Rb colors was quantified in **g, k**. In **h, l**, of the marked plaque cells, the percent of cells of each color was quantified for each age group. In a given plaque, color 1 is color with the greatest number of cells in plaque (or media), color 2 is second most common color and color 3 is least frequent color. n=5-7 mice and 18 plaques per age group, 5 sections with a total of ∼1200-1600 cells and spanning 200 μm per plaque. Similarly, in **i**, of the marked cells of the underlying media, the precent of cells of each color was quantified. n=5 mice and 12 plaques analyzed per each group. Lu, lumen; Med, media; Pl, plaque. All data are mean ± SD, and Student’s *t*-test was used. Scale bars, 100 μm (**b, c**) and 50 μm **(f, j)**.

### Age of the BM is the key determinant of clonality of SMC-derived cells in the plaque

In atheroprone mice, transplant of aged or *Itgb3^(-/-)^*BM worsens WD-induced atherosclerosis, and in the latter case, results in recruitment and expansion of multiple SMCs in the plaque (Du et al., 2016; Misra et al., 2018; Schneider et al., 2007). We next queried whether aged BM is sufficient to induce polyclonality of SMC-derived plaque cells. Young *Ldlr^(-/-)^, Myh11-CreER^T2^*, *ROSA26R^(Rb/+)^* recipients were induced with tamoxifen and transplanted with BM from young and aged wild type donors (Figure 2a). Four weeks after BMT, engraftment was confirmed (Figure 2b), and then recipients were fed a WD for 16 weeks. The percentage of aortic root plaque cells derived from SMCs in recipients of aged BM is twice that in young BMT recipients (Figure 2c, d). Furthermore, labeled plaque cells in recipients of young BM are dominated by a single Rb color (95±2%) whereas in recipients of aged BM, these cells are more equally distributed (37±3%, 34±6%, 29±3%; Figure 2e). Of note, a distinct mouse model of hypercholesterolemia-induced atherosclerosis yields similar results (Figure S1a-g). Importantly, the distribution of marked media cells underlying plaques are similar in young and aged BMT groups (Figures 2f and S1h). As a complementary approach, we transplanted aged *Myh11-CreER^T2^, ROSA26R^(Rb/+)^* mice with young or aged BM and found that with young BMT, atherosclerotic lesion burden is attenuated and SMC-derived cells make up a smaller percentage of the plaque cells and are dominated by a single clone (Figure S2).

**Figure 2.**
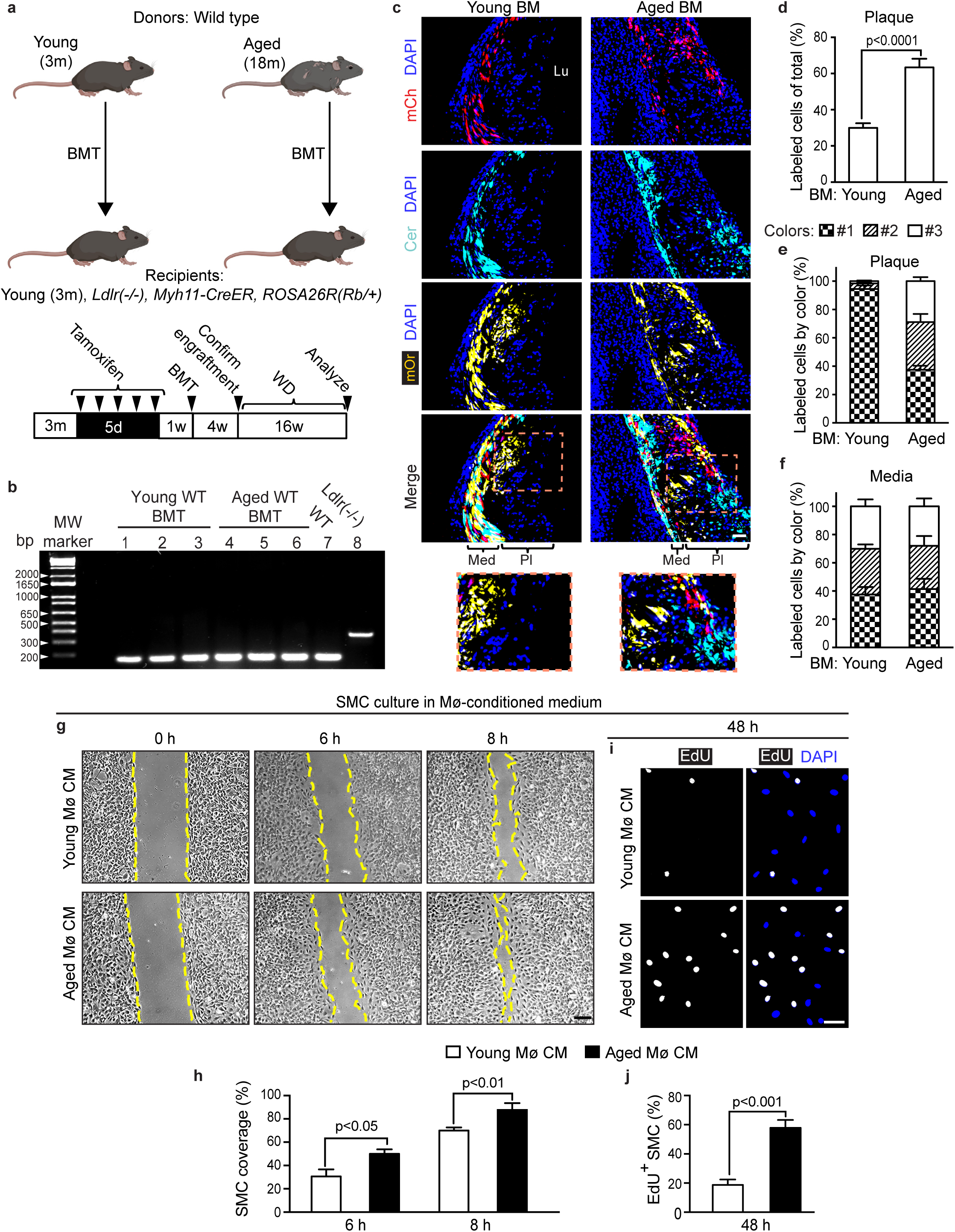
Aged bone marrow promotes expansion of multiple SMMHC^+^ progenitors in the plaque. **a-f,** Young (3 month) *Ldlr^(-/-)^*, *Myh11-CreER^T2^*, *ROSA26R^(Rb/+)^* mice were induced with tamoxifen, irradiated and then transplanted with bone marrow (BM) from young or aged (18 month) mice and fed a Western diet (WD) for 16 weeks. In **a,** experimental schematic is shown. In **b,** genomic DNA prepared from peripheral blood of *Ldlr^(-/-)^, Myh11-CreER^T2^, ROSA26R^(Rb/+)^* recipient mice following BMT from either young or aged wild type (WT) mice was PCR amplified using primers for *Ldlr*. Lanes 1-3 are from recipients with young BMT, and lanes 4-6 are from recipients with aged BMT. Lane 7 is from WT mice and lane 8 is from *Ldlr^(-/-)^* mice. In **c,** transverse aortic root sections were stained for nuclei (DAPI) and directly imaged for Rb colors (mCh, mOr, Cer) with boxed regions shown as close-ups below. In **d,** percent of DAPI^+^ plaque cells that were marked by any of the Rb colors was quantified. Of the marked plaque **(e)** or underlying medial cells **(f)**, percent of cells of each color was quantified for each BMT group. In a given plaque (or media), color 1 is color with the greatest number of cells, color 2 is second most common color and color 3 is least frequent color. n=5 mice and 15 plaques per BMT group, 5 sections with a total of ∼1000-1500 cells and spanning 200 μm per plaque. Lu, lumen; Med, media; Pl, plaque. **g-j,** BM of young and aged wild type mice was harvested and differentiated into macrophages. Macrophage-conditioned medium (CM) was added to murine aortic SMCs for migration and proliferation assays. In **g, h,** for migration assay, confluent SMCs with a central acellular area were cultured in CM for 0, 6, or 8 h as indicated. Brightfield images **(g)** and quantification of the percent **(h)** of SMC coverage at 6 or 8 h of uncovered area of 0 h are shown, respectively. n=3. In **i, j,** for proliferation assay, SMCs were incubated with CM for 48 h, and EdU was added for last 8 h. SMCs were stained for EdU and nuclei (DAPI; **i)**, and percent of cells expressing EdU was quantified **(j)**. n=3 mice. All data are averages ± SD, and Student’s *t*-test was used. Scale bars, 50 μm **(c, g)** and 25 μm **(i)**.

Macrophages regulate SMC migration and proliferation (Misra et al., 2018; Ntokou et al., 2021), which are integral for recruitment and clonal expansion, and thus, we evaluated whether aging influences this regulation. Monocytes were harvested from BM of young and aged mice and differentiated into macrophages in culture, and aortic SMCs were isolated from young mice. Incubating SMCs with conditioned medium from aged as opposed to young macrophages for 6 or 8 h induces SMC migration without altering proliferation at 8 h (Figures 2g, h and S1i, j). However, culturing SMCs for 48 h in aged macrophage conditioned medium enhances proliferation by ∼3 fold (Figure 2i, j). Taking findings from Figures 1, 2, S1 and S2 together, the age of the BM (presumably macrophages) is a decisive factor determining the recruitment and clonality of SMC-derived plaque cells.

### *Tet2* in BM cells regulates clonality of SMC-derived cells in plaques

Loss-of-function mutations in a few genes encoding epigenetic regulators, such as *TET2*, account for most CHIP cases (Jaiswal and Libby, 2020). Transplant of *Tet2*-deficient BM to *Ldlr^(-/-)^* mice exacerbates WD-induced atherosclerosis (Fuster et al., 2017; Jaiswal et al., 2017); however, the effect of BM TET2 on SMC clonality is not delineated. Thus, young *Ldlr^(-/-)^, Myh11-CreER^T2^*, *ROSA26R^(Rb/+)^* recipient mice were induced with tamoxifen, transplanted with *Tet2^(-/-)^* or wild type BM and fed a WD for 16 weeks (Figure 3a, b). The percent of plaque cells that are SMC-derived is enhanced with *Tet2^(-/-)^* BMT (Figure 3c, d). Moreover, labeled cells in plaques of the wild type BMT group are dominated by a single Rb color (88±3%, 8±3%, 4±1%) in contrast to the more even distribution in *Tet2^(-/-)^* BMT group (48±3%, 30±4%, 22±3%), without altering labeled cells in the underlying media (Figure 3e, f). Additionally, similar to studies with aged macrophages, conditioned medium from *Tet2^(-/-)^* macrophages induces SMC proliferation and migration (Figures 3g-j and S1k, l). Thus, *Tet2^(-/-)^* BM-derived cells (most likely macrophages) facilitate the recruitment and polyclonal expansion of multiple pre-existing SMCs into the plaque.

**Figure 3.**
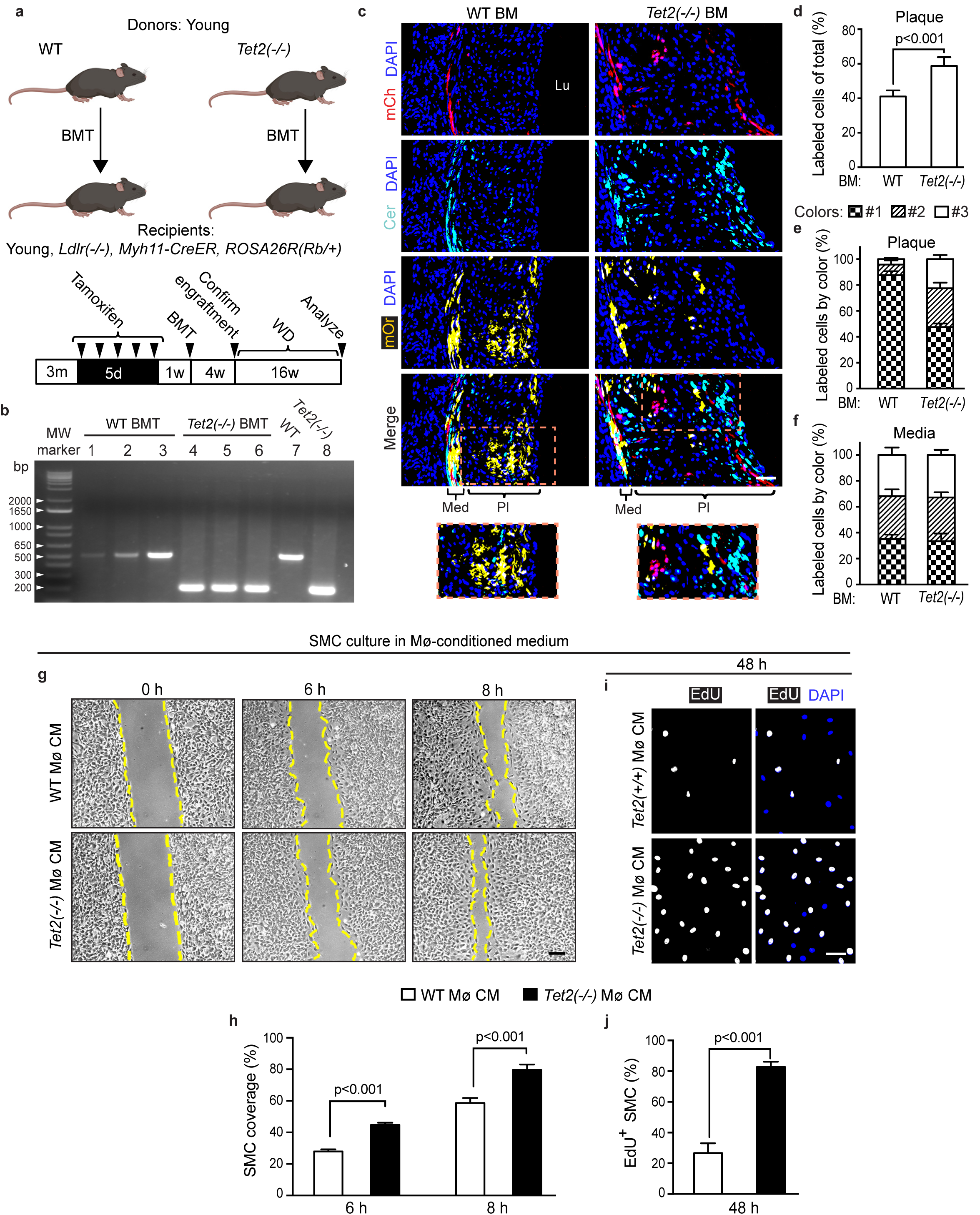
*Tet2* null BM induces expansion of multiple SMC progenitors in the plaque. **a-f,** Young (3 month) *Ldlr^(-/-)^*, *Myh11-CreER^T2^*, *ROSA26R^(Rb/+)^* mice were induced with tamoxifen, irradiated and then transplanted with BM from wild type (WT) or *Tet2^(-/-)^* mice and fed a WD for 16 weeks. In **a**, experimental schematic is shown. In **b**, genomic DNA prepared from peripheral blood of *Ldlr^(-/-)^, Myh11-CreER^T2^, ROSA26R^(Rb/+)^* recipient mice after either WT or *Tet2^(-/-)^*BM was PCR amplified using primers for *Tet2*. Lanes 1-3 are from recipient mice after WT BMT, lanes 4-6 are from recipient mice after *Tet2^(-/-)^* BMT, lane 7 is from WT mice and lane 8 is from *Tet2^(-/-)^* mice. In **c,** transverse aortic root sections were stained with DAPI and imaged for Rb colors. Boxed regions are shown as close-ups below. In **d,** percent of DAPI^+^ plaque cells that were marked by any Rb color was quantified. Of the marked plaque **(e)** or underlying medial cells **(f)**, percent of cells of each color was quantified for each BMT group. In a given plaque (or media), color 1 is color with greatest number of cells, color 2 is second most common color and color 3 is least frequent color. n=5 mice and 12 plaques per group, 5 sections with a total of ∼1000-1500 cells and spanning 200 μm per plaque. Lu, lumen; Med, media; Pl, plaque. **g-j,** BM from *Tet2*^(-/-)^ and wild type mice was harvested and differentiated into macrophages. Macrophage-conditioned medium (CM) was added to murine aortic SMCs for migration and proliferation assays. In **g, h,** for migration assay, confluent SMCs with a central area lacking cells were cultured in CM medium for 0, 6, or 8 h as indicated. Brightfield images **(g)** and quantification of percent **(h)** of SMC coverage at 6 or 8 h of uncovered area of 0 h are shown, respectively. n=3 mice. In **i, j,** for proliferation assay, murine aortic SMCs were incubated with CM for 48 h, and EdU was added for last 8 h. SMCs were stained for EdU and nuclei (DAPI; **i**), and percent of cells expressing EdU was quantified **(j)**. n=3. All data are averages ± SD, and Student’s *t*-test was used. Scale bars, 50 μm **(c, g)** and 25 μm **(i)**.

### In BM-derived monocytes, reduced TET2 is a link between aging and reduced integrin β3

To begin to evaluate how aged BM cells exacerbate atherosclerosis and induce SMC recruitment/expansion in the plaque, we investigated the effects of age on the murine and human monocyte transcriptome. CD3^-^CD19^-^Cd11b^+^Ly6C^+^ monocytes were isolated from the BM of young and aged mice by flow-activated cell sorting (FACS) and subjected to bulk RNA-sequencing (seq) (Dataset S1) and pathway analysis. Gene Set Enrichment Analysis (GSEA) indicates that inflammatory and immune pathways are upregulated in aged monocytes (Figure S3a). Ingenuity pathway analysis (IPA) shows that atherosclerosis is one of the top overrepresented and activated diseases/functions (p = 3.74 x 10^-8^, z-score=0.65). In addition, 32 differentially expressed genes significantly overlap with atherosclerosis in the Ingenuity Knowledge Base (Benjamini-Hochberg FDR p=1.28 x 10^-6^; Figure 4a). Notably, Itgb3 is downregulated in aged monocytes (Figure 4a), and in atheroprone mice, *Itgb3^(-/-)^*BMT exacerbates atherosclerosis and induces polyclonal expansion of SMC progenitors in the plaque (Misra et al., 2018; Schneider et al., 2007). However, a link between integrin β3 deficiency in monocytes/macrophages and aging has not previously been established.

**Figure 4.**
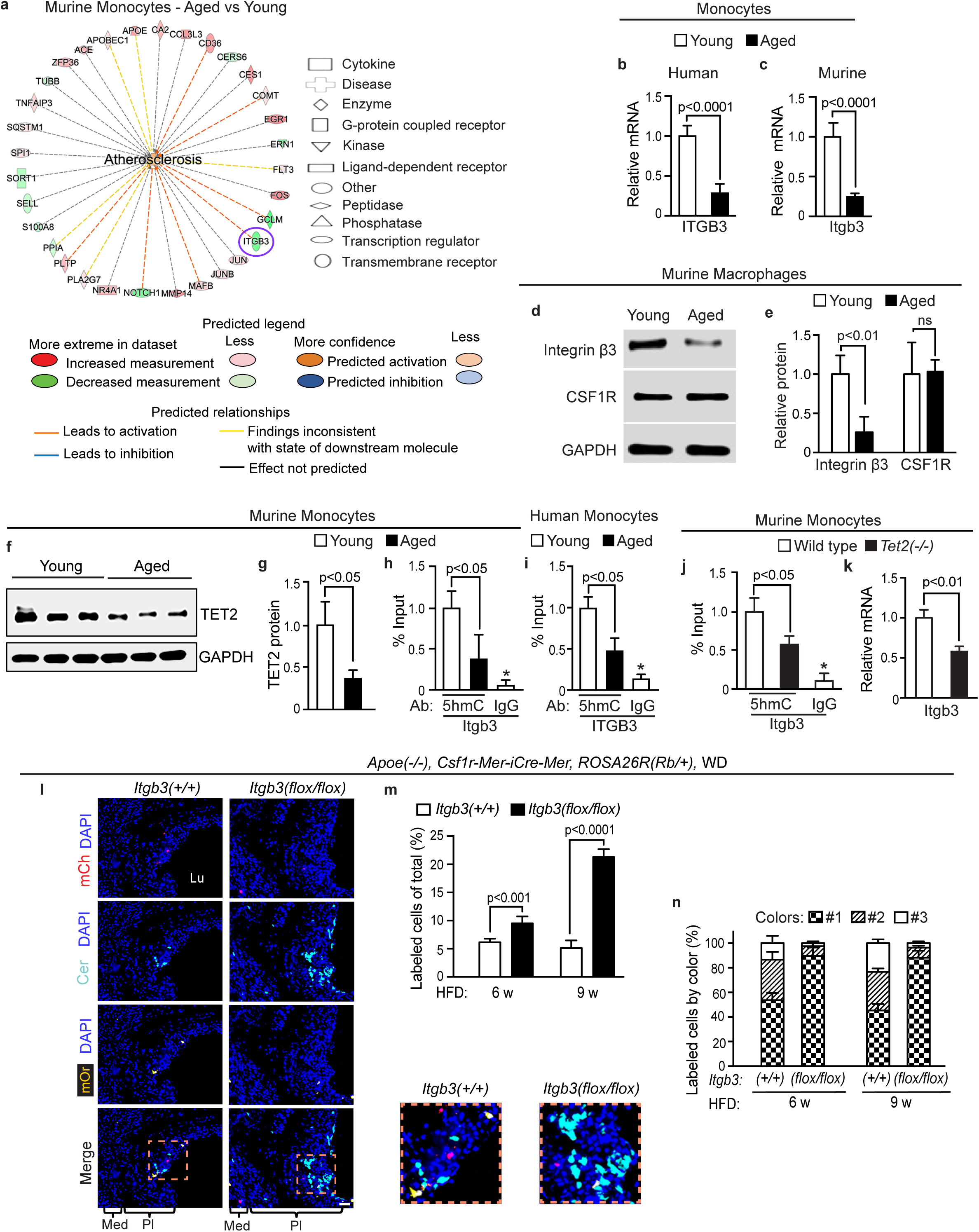
Reduced TET2 contributes to decreased integrin β3 in aged myeloid cells and *Itgb3*-deficient myeloid cells clonally expand in the plaque. **a, c-h, j, k,** BM of young (3 month) and aged (18 month) mice (**a, c-h**) or young wild type or *Tet2^(-/-)^* mice (**j, k**) was harvested and either subjected to FACS to isolate CD3^-^CD19^-^Cd11b^+^Ly6C^+^ monocytes or differentiated into macrophages. **b, i**, Peripheral blood mononuclear cells were isolated from healthy young (25±3 yr) or older (55±7 yr) humans by Ficoll density gradient centrifugation. CD3^-^CD19^-^CD14^+^ monocytes were isolated by FACS. In **a**, monocyte lysates underwent bulk RNA-seq (n=3 mice per age group) and pathway analysis. Thirty two differentially expressed genes (e.g., ITGB3 with circle) significantly overlap with atherosclerosis in the Ingenuity Knowledge Base (p=1.28 x 10^-6^). In **b, c**, qRT-PCR was used to measure monocyte ITGB3 RNA levels. In **d, e**, after macrophage differentiation, the lysates of adherent cells were subjected to Western blot for integrin β3 and myeloid marker CSF1R. Densitometry of proteins relative to GAPDH and normalized to young mice. ns, not significant. In **f, g**, monocyte lysates were subjected to Western for TET2 and GAPDH with densitometry of TET2 relative to GAPDH. In **h-j**, hydroxymethyl DNA immunoprecipitation was performed with antibodies against 5hmC or IgG, followed by qRT-PCR with primers targeting upstream of the *ITGB3* transcription start site (chr11:104,608,100-104,608,250 in mice; chr17:47,240,300-47,240,400 in human). *\*p<*0.0001 vs. 5hmC, young (**h, i**) and wild type (**j**). In **k**, Itgb3 mRNA levels were measured in monocytes from wild type or *Tet2^(-/-)^* mice. n=12 humans and n=3-5 mice per age or genotype group. **l-n**, *Apoe^(-/-)^*, *Csf1r-Mer-iCre-Mer, ROSA26R^(Rb/+)^* mice also carrying *Itgb3^(flox/flox)^* or wild type for *Itgb3* were induced with tamoxifen, rested, fed a WD for 6 or 9 weeks (w). Transverse aortic root sections were stained with DAPI and directly imaged for mCh, mOr, Cer with boxed regions shown as close-ups on the right (9 weeks WD in **l**). In **m**, percent of DAPI^+^ plaque cells that are also marked by any Rb color is shown. In **n**, of the marked plaque cells, the percent of cells of each color was quantified per age and genotype group. In a given plaque, color 1 has the greatest number of cells, color 2 is second most common and color 3 is least frequent. n=5-7 mice and 14 plaques per group, 5 sections with a total of ∼1200-1400 cells and spanning 200 μm per plaque. Data are mean ± SD. Student’s *t*-test (**b, c, e, g, k, m**) and multifactor ANOVA with Tukey’s post hoc test (**h-j**) were used. Scale bar, 50 μm (**l**).

We further explored this potential link by directly evaluating specific gene product levels in monocytes/macrophages from mice and humans. BM was harvested from young and aged mice, and we either isolated monocytes by FACS or differentiated them into macrophages in culture. Additionally, FACS was used to isolate peripheral blood CD3^-^CD19^-^CD14^+^ monocytes from healthy young and older humans (25±3 yr and 55±7 yr, respectively; Supplemental Table 1). qRT-PCR of monocyte lysates demonstrate substantially reduced ITGB3 mRNA levels in aged humans and mice (Figure 4b, c). Furthermore, in comparison to young macrophages, aged macrophages have a 75±19% reduction in integrin β3 protein levels but similar levels of the monocyte/macrophage marker CSF1R (Figure 4d, e).

We then queried whether Itgb3 levels are modulated by TET2 which catalyzes the oxidation of 5-methylcytosine (5mC) to 5-hydroxymethylcytosine (5hmC), a key step during DNA demethylation and thus, gene expression. Monocytes from aged mice have reduced TET2 protein levels, and 5hmC levels in proximity to the *ITGB3* transcription start site in aged human and murine monocytes are decreased (Figure 4f-i). Similarly, monocytes isolated from *Tet2^(-/-)^* mice have reduced 5hmC levels at the *Itgb3* proximal promoter and reduced Itgb3 transcript levels (Figure 4j, k). Hence, the data suggest that reduced TET2 in aged monocytes leads to reduced ITGB3 levels by attenuating transcription.

### *Itgb3*-deficient monocytes/macrophages exacerbate atherosclerosis and clonally expand in the plaque

Although transplant of atheroprone mice with *Itgb3^(-/-)^*BM worsens atherosclerosis (Misra et al., 2018; Schneider et al., 2007), the *in vivo* role of integrin β3 in myeloid cells has not been elucidated. *Apoe^(-/-)^, Csf1r-Mer-iCre-Mer* mice carrying *Itgb3^(flox/flox)^* or *Itgb3^(+/+)^*were induced with tamoxifen and fed a WD for 16 weeks. Deletion of *Itgb3* in CSF1R^+^ cells exacerbates aortic root plaque burden, lipid accumulation and acellular necrotic core area (Figure S3b-g).

We next studied mice of the same genotypes and additionally with *ROSA26R^(mTmG/+)^* or *ROSA26R^(Rb/+)^*. Five days after tamoxifen induction of *Apoe^(-/-)^, Csf1r-Mer-iCre-Mer, ROSA26R^(mTmG/+)^* mice also carrying *Itgb3^(flox/flox)^*or *Itgb3^(+/+)^*, cells expressing GFP and the monocyte marker Ly6C were isolated from BM by FACS and peripheral blood was collected as well. qRT-PCR analysis of Ly6C^+^GFP^+^ BM cells demonstrated a 94±4% reduction in Itgb3 levels in mice carrying *Itgb3^(flox/flox)^*(Figure S3h). Deletion of *Itgb3* in CSF1R^+^ cells does not alter circulating levels of total cholesterol, triglycerides or leukocyte sub-types (Figure S3i-k). To evaluate the fate and proliferation of the lineage derived from CSF1R^+^ cells, following tamoxifen induction, mice of these same genotypes were fed a WD for 16 weeks and then injected with EdU intraperitoneally 12 h prior to euthanasia. *Csf1r-Mer-iCre-Mer-*mediated deletion of *Itgb3* leads to proliferation and accumulation of cells of this lineage in the plaque as well as an increased percentage of total plaque cells that express the macrophage marker CD68 (Figure S3l-p). Given that CSF1R^+^ cells clonally expand in the plaques of aged mice (see Figure 1j-l) and the link between aging and monocyte/macrophage Itgb3 deficiency (see Figure 4a-i), we next queried whether CSF1R^+^ cells deficient in *Itgb3* clonally expand. *Apoe^(-/-)^, Csf1r-Mer-iCre-Mer, ROSA26R^(Rb/+)^*mice also harboring *Itgb3^(flox/flox)^* or *Itgb3^(+/+)^*were induced with tamoxifen and then fed a WD for 6 or 9 weeks. For aortic root plaques in mice carrying *Itgb3^(flox/flox)^*, a higher percentage of cells derive from the CSF1R^+^ lineage and a single Rb color gives rise to ∼90% of marked cells at each WD duration, whereas in mice wild type for *Itgb3*, there is a more equitable distribution of the Rb colors (Figure 4l-n). Taken together, integrin β3 deficiency in monocytes/macrophages leads to clonal expansion of this population in the atherosclerotic plaque and more severe disease.

### *Itgb3*-deficient SMCs contribute less to the plaque and attenuate atherosclerosis

In addition to myeloid cells, SMCs express high levels of integrin β3, and on an atheroprone background, global *Itgb3* nulls have worse atherosclerosis (Misra et al., 2018); however, in this context, the cell autonomous role of integrin β3 in SMC-derived cells is not defined. Thus, we used *Myh11-CreER^T2^* to delete *Itgb3* in SMCs, which attenuates atherosclerosis severity and proliferation and accumulation of SMC-derived cells without altering clonality of this lineage in the plaque (Figure S4). To evaluate mechanisms that may link integrin β3 deficiency in SMCs to diminished atherogenesis, we studied signaling pathways implicated in inducing SMC proliferation and migration (Fernández-Hernando et al., 2009; Gerthoffer, 2007; Nelson et al., 1998; Stabile et al., 2003). Western blots indicate a 72±7% reduction in phosphorylated AKT in aortic homogenates from *Itgb3^(-/-)^* mice (Figure S5a, b). Alternatively, siRNA targeting of Itgb3 in isolated wild type aortic SMCs significantly reduces protein levels of phosphorylated AKT, extracellular regulated kinase and p21 activated kinase-1 (Figure S5c, d). Moreover, Itgb3 silencing abrogates platelet-derived growth factor-B-induced dorsal ruffle formation (Figure S5e, f), an early step in cell migration (Gu et al., 2011). In sum, integrin β3-deficient SMCs have decreased proliferation and migration, which likely accounts for their reduced contribution to the atherosclerotic plaque.

### *Itgb3^(-/-)^* monocytes/macrophages are pro-inflammatory

In contrast to SMCs, integrin β3 deficiency in macrophages non-cell autonomously enhances SMC migration and proliferation, and transplant of *Itgb3^(-/-)^* BM induces SMC polyclonal expansion in the atherosclerotic plaque (Misra et al., 2018) (see Figures S3-S5). To further interrogate the role of integrin β3 in monocytes/macrophages, we performed single cell (sc) RNA-seq in BM cells from *Apoe^(-/-)^* mice that are also wild type or null for *Itgb3*. Unsupervised clustering was used to generate t-distributed stochastic neighbor embedding (t-SNE) plots, separating the cells into four major cell type groups: neutrophils, monocytes/macrophages, B-cells, T-cells (Figures 5a, b and S6a). We evaluated the expression pattern of classical markers of monocytes/macrophages (Figure S6b-h), and GSEA of these cells on an *Apoe^(-/-)^* background demonstrates that *Itgb3* deletion leads to activation of inflammation, including cytokine signaling pathways (e.g., nuclear factor-kappa B [NF-kB], TNF; Figure 5c-e). In addition, IPA predicts that *Itgb3* deletion induces the inflammatory response and monocyte/macrophage migration and proliferation (Figure S6i). Moreover, sub-cluster analysis of monocytes/macrophages demonstrates diverse subpopulations in both genotypes, with an increased abundance of transcriptionally distinct M1-like and M2-like macrophage populations in the *Itgb3* null genotype (Figures 5f, g and S6j). Furthermore, TNF and NF-kB signaling pathways are enriched in *Itgb3* null subpopulations, most dramatically in M1-like macrophages, compared to *Itgb3^(+/+)^* control (Figure 5h, i and S6k, l).

**Figure 5.**
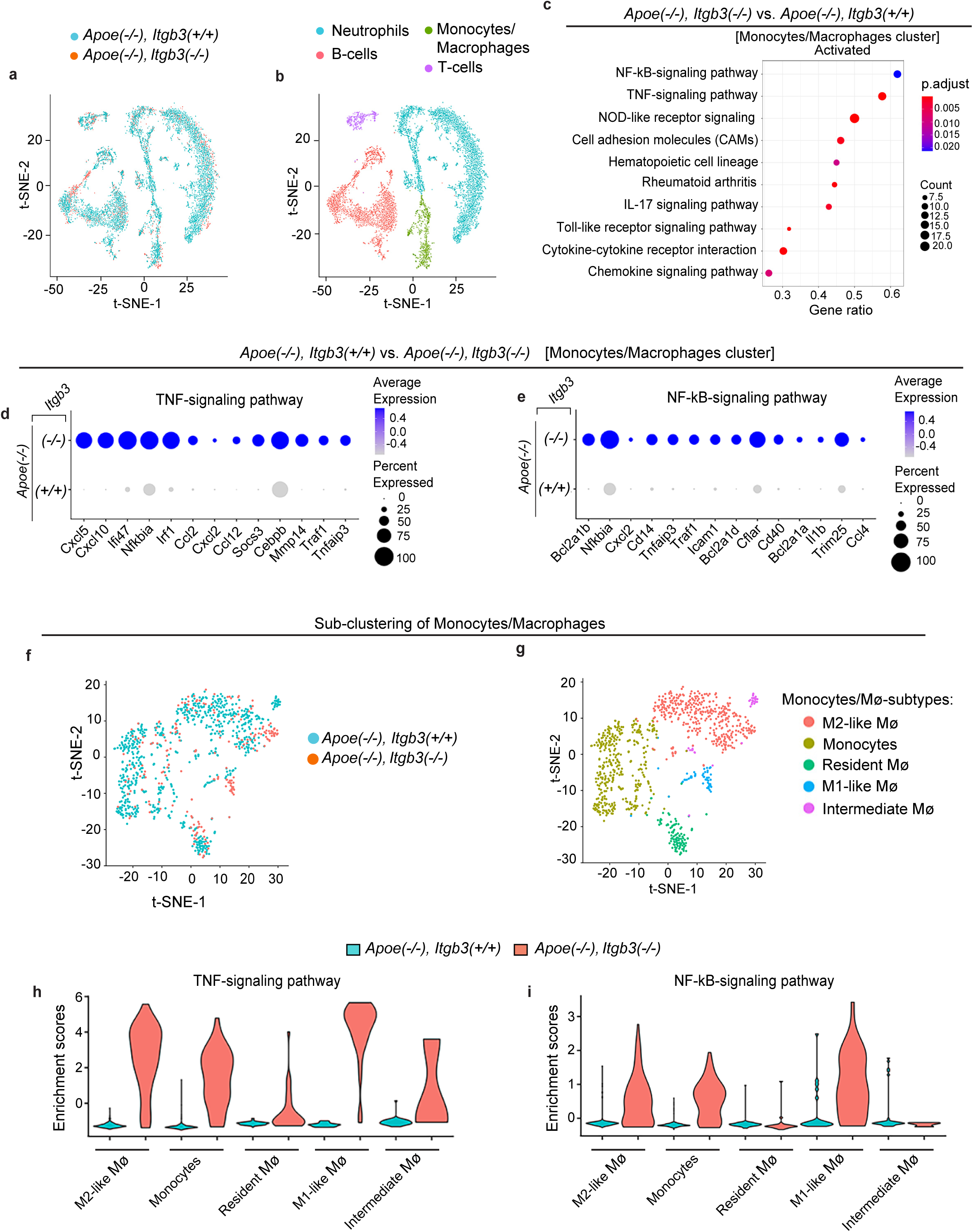
*Itgb3^(-/-)^* monocytes/macrophages induce inflammatory pathways. scRNA-seq was conducted on BM cells isolated from *Apoe^(-/-)^* mice that are also either wild type or null for *Itgb3.* **a,** Overlayed t-SNE plots of scRNA-seq of BM cells from mice carrying *Itgb3^(+/+)^* (blue) or *Itgb3^(-/-)^* (red) are shown. **b**, t-SNE plot combining cells of both genotypes with distinct clusters that are identified as major cell types based on expression levels of cluster markers (see Supplemental Table 4). **c-e,** Pathway and gene expression analysis of single cell transcriptomes of the monocyte/macrophage cluster. In **c,** GSEA identifies activated pathways. Gene ratio indicates the number of activated genes relative to the total number of genes in the corresponding pathway, and dot size represents the number of dysregulated genes in each pathway. In **d, e,** dot plot analyses of differentially expressed genes in TNFα and NF-kB signaling pathways. Dot size and blue color represent the fraction of cells expressing the gene and the average gene expression, respectively. **e**, IPA predicts the effects of differentially expressed genes on the inflammatory responses and monocyte/macrophage migration or proliferation. n=3 mice pooled per genotype. **f, g,** t-SNE plots sub-clustering monocytes/macrophages from mice carrying *Itgb3^(+/+)^*or *Itgb3^(-/-)^* are shown. In **f**, cells are colored by genotype. In **g,** plot combining both genotypes with distinct sub-clusters identified (see Supplemental Table 5). **h, i,** Enrichment scores, defined by differentially expressed genes in TNFα and NF-kB signaling pathways, were generated by using “Add Module Score” function in the Seurat package. n=3 mice pooled per genotype.

### TNFα regulates clonal expansion of SMC-derived plaque cells

Treatment with an anti-TNFα antibody prolongs survival of WD fed *Apoe^(-/-)^, Itgb3^(-/-)^* mice (Schneider et al., 2007); however, effects of TNFα inhibition on atherosclerotic burden are controversial (Branen et al., 2004; Oberoi et al., 2018), and effects on SMC clonality are not defined. To this end, *Apoe^(-/-)^, Csf1r-Mer-iCre-Mer, Itgb3^(flox/flox)^* mice were induced with tamoxifen and injected intraperitoneally with an anti-TNFα neutralizing antibody or IgG2a isotype control twice per week during 12 weeks of WD feeding (Figure 6a). TNFα inhibition attenuates plaque area and lipid burden in the aortic root (Figure 6b-e). For clonal analysis, *Apoe^(-/-)^, Myh11-CreER^T2^, ROSA26R^(Rb/+)^* mice were induced with tamoxifen, transplanted with *Apoe^(-/-)^* BM that was also wild type or null for *Itgb3* and started on a 12 week regimen of WD with concomitant twice weekly injections of anti-TNFα antibody or control IgG2a (Figure 6f-h). In both *Itgb3* wild type or *Itgb3^(-/-)^* BMT groups, inhibition of TNFα reduces SMC-derived plaque cells by more than 50% (Figure 6i-k). Furthermore, in mice transplanted with the *Itgb3^(-/-)^* BM, anti-TNFα treatment induces predominance of a single SMC-derived clone (87±6%, 10±3%, 3±2%) in comparison to the more evenly distributed polyclonality with IgG2a control (40±4%, 33±2%, 26±3%) (Figure 6l). Anti-TNFα treatment does not alter clonal distribution of SMC-derived cells in either plaques of the *Itgb3^(+/+)^*BMT group or the underlying media with each BMT group (Figure 6l, m).

**Figure 6.**
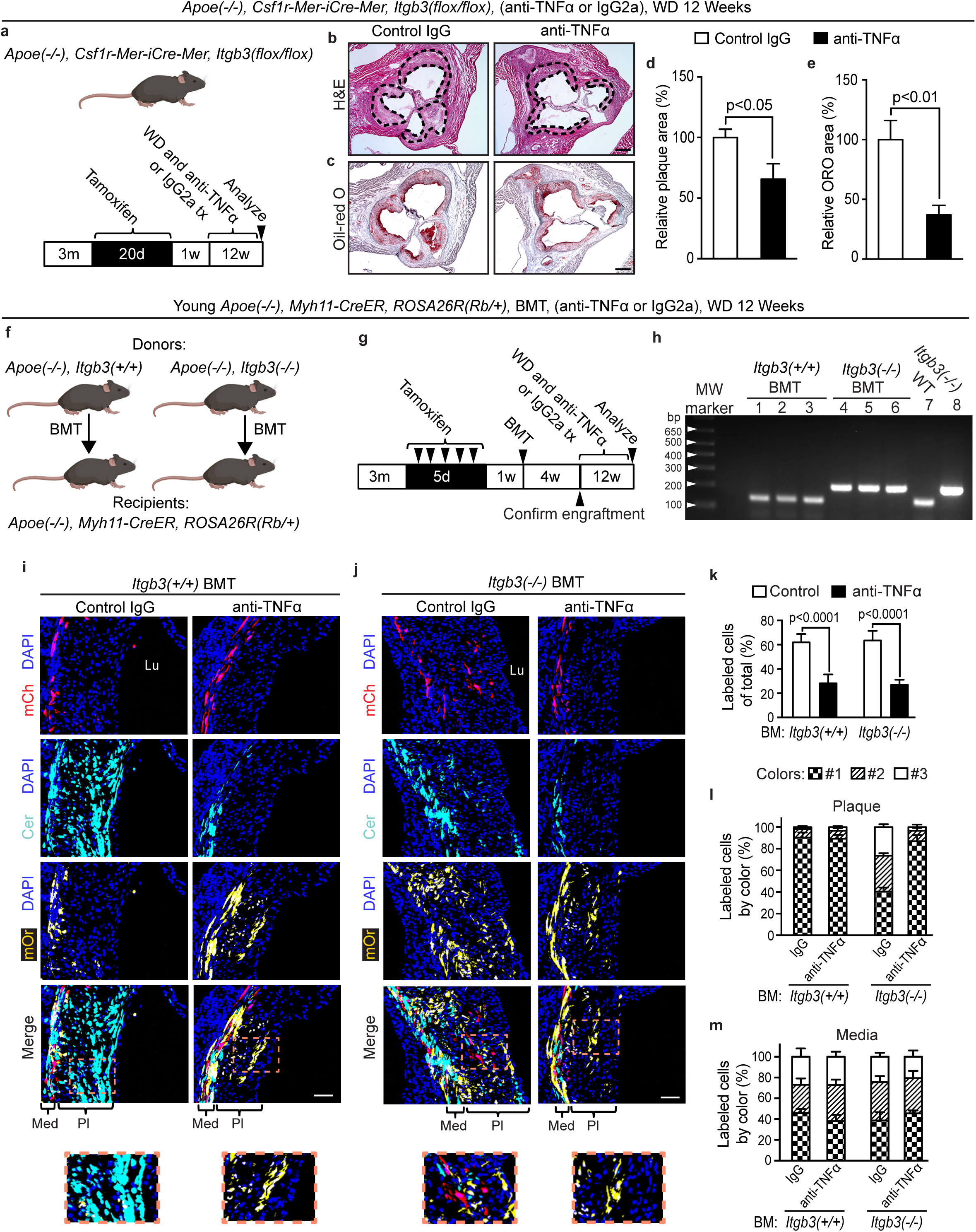
In the context of *Itgb3^(-/-)^* BM, TNFα antagonism preserves predominance of a single SMC clone in plaques. Mice were treated with 12 weeks of WD with concomitant two times per week injections of isotype control IgG2a or anti-TNFα antibody (20 mg/kg) and then transverse aortic root sections were analyzed. **a-e,** Prior to these treatments, *Apoe^(-/-)^*, *Csf1r-Mer-iCre-Mer* mice also carrying *Itgb3^(flox/flox)^* or wild type for *Itgb3* were induced with tamoxifen. In **a,** experimental schematic is shown. Sections were stained with H&E (dashed lines demarcate lesion; **b)** and Oil Red O **(c)**, and lesion area **(d)** and lipid content **(e)** were quantified. **f-m**, Prior to the WD and treatment with control or anti-TNFα antibody, *Apoe^(-/-)^*, *Myh11-CreER^T2^*, *ROSA26R^(Rb/+)^* mice were induced with tamoxifen, irradiated and then transplanted with *Apoe^(-/-)^* BM also carrying *Itgb3^(+/+)^*or *Itgb3^(-/-)^* as indicated. In **f, g,** schematic is shown. In **h,** genomic DNA prepared from peripheral blood of *Apoe^(-/-)^, Myh11-CreER^T2^, ROSA26R^(Rb/+)^* recipient mice with *Apoe^(-/-)^, Itgb3^(+/+)^* or *Apoe^(-/-)^, Itgb3^(-/-)^* BM or of WT or *Itgb3^(-/-)^*mice was amplified using primers for *Itgb3.* In **i, j,** sections were stained with DAPI and directly imaged for Rb colors. In **k,** percent of DAPI^+^ plaque cells that were marked by any Rb color was quantified. Of the marked plaque **(l)** or underlying medial cells **(m)**, the percent of cells of each color was quantified for each BMT and treatment group. In a given plaque (or media), color 1 is the most dominant color of cells, color 2 is the second most prevalent and color 3 is least frequent. n=5 mice and 12 plaques per group, 5-6 sections with a total of ∼1150-1650 cells and spanning 200 μm per plaque. All data are averages ± SD, and Student’s *t*-test was used. Lu, lumen; Med, media; Pl, plaque. Scale bars, 200 μm **(b, c)** and 50 μm **(i, j)**.

In addition, as aged myeloid cells have reduced integrin β3 (see Figure 4a-e) and increased Tnfa transcript levels (Figure S7a), we similarly assessed the role of TNFα in regulating SMC clonality of the atherosclerotic plaque following aged BMT. Importantly, aged BMT increases plasma TNFα levels by ∼4.5-fold compared to young BMT (Figure S7b). Young *Apoe^(-/-)^, Myh11-CreER^T2^, ROSA26R^(Rb/+)^* mice were induced with tamoxifen, transplanted with BM from aged wild type mice and then fed a WD for 12 weeks with concomitant twice weekly anti-TNFα antibody or control IgG2a injections (Figure S7c-e). Following aged BMT, TNFα inhibition attenuates plaque burden and lipid deposition (Figure 7a-d). Moreover, in this context, anti-TNFα treatment reduces the contribution of SMC-derived cells to the plaque and promotes expansion of a single SMC-derived clone in comparison to IgG2a treatment without altering clonality in the media (Figure 7e-h). In addition, incubating aortic SMCs from a young mouse with anti-TNFα antibody-treated conditioned medium from aged macrophages attenuates SMC proliferation at 48 h and migration at 8 h (but not proliferation at 8 h) in comparison to exposure to IgG2a-treated conditioned medium (Figures 7i, k and S7f-i). Similarly, RNA silencing of Tnfr1 in isolated murine aortic SMCs attenuates aged macrophage conditioned medium-induced SMC proliferation. (Figure 7j, k and S7j, k).

**Figure 7.**
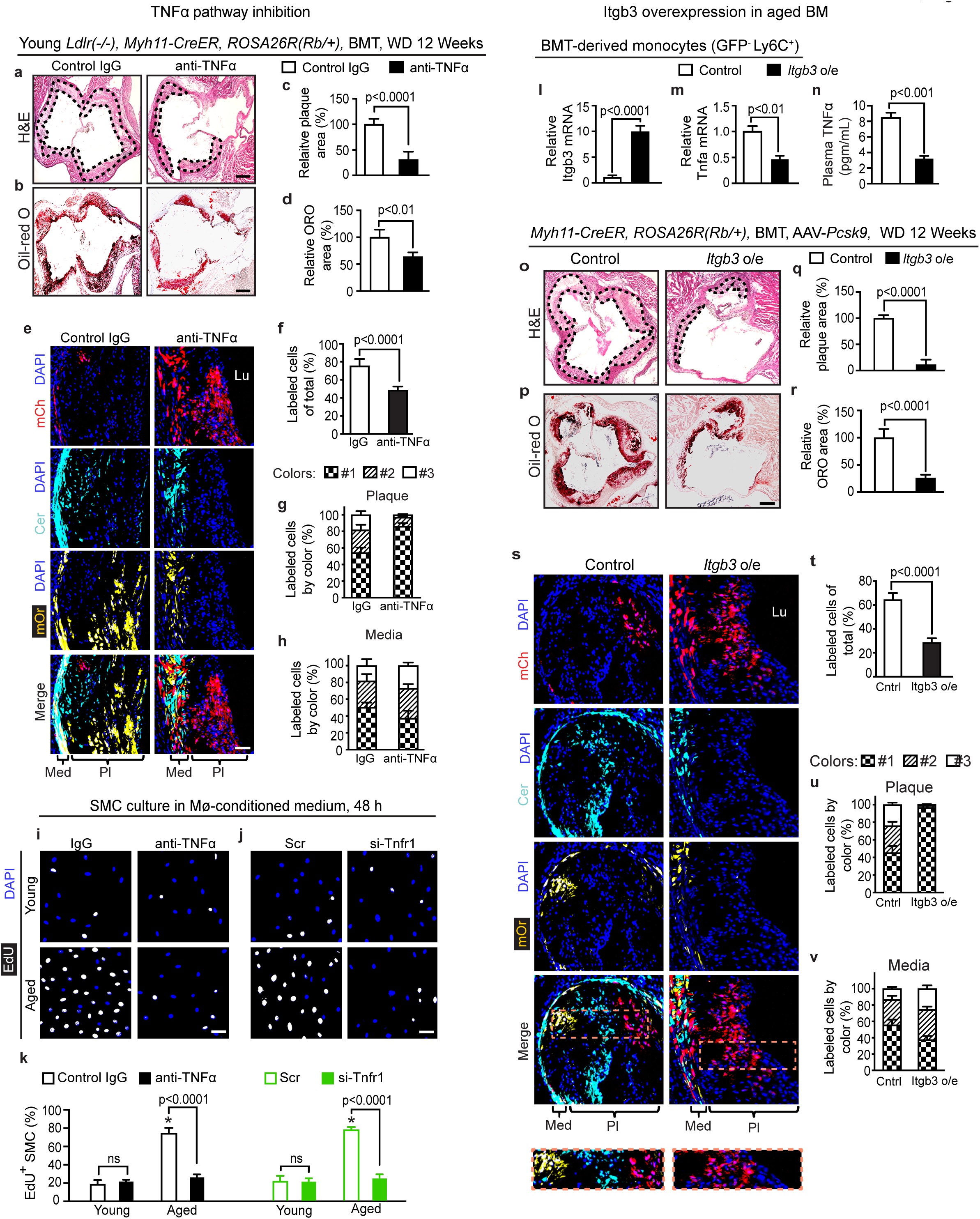
Aged BM induces SMC polyclonal expansion, proliferation and atherosclerosis via the TNFα-TNFR1 axis and BM *Itgb3* overexpression attenuates these effects. **a-h,** Young *Ldlr^(-/-)^, Myh11-CreER^T2^, ROSA26R^(Rb/+)^* recipient mice were transplanted with aged BM and then treated with 12 weeks of WD and concomitant two times per week injections of anti-TNFα antibody (20 mg/kg) or isotype control IgG2a. Transverse aortic root sections were analyzed. Sections were stained with H&E (dashed lines demarcate lesion; **a)** and Oil Red O (**b**), and lesion area (**c**) and lipid content (**d**) were quantified. n=5 mice per BMT group, 8-10 plaques and triplicate from each plaque were analyzed. In **e**, sections were stained with DAPI and directly imaged for Rb colors. The percent of DAPI^+^ plaque cells marked by any Rb color was quantified (**f**). Of the marked plaque (**g**) or underlying medial cells (**h**), the percent of cells of each color was quantified for control IgG and anti-TNFα treatment groups. Colors #1-3 are defined as in previous figures. n=5 mice and 10 plaques per group, 5-6 sections with a total of ∼1245-1400 cells and spanning 200 μm per plaque. **i-k**, BM of young or aged wild type mice was harvested and differentiated into macrophages, and isolated mouse aortic SMCs were cultured with macrophage-conditioned medium for 48 h with EdU included for the last 8 h. In **i**, anti-TNFα blocking antibody or control IgG was added to macrophage-conditioned medium (CM), 1 h prior to incubation with SMCs. In **j**, isolated SMCs were isolated and treated with Tnfr1 siRNA or scrambled (Scr) RNA prior to CM exposure. SMCs were stained for EdU and nuclei (DAPI; **i, j**), and percent of EdU^+^ cells was quantified (**k**). n=3 mice. * vs. young, p<0.0001. **l-v**, BM cells harvested from aged wild type mice were infected with control or *Itgb3* overexpressing (o/e) lentivirus. After 1 day of infection, aged *Myh11-CreER^T2^*, *ROSA26R^(Rb/+)^* mice were induced with tamoxifen, irradiated and then transplanted with control or *Itgb3* o/e lentivirus-treated BM cells. In **l-n**, 4 weeks after BMT, Itgb3 and Tnfa transcript levels were measured by qRT-PCR in GFP^-^ Ly6C^+^ BM-derived monocytes and plasma TNFα levels were measured by ELISA. In **o-v**, 4 weeks after BMT, mice were injected with AAV-*Pcsk9* and then fed a WD for 12 weeks, and transverse aortic root sections were analyzed. Sections were stained with H&E (dashed lines demarcate lesion; **o**) and Oil Red O (**p**), and lesion area (**q**) and lipid content (**r**) were quantified. n=4 mice per BMT group, 10-12 plaques and triplicate from each plaque. Sections were stained with DAPI and directly imaged for Rb colors with boxed regions displayed as close-ups below (**s**). The percent of DAPI^+^ plaque cells marked by any Rb color was quantified (**t**). Of the marked plaque (**u**) or underlying medial cells (**v**), the percent of cells of each color was quantified for control and *Itgb3* o/e groups. n=4 mice and 12 plaques per group, 5-6 sections with a total of ∼1275-1350 cells and spanning 200 μm per plaque. Lu, lumen; Med, media; Pl, plaque. All data are averages ± SD. Multifactor Student’s *t*-test (**c, d, f-h, l-n, q, r, t-v**) and ANOVA with Tukey’s post hoc test (**k**) were used. Scale bars, 100 µm (**a, b, o, p**), 50 μm (**e, s**), 25 μm (**i, j**).

Finally, given the potential central role of reduced monocyte/macrophage Itgb3 in age-induced increased TNFα and thus, polyclonal SMC progenitor expansion in the plaque, we evaluated the effect of overexpressing *Itgb3* in BM cells in aged mice. To this end, BM harvested from aged wild type mice was infected with *Itgb3*-expressing or control lentivirus and transplanted into aged *Myh11-CreER^T2^, ROSA26R^(Rb/+)^* mice (Figure S7l-n). These mice were then induced with AAV-*Pcsk9* and fed a WD (Figure S7m). *Itgb3* overexpression in BM-derived monocytes markedly reduces Tnfa mRNA and plasma TNFα, but not lipid, levels at 4 weeks after BMT (Figures 7l-n and S7o, p). Furthermore, enhanced *Itgb3* expression in aged BM cells reduces plaque burden and leads to dominance of a single SMC-derived clone in the plaque in comparison to control (Figure 7o-v). Taken together, these studies implicate in aged macrophages, an inductive effect of integrin β3 deficiency on TNFα which signals to SMC progenitors, likely through TNFR1, to induce polyclonal expansion and worse atherosclerosis.

## DISCUSSION

The main finding of this work is that the age of the BM non-cell autonomously dictates clonality of SMCs in the atherosclerotic plaque, regardless of the age of SMCs or other non-hematopoietic cells (Figures 1, 2, S1 and S2). Up to two-thirds of the cellularity of an advanced atherosclerotic plaque in a young mouse derives from clonal expansion of one or two SMC progenitors (Basatemur et al., 2019; Chappell et al., 2016; Jacobsen et al., 2017; Misra et al., 2018; Shankman et al., 2015; Wang et al., 2020). Herein, we find that in plaques of aged mice, SMC expansion is instead polyclonal as a result of aged BM-derived cells, most likely macrophages (Figures 1, 2). Similar to atherogenesis, neointima formation in ligated carotid arteries of young mice is characterized by oligoclonal expansion of rare SMCs (Chappell et al., 2016), and interestingly, SMC senescence does not alter clonality in this model (Uryga et al., 2021).

In regard to clonality of leukocytes, pivotal incompletely understood CHIP-related questions include: i) do leukocyte clones accumulate in atherosclerotic plaques? ii) mutations in epigenetic modulators induce expression changes in which downstream genes? and iii) how do these mutations exacerbate atherosclerosis? Our results demonstrate that TET2 and integrin β3 are reduced in monocytes/macrophages of aged mice and/or humans, and aged or *Itgb3*-deficient monocytes/macrophages clonally expand in plaques (Figures 1, 4). *Tet2* null BM cells non-cell autonomously induce polyclonal expansion of SMCs in the plaque, and conditioned medium from *Tet2^(-/-)^* macrophages induces SMC migration and proliferation (Figure 3). A regulatory role for *ITGB3* promoter methylation has not been previously reported, but herein, we find decreased 5hmC on the *ITGB3* promoter with aging or *Tet2* deletion (Figure 4). These effects of aging are cell type-specific as ITGB3 levels are in fact increased in aged murine liver or human fibroblasts (Rapisarda et al., 2017).

In humans, a polymorphism of the *ITGB3* gene (T1565C) is associated with coronary artery disease (Bogatyreva et al., 2018). Mice with global *Itgb3* deletion or with *Itgb3* null BM have polyclonal SMC expansion in the atherosclerotic plaque and worse disease (Misra et al., 2018; Schneider et al., 2007; Weng et al., 2003). Deletion of *Itgb3* in monocytes/macrophages mimics results with *Itgb3^(-/-)^* BMT in terms of worsening atherosclerotic burden (Figure S3). In contrast, SMC deletion of *Itgb3* attenuates disease and SMC contribution to plaques without altering clonality (Figure S4). Interestingly, aged arteries have increased levels of the anti-efferocytic protein CD47, which was initially co-purified with integrin β3 and regulates ligand binding of integrin αvβ3 (Brown et al., 1990; Ghimire et al., 2020; Lindberg et al., 1993). Anti-CD47 treatment attenuates atherosclerosis and induces a more even distribution of colors of Rb-marked SMC-derived cells (i.e., polyclonal expansion) (Wang et al., 2020). Future studies of cell type-specific roles of CD47, its interplay with integrin β3 and downstream signaling in the context of atherosclerosis and aging are warranted.

The most significantly upregulated functional class of genes in *Tet2* null BM-derived macrophages consists of cytokines, chemokines and their receptors (Jaiswal et al., 2017), and herein, scRNA-seq of *Itgb3* null BM-derived monocytes/macrophages demonstrates enhanced chemokine and cytokine signaling and in particular TNF signaling (Figure 5). Plasma TNFα increases with aging, and studies suggest that the TNFα pathway positively regulates atherosclerosis and myocardial infarction (Bauernfeind et al., 2016; Bruunsgaard et al., 2000; Ridker et al., 2000). Two SNPs of the *TNF receptor-1* (*TNFR1*) gene are associated with coronary artery disease in aged adults (Zhang et al., 2010). Furthermore, carotid arteries from aged and young *Tnfr1^(-/-)^*mice were grafted into *Apoe^(-/-)^* mice, and most interestingly, arteries from aged *Tnfr1^(-/-)^*mice are protected from age-induced atherosclerosis (Zhang et al., 2010). We now report that pharmacological inhibition of TNFα attenuates atherosclerosis, limiting the recruitment / clonal expansion of SMCs with *Itgb3^(-/-^*^)^ or aged BM and that Tnfr1 knockdown in SMCs or TNFα inhibition reduces proliferation and/or migration induced by aged macrophage-conditioned medium (Figures 6, 7). Of note, rheumatoid arthritis is associated with increased atherosclerosis-related disease, especially among the aged (Crowson et al., 2013), and we suggest that some of the cardiovascular benefit of anti-TNFα drugs in this context (Dixon et al., 2007; Westlake et al., 2011) may result from limiting clonal expansion of SMC progenitors. On the other hand, *Itgb3* overexpression in the BM of aged mice via BMT reduces plasma TNFα and thus, prevents the clonal expansion of multiple SMC-derived progenitors in the plaque and attenuates plaque burden (Figure 7). It is intriguing to consider this result in light of a recent study demonstrating efficacy of autologous transplantation of genetically modified hematopoietic stem and progenitor cells as a treatment for patients with sickle cell disease (Kanter et al., 2022).

Aging is the predominant risk factor for atherosclerosis. We demonstrate that aged macrophages express reduced levels of TET2 inhibiting Itgb3 expression, and decreased integrin β3 in macrophages enhances TNFα levels which induces polyclonal expansion of SMCs in the atherosclerotic plaque and worsens disease. Thus, our studies put forth deficient regulation of SMC clonal expansion by aged BM-derived macrophages as a critical underlying factor in atherogenesis. Looking forward, this concept of impaired regulation of clonal expansion by aged BM (perhaps, via myeloid cell *Itgb3* deficiency) warrants intense investigation in other contexts, such as normal aging of the esophagus, endometrium, skin and bronchi as well as in cirrhosis and neurodegeneration (Brunner et al., 2019; Martincorena et al., 2018; Martincorena et al., 2015; Moore et al., 2020; Tay et al., 2017; Yoshida et al., 2020).

## METHODS

### Animals

Wild type (C57BL/6), *Ldlr^(-/-)^, Apoe^(-/-)^, Csf1r-Mer-iCre-Mer, ROSA26R^(mTmG/mTmG)^, Itgb3^(-/-)^* and *Itgb3^(flox/flox)^* mice were purchased from The Jackson Laboratory(Hodivala-Dilke et al., 1999; Morgan et al., 2010; Muzumdar et al., 2007; Qian et al., 2011). Mice carrying *Myh11-CreER^T2^* or the multi-color Rainbow (Rb) Cre reporter *ROSA26R^Rb^* have been described (Greif et al., 2012; Rinkevich et al., 2011; Wirth et al., 2008). Experiments with *Myh11-CreER^T2^* mice are restricted to males as this transgene incorporated on the Y chromosome (Wirth et al., 2008). Otherwise, studies utilized adult both male and female mice. Three month (young) or 18 month old (aged) mice were used for most experiments.

### Induction of atherogenesis and administration of AAV-*Pcsk9*

For atherosclerosis studies, we used mice null for *Ldlr*, *Apoe* and/or injected with recombinant adeno-associated virus (AAV) encoding constitutively active PCSK9 (*rAAV8.ApoEHCR-hAAT.D377Y-mPCSK9.bGH [*AAV*-Pcsk9]*); Penn Vector Core, University of Pennsylvania). For AAV*-Pcsk9* studies, one week after either tamoxifen treatment or, if relevant, four weeks after BMT, a single retro-orbital injection containing 1.0 x 10^11^ genome copies was administered. Four weeks later, mice were fed a WD (40% fat by calories, 1.25% cholesterol by weight; Research Diets Inc., D12108C) for up to 16 weeks. For mice not injected with AAV*-Pcsk9,* a WD was initiated four weeks after BMT.

### Bone marrow transplantation

One week after tamoxifen treatment, mice that were to receive a BMT were first lethally irradiated with two doses of 550 rads (5.5 Gy) from a X-RAD 320 unit (Precision X-ray) administered 4 h apart. BM was collected from femurs and tibias of donor mice by flushing with sterile Opti-MEM medium (Thermo Fisher). Each recipient mouse was injected retro-orbitally with 2 × 10^6^ BM cells. Four weeks after BMT, peripheral blood was collected by retro-orbital venous plexus puncture, and genomic DNA was extracted with QIAamp DNA Blood Mini Kit (Qiagen, 51104) for PCR analysis of BM reconstitution. Primer sequences are provided in Supplemental Table 2. Mice were then switched to a WD for up to 16 weeks and euthanized.

### Tissue preparation, histology and morphometric analysis

Following euthanasia, the mouse heart with proximal ascending aorta attached was perfused with 10 ml of phosphate-buffered saline (PBS) through the left ventricle and then incubated in 4% paraformaldehyde (PFA) overnight at 4°C. The tissue was then washed with PBS and placed in 30% sucrose in PBS at 4°C until the next day. Finally, the heart with attached aorta was embedded in OCT (Tissue Tek), frozen, and stored at -80°C. Serial cryosections of the aortic root were cut at 10 µm thickness using a cryostat. Every fifth pair of adjacent sections were stained with hematoxylin and eosin (H&E) and oil red O (ORO) for quantification of atherosclerotic plaque area and lipid deposition area, respectively. The size of the plaque and lipid areas for each mouse were obtained by averaging these areas from at least five sections. The necrotic core was measured as the average of acellular area of the plaque from three sections of the same mouse that were each at least 50 µm apart and reported as a percentage of the plaque area.

### Immunohistochemistry

The aortic root was cryosectioned in the transverse axis, and 10 µm sections were incubated with blocking solution (0.1% Triton X-100 in PBS (PBS-T) supplemented with 5% goat serum) and then with primary antibodies diluted in blocking solution overnight at 4°C. On the next day, sections were washed with PBS-T and then incubated with secondary antibodies for 1 h. Primary antibodies used were anti-GFP (1:500, Abcam, ab13970), anti-CD68 (1:200, Bio-Rad, MCA1957), directly conjugated Cy3 anti-SMA (1:500, Sigma-Aldrich, C6198). Secondary antibodies were conjugated to either FITC or Alexa-488, Alexa−647 fluorophores (1:500, Invitrogen). Nuclei were visualized by DAPI staining (1:1000, Sigma-Aldrich, D9542). Note, membrane-localized tdTomato (mT) fluorescence in the aorta of *ROSA26R^(mTmG/+)^*mice is very weak and essentially undetectable in comparison to the strong fluorescence in the red channel of staining with the anti-SMA antibody.

### Fate mapping, clonal analysis and proliferation in atherogenesis

Mice carrying an inducible Cre recombinase and a Cre reporter [*ROSA26R^(mTmG/+)^* or *ROSA26R^(Rb/+)^*] were induced with tamoxifen (1 mg/day) for 5 or 20 days in the case of *Myh11-CreER^T2^*or *Csf1r-Mer-iCre-Mer*, respectively and then rested for one week. Following BMT and/or injection with AAV*-Pcsk9*, mice were fed a WD for up to 16 weeks and euthanized. For in vivo proliferation studies, 2.5 mg of 5-ethynyl-2’ deoxyuridine (EdU) (Thermo Fisher Scientific, A10044) was injected intraperitoneally 12 h prior to euthanasia. Click-iT EdU Alexa Fluor 647 Imaging Kit (Thermo Fisher Scientific, C10340) was used as per manufacturer’s instructions to detect EdU incorporation into proliferating cells. Serial sections through the aortic root were stained for nuclei (DAPI). In the case of mice carrying the *ROSA26R^(mTmG/+)^* reporter, sections were also stained for GFP and SMA and either EdU or CD68. For *ROSA26R^(Rb/+)^*mice, sections were directly imaged using fluorescent filters for Cerulean, mOrange, and mCherry, and marked cells in the media and plaque were quantified by scoring for expression of these fluorophores. In all quantitative cellular studies, total cells were determined by counting DAPI^+^ nuclei.

### Anti-TNFα treatment

Mice underwent BMT, were rested for four weeks and then fed a WD for 12 weeks. During WD feeding, mice were intraperitoneally injected twice per week with mouse monoclonal anti-TNFα blocking antibody or IgG2a isotype matched control (Janssen Research and Development LLC, Spring House, PA) at a dose of 20 mg/kg body weight. For experiments with cultured cells, medium conditioned by young or aged BM-derived macrophages was pretreated with anti-TNFα antibody or IgG2a for 1 h at 37°C prior to initiating SMC migration or proliferation assays.

### Isolation of mononuclear cells and monocytes from BM of mice

BM cells were harvested by flushing the femurs and tibias of mice in PBS with 2% fetal bovine serum (FBS) on ice. Mononuclear cells were isolated from BM cells by density gradient centrifugation using Lympholyte (Cedarlane, CL0531). After centrifugation at 1300 g for 20 min, mononuclear cells were collected from the interface and washed in Ca^2+^/Mg^2+^ - free HBSS. The cells were dissociated to a single-cell suspension by filtering through a 70-µm nylon mesh. These mononuclear cells were subjected to scRNA-seq, monocyte isolation by FACS or differentiation to macrophages. For monocyte isolation, a suspension of mononuclear cells was stained with Alexa 700 anti-CD3 (1:100, BioLegend, 100215), PE-Cy7 anti-CD19 (1:100, BioLegend, 115520), FITC anti-CD11b (1:100, BD Pharmingen, 553310) and APC anti-Ly6c antibodies (1:100, BioLegend, 128015) in PBS with 0.5% FBS for 30 min. CD11b^+^Ly6c^+^ monocytes were isolated with a BD FACSAria ΙΙ cell sorter.

### Macrophage differentiation

Similar to our previous approach (Misra et al., 2018), BM-derived mononuclear cells were resuspended in Iscove’s DMEM with 20% FBS, 20% L-929 cell conditioned medium, and 25 μg/ml fungizone and plated at a density of 1.8 x 10^6^ cells/ml on uncoated plates. After 7 days of culture, non-adherent cells were washed away. Adherent cells were cultured in RPMI, 10% FBS for 12 h and then centrifuged at 1300 g for 5 min. The supernatant (i.e., macrophage conditioned medium) and pellet (i.e., macrophages) were used immediately or stored at -80°C.

### Human peripheral blood monocyte isolation

Human peripheral blood mononuclear cells were isolated by Ficoll (GE Healthcare) density gradient centrifugation and then stained with PE anti-CD3 (1:100, BioLegend, 300308), PE-Cy7 anti-CD19 (1:100, BD Pharmingen, 115520) and FITC anti-CD14 (1:100, BioLegend, 325604). Human CD14^+^CD3^-^CD19^-^ monocytes were sorted in a BD FACSAria ΙΙ cell sorter.

### Aortic SMC isolation

Aortic SMCs were isolated by modification of a previously described protocol (Karnik et al., 2003). Briefly, aortas from the root to the iliac bifurcation were harvested from adult wild type mice and opened longitudinally. Harvested aortas were enzymatically digested with 175 U/ml collagenase and 1.25 U/ml elastase (Worthington, LS004176 and LS002279, respectively) in PBS at 37°C in a shaking incubator for 30 min. The adventitia was manually peeled off, and the endothelium was removed by gentle scraping. Aortas were sequentially washed with 1% penicillin/streptomycin in PBS and then with 100% FBS. Washed aortas were cut into small pieces and cultured in plastic dishes in DMEM supplemented with 20% FBS, 1% penicillin/streptomycin. After 3 days, the medium was replaced by fresh medium that was the same except with 10% instead of 20% FBS. SMCs that migrated out of the aortic pieces and adhered to the dish were trypsinized, expanded, and passaged. SMCs were used until the fifth passage.

### Proliferation and migration assays in cell culture

For proliferation studies, wild type aortic SMC were cultured in DMEM, 10% FBS, 1% penicillin/streptomycin until they reached ∼60% confluency. Cells were then serum starved in DMEM overnight, followed by incubation with macrophage-conditioned medium for 8 h or 48 h. During the last 8 h of this incubation, EdU (10 µM) from the Click-iT EdU Alexa Fluor 647 Imaging Kit (Thermo Fisher Scientific) was added to the conditioned medium. Subsequently, cells were fixed with 4% PFA for 30 min, permeabilized in 0.5% Triton X-100 in PBS and stained for EdU and nuclei (DAPI).

The migration assay used cell culture inserts (Ibidi), composed of two chambers flanking a central insert that prevents cell growth. Wild type aortic SMCs were added to both chambers and allowed to attach and grow to confluency. SMCs were serum starved overnight and washed with PBS prior to removal of the insert and then cultured in macrophage-conditioned medium for 6 h and 8 h. The cell coverage of the area that was blocked by the insert was measured immediately after insert removal and 6 h and 8 h later.

### RNA isolation and quantitative real-time PCR

Total RNA was isolated from murine and human monocytes with the RNeasy Plus Micro Kit (Qiagen), and then 200 ng of RNA was reverse transcribed using the iScript RT Supermix (Bio-Rad), following instructions from the manufacturers. Quantitative real-time PCR was conducted in triplicate using SsoFast EvaGreen Supermix (Bio-Rad) on a Real Time Detection System (Bio-Rad). Transcript levels are relative to that of Gapdh. Forward and reverse primers are listed in Supplemental Table 3.

### Western blot

Murine cells or aortic tissue were lysed in RIPA buffer containing complete protease inhibitor and phosphoSTOP phosphatase cocktails (Roche Applied Science). Lysate was centrifuged at 10,000 g for 10 min at 4°C, and the supernatant was collected. After estimating protein concentration with the BCA assay (Pierce), proteins were separated by SDS-PAGE and transferred overnight to nitrocellulose membranes (Millipore). Membranes were blocked with 5% bovine serum albumin in tris-buffered saline with 0.05% Tween-20 (TBS-T) for 1 h and incubated with primary antibody overnight at 4°C. Membranes were washed with TBS-T incubated with HRP-conjugated secondary antibodies (Dako). After washing in TBS-T again, protein detection was performed with Super Signal West Pico Chemiluminescent Substrate (Thermo Scientific) and GBOX Imaging System (Syngene). Primary antibodies used for Western blot analysis were anti-integrin β3 (1:1000, Abcam, ab197662), anti-phospho (S473) AKT (1:1000, Cell Signaling Technology, 4060), anti-total AKT (1:1000, Cell Signaling Technology, 4691), anti-phospho PAK1 (1:1000, Cell Signaling Technology, 2606), anti-total PAK1 (1:1000, Cell Signaling Technology, 2602), anti-phospho ERK (1:1000, Cell Signaling Technology, 4376), anti-total ERK (1:1000, Cell Signaling Technology, 4695), anti-TET2 (1:1000, Abcam, ab213369), anti-TNFR1 (1:1000, Abcam, ab223352) and anti-GAPDH (1:1000, Cell Signaling Technology, 2118).

### Hydroxymethyl DNA immunoprecipitation

Hydroxymethyl DNA immunoprecipitation (hMeDIP) was carried out using a hMeDIP kit (Diagenode, C02010031) as previously published (Liu et al., 2013). Briefly, genomic DNA from murine or human monocytes was isolated using AllPrep DNA/RNA/miRNA Universal Kit (Qiagen, 80224). DNA (1 µg) was sheared with a sonicator, and DNA fragments (200 - 500 bp) were immunoprecipitated with a mouse monoclonal anti-5hmC antibody (2.5 μg per immunoprecipitation) following the protocol from Diagenode. The DNA-antibody mixture was incubated with magnetic beads overnight at 4°C, and then DNA was isolated. Multiple primers were designed to scan the proximal promoter region of *ITGB3* in murine and human monocytes. Open chromatin regions were identified using the UCSC genome browser (https://genome.ucsc.edu/). The forward and reverse primer pair spanning the proximal promoter region upstream of the transcription start site of mouse *Itgb3* were 5’-AGGATGCGAGCGCAGTG -3’ and 5’-CGCACCTCTGCTTCTCAGT -3’, respectively. The corresponding PCR amplification product is located at chr11:104,608,100 - 104,608,250. For human *ITGB3*, the forward and reverse primer pair spanning the promoter region upstream of the transcription start site were 5’-GAAGTGGTCAGGACCTGGAA-3’ and 5’-TTCTGTGCCACTAGCCTGAG-3’. The location of this PCR amplification product is chr17:47,240,300 - 47,240,400. Hydroxymethylated DNA and input fractions were analyzed with hMeDIP-qPCR using *ITGB3* primers per the manufacturer’s protocol to confirm enrichment of the hydroxymethylated gene.

### Construction of 10X Genomics single cell 3’ RNA-seq libraries and sequencing

#### a) Gel Beads-In-Emulsion generation and barcoding

Single cell suspension of BM-derived mononuclear cells in RT Master Mix (75 µl) was loaded on the Single Cell A Chip and partitioned with a pool of ∼750,000 barcoded gel beads to form nanoliter-scale Gel Beads-In-Emulsions (GEMs). Each gel bead has primers containing: (i) an Illumina R1 sequence (read 1 sequencing primer), (ii) a 16 nt 10x Barcode, (iii) a 10 nt Unique Molecular Identifier (UMI), and (iv) a poly-dT primer sequence. Upon dissolution of the Gel Beads in a GEM, the primers were released and mixed and incubated with cell lysate and RT Master Mix, producing barcoded, full-length cDNA from poly-adenylated mRNA.

#### b) Post GEM reverse transcription cleanup, cDNA amplification and library construction

Silane magnetic beads were used to isolate cDNA from leftover biochemical reagents and primers in the post GEM reaction mixture. Full-length, barcoded cDNA was then amplified by PCR to generate sufficient DNA for library construction. Enzymatic fragmentation and size selection were used to optimize cDNA amplicon size prior to library construction. R1 (read 1 primer sequence) were added during GEM incubation. P5, P7, a sample index, and R2 (read 2 primer sequence) were added during library construction via end repair, A-tailing, adaptor ligation and PCR. The final libraries contain the P5 and P7 primers used in Illumina bridge amplification.

#### c) Sequencing libraries

The single cell 3’ library comprised standard Illumina paired-end constructs which begin and end with P5 and P7. The single cell 3’ 16 bp 10x Barcode and 10 bp UMI were encoded in Read 1 and Read 2 was used for sequencing the cDNA fragment. Sequencing the single cell 3’ library produced standard Illumina Binary Base Call data which includes the paired-end Read 1 and Read 2 and the sample index in the i7 index read.

#### d) scRNA-seq data processing and analysis

Cellranger v3.1.0 was used to align scRNA-seq samples to the mouse genome (mm10). Downstream analysis was performed using Seurat v3.1.5 R package, including cell type identification and comparative analyses between the *Apoe^(-/-)^, Itgb3^(-/-)^* and *Apoe^(-/-)^, Itgb3^(+/+)^* samples. For quality control, cells with the number of expressed genes <100 or >6000 were filtered out. Cells were also excluded if their mitochondrial gene percentages were over 50%. For each sample, we first normalized the raw count matrix and then defined 2000 top variable genes. Seurat Integration was leveraged to find common anchoring features between these two samples and integrate them with default settings. We then applied principal component analysis for dimensionality reduction and retained 30 leading principal components for further visualization and cell clustering. The t-distributed stochastic neighbor embedding (t-SNE) projection (Linderman et al., 2019) was used to visualize the cells on a two-dimensional space with a perplexity of 100. Subsequently, the share nearest neighbor graph was constructed by calculating the Jaccard index between each cell and its 20-nearest neighbors, which was then used for cell clustering based on Louvain algorithm (with a resolution of 0.125). After identifying cluster-specific genes, we annotated cell types based on some canonical marker genes. We removed erythroblasts and screened each cell type for differentially expressed genes between the two conditions. The gene lists served as the input to KEGG gene set enrichment analysis by using clusterProfiler v3.14.3 R package. IPA (Version 52912811, Ingenuity Systems, QIAGEN) was used to delineate diseases and functions represented by differentially expressed genes.

### Bulk RNA sequencing

#### a) RNA isolation and quality control

Total RNA from FACS-isolated monocytes was obtained using PureLink^TM^ RNA Minikit (Invitrogen). RNA quality was determined by measuring A260/A280 and A260/A230 ratios via nanodrop. RNA integrity was determined by running an Agilent Bioanalyzer gel to measure the ratio of ribosomal peaks. Samples used for library preparation had RIN values greater than 8.

#### b) Bulk RNA-seq library preparation

mRNA was purified from ∼200 ng of total RNA with oligo-dT beads and sheared by incubation at 94°C in the presence of Mg^2+^ with the KAPA mRNA HyperPrep kit (Roche). Following first-strand synthesis with random primers, second strand synthesis and A-tailing were performed with dUTP to generate strand-specific sequencing libraries. Adapter ligation with 3’ dTMP overhangs were ligated to library insert fragments. Library amplification of fragments carrying the appropriate adapter sequences at both ends was undertaken. Strands marked with dUTP were not amplified. Indexed libraries were quantified by qRT-PCR using a commercially available kit (Roche, KAPA Biosystems) and insert size distribution was determined with the Agilent Bioanalyzer. Samples with a yield of ≥ 0.5 ng/µl and a size distribution of 150-300 bp were used for sequencing.

#### c) Flow cell preparation and sequencing

Samples at a concentration of 1.2 nM were loaded onto an Illumina NovaSeq6000 flow cell to yield 25 million passing filter clusters per sample. Samples were sequenced using 100 bp paired-end sequencing per Illumina protocols. Reads were trimmed to remove low quality base calls.

#### d) Bulk RNA-seq analysis

STAR v2.5.3 was used to align the raw sequencing reads to the mouse genome (mm10) with default parameters. The number of reads for each gene was then counted with HTSeq v0.6.1. Subsequently, we used DESeq2 v1.26.0 R package to identify significantly differential expressed genes between aged and young BM-derived monocytes and then performed KEGG gene set enrichment analysis (GSEA) with clusterProfiler v3.14.3 R package.

### Mouse *Itgb3* cloning and lentivirus production

Mouse *Itgb3* was cloned into the pLVX-EF1α-IRES-Puro vector (Takara Bio) per a previously described cloning strategy (Dave et al., 2016). Briefly, total RNA was isolated from mouse aorta using PureLink^TM^ RNA Minikit (Invitrogen), and cDNA was synthesized with ProtoScript II First Strand cDNA Synthesis Kit (New England Biolabs) using 1 µg RNA. PCR was performed using 1 µl of cDNA template with AccuPrime Taq HIFI Polymerase (Invitrogen) in a 50 µl reaction volume using *Itgb3* specific cloning primers to incorporate EcoRI and BamHI flanking sites. Primers used were: Itgb3-EcoRI forward: 5’ AGCAGAATTCATGCGAGCGCAGTGG 3’ and Itgb3-BamHI reverse: 5’AGCAGGATCCTTAAGTCCCCCGGTAGGTG 3’. The *Itgb3* amplicons were cleaned with the Qiagen QIAquick PCR Purification Kit, digested with EcoRI and BamH1 along with the pLVX-EF1α-IRES-Puro vector at 37°C for 1 h, gel purified with the Qiagen QIAquick Gel Extraction Kit and ligated overnight at 14°C using T4 ligase. Positive clones were verified by sequencing.

Lenti-X Expression System (Takara Bio) was utilized to generate lentivirus as per manufacturer’s instructions. Briefly, 5x10^6^ Lenti-X 293T cells were co-transfected with Lenti-X Packaging Single Shots VSV-G and 7 µg of pLVX-EF1α-IRES-Puro vector with or without *Itgb3* insert. Lentiviral supernatants were collected 48 h post-transfection and concentrated by adding one volume of Lenti-X Concentrator (Takara Bio) to three volumes of lentivirus-containing supernatant and incubating at 4°C overnight. On the next day, lentiviral pellet was obtained by centrifuging the mixture at 1500xg for 45 min, resuspended in 1 ml sterile PBS, aliquoted and stored in -80°C. Prior to transplant, BM cells purified by Ficoll density gradient centrifugation were transduced with empty vector (control) or *Itgb3* overexpressing lentivirus in X-VIVO 15 medium (Biowhittaker) supplemented with SCF (100 ng/ml), TPO (50 ng/ml), Flt3 ligand (50 ng/ml) and IL-3 (20 ng/ml) onto plates coated with retronectin (Takara Bio).

### Imaging

Images were acquired with Nikon microscopes (Eclipse 80i upright fluorescent or Eclipse TS100 inverted) or Leica SP8 confocal microscope. For image processing, analysis, and cell counting, Adobe Photoshop and Image J software were used.

### Quantification and Statistical Analysis

Two-tailed Student t-test and one-way ANOVA with Tukey’s multiple comparison test were used to analyze data (GraphPad Prism 8). Data are presented as average ± SD. A p-value of ≤ 0.05 was considered statistically significant.

### Study Approval

All mouse experiments were approved by the IACUC at Yale University and performed in accordance with relevant ethical guidelines. All procedures involving human subjects were approved by the Institutional Review Boards of Yale University (IRB# 1005006865), and we complied with all relevant ethical regulations. Written informed consent was obtained from all participants prior to inclusion in the study.

## Supporting information

https://www.biorxiv.org/content/biorxiv/early/2022/01/21/2022.01.18.476756/DC1/embed/media-1.pdf?download=true

https://www.biorxiv.org/content/biorxiv/early/2022/01/21/2022.01.18.476756/DC2/embed/media-2.pdf?download=true

## ACKNOWLEDGEMENTS

We thank Greif laboratory members and Jeff Testani for input. I.K. was supported by a Postdoctoral Fellowship from the American Heart Association (18POST34030015). Funding was also provided by the NIH (R35HL150766, R01HL125815, R01HL142674, R21AG062202, 1R21NS123469 to D.M.G.), American Heart Association (Established Investigator Award, 19EIA34660321 to D.M.G.). No conflicts of interest.

## AUTHOR CONTRIBUTIONS

I.K., J.D., R.C., R.Q., N.R., K.A.M., C.F.H., and D.M.G. conceived of and designed experiments. I.K., X.Z., J.D., R.C., R.R.C., A.N., E. G-V., B.A. and N.R. performed them. R.Q. and Y.K. conducted scRNA-seq analysis, and R.G-M. helped with bulk RNA-seq analysis. J.H. provided infrastructure for access to human blood. I.K. and D.M.G. analyzed the results, prepared the figures and wrote the manuscript. All authors reviewed and provided input on the manuscript.

