## Supplementary material for "Age of the bone marrow dictates clonality of smooth muscle-derived cells in the atherosclerotic plaque": https://www.biorxiv.org/content/biorxiv/early/2022/01/21/2022.01.18.476756/DC1/embed/media-1.pdf?download=true

**List of Supplemental Items:**

- Supplemental Figures S1-S7
- Supplemental Legends S1-S7
- Supplemental Tables S1-5
- Supplemental Methods

Young, *Apoe(-/-), Myh11-CreER, ROSA26R(Rb/+)*, BMT (Young vs Aged), AAV-*Pcsk9*, WD 16 Weeks

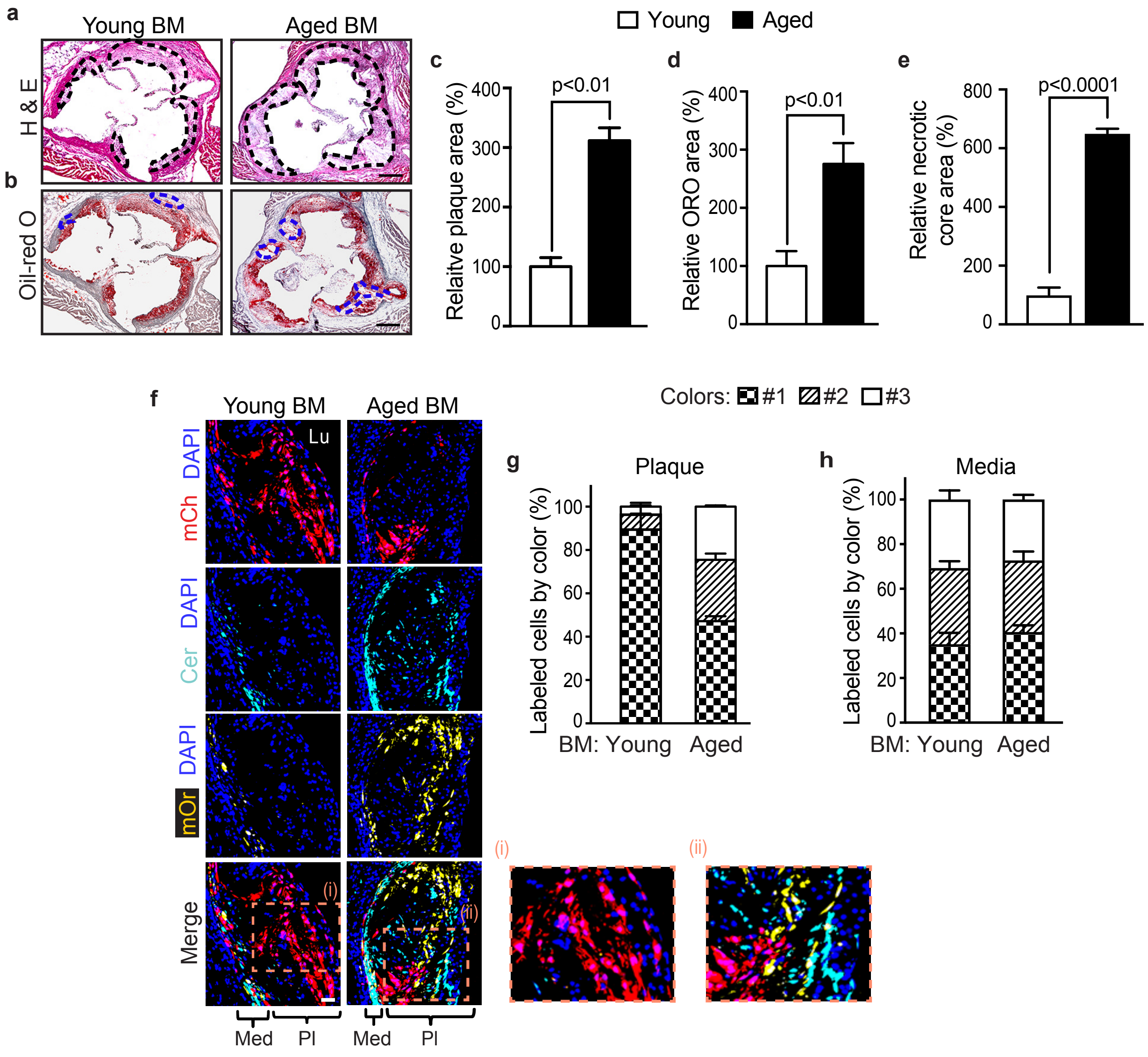


SMC culture in Mø-conditioned medium, 8 h

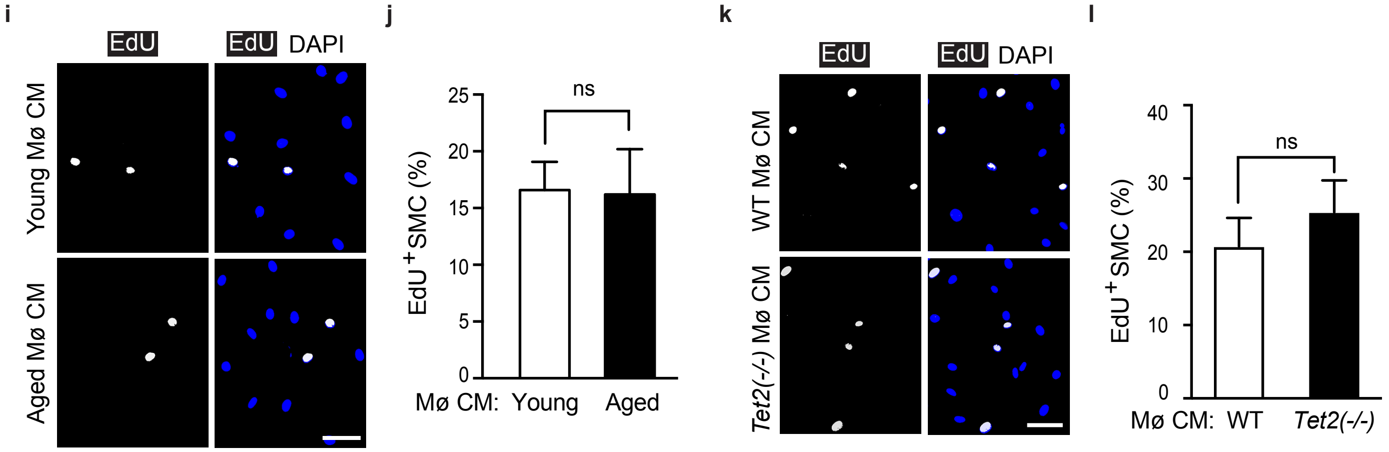

Aged, *Myh11-CreER, ROSA26R(Rb/+),* AAV-*Pcsk9*, BMT (Young vs Aged), WD 16 Weeks

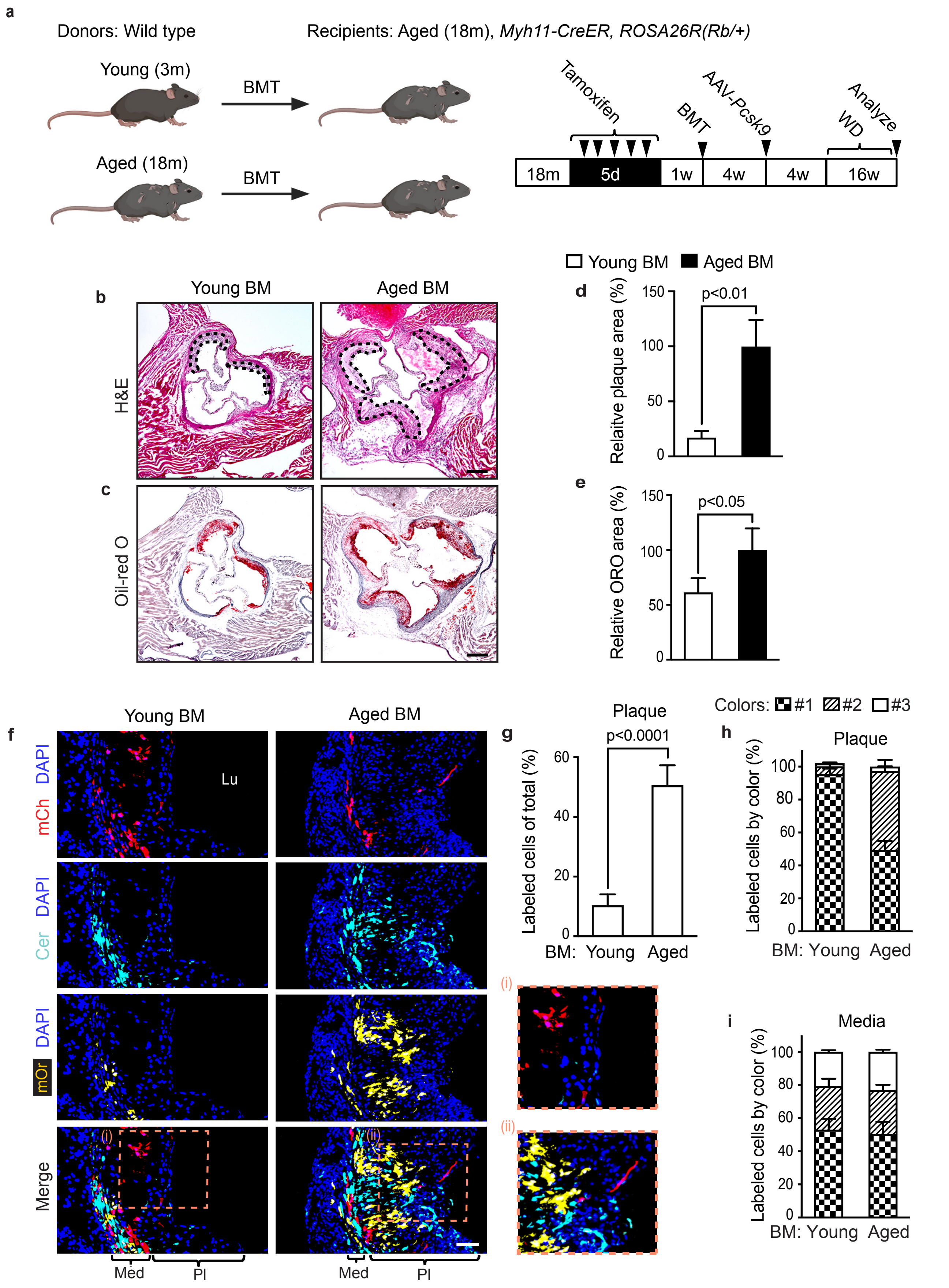

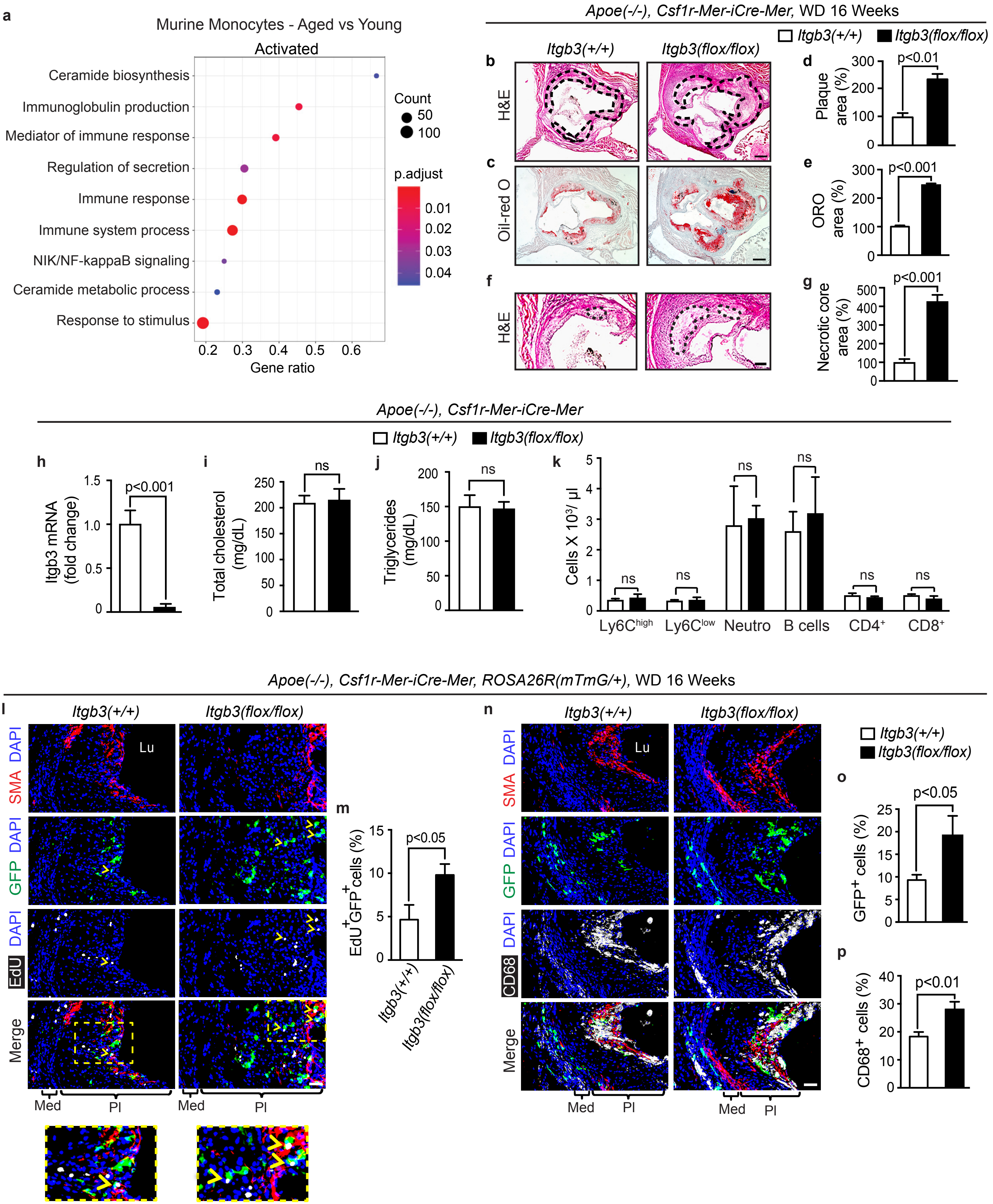
a
Murine Monocytes - Aged vs Young
Activated
Ceramide biosynthesis
Immunoglobulin production
Mediator of immune response
Regulation of secretion
Immune response
Immune system process
NIK/NF-kappaB signaling
Ceramide metabolic process
Response to stimulus
Gene ratio
Count
50
100
p.adjust
0.01
0.02
0.03
0.04
Apoe(-/-), Csf1r-Mer-iCre-Mer, WD 16 Weeks
Itgb3(+/+)
Itgb3(flox/flox)
b
c
d
e
f
g
H&E
Oil-red O
Plaque area (%)
ORO area (%)
Necrotic core area (%)
p<0.01
p<0.001
p<0.001
Apoe(-/-), Csf1r-Mer-iCre-Mer
h
i
j
k
Itgb3 mRNA (fold change)
Total cholesterol (mg/dL)
Triglycerides (mg/dL)
Cells X 10³/ µl
ns
Ly6C^high
Ly6C^low
Neutro
B cells
CD4⁺
CD8⁺
Apoe(-/-), Csf1r-Mer-iCre-Mer, ROSA26R(mTmG/+), WD 16 Weeks
l
m
n
o
p
Lu
DAPI
SMA
GFP
EdU
CD68
Merge
Med
Pl
EdU⁺GFP⁺cells (%)
p<0.05
GFP⁺ cells (%)
CD68⁺ cells (%)
p<0.01

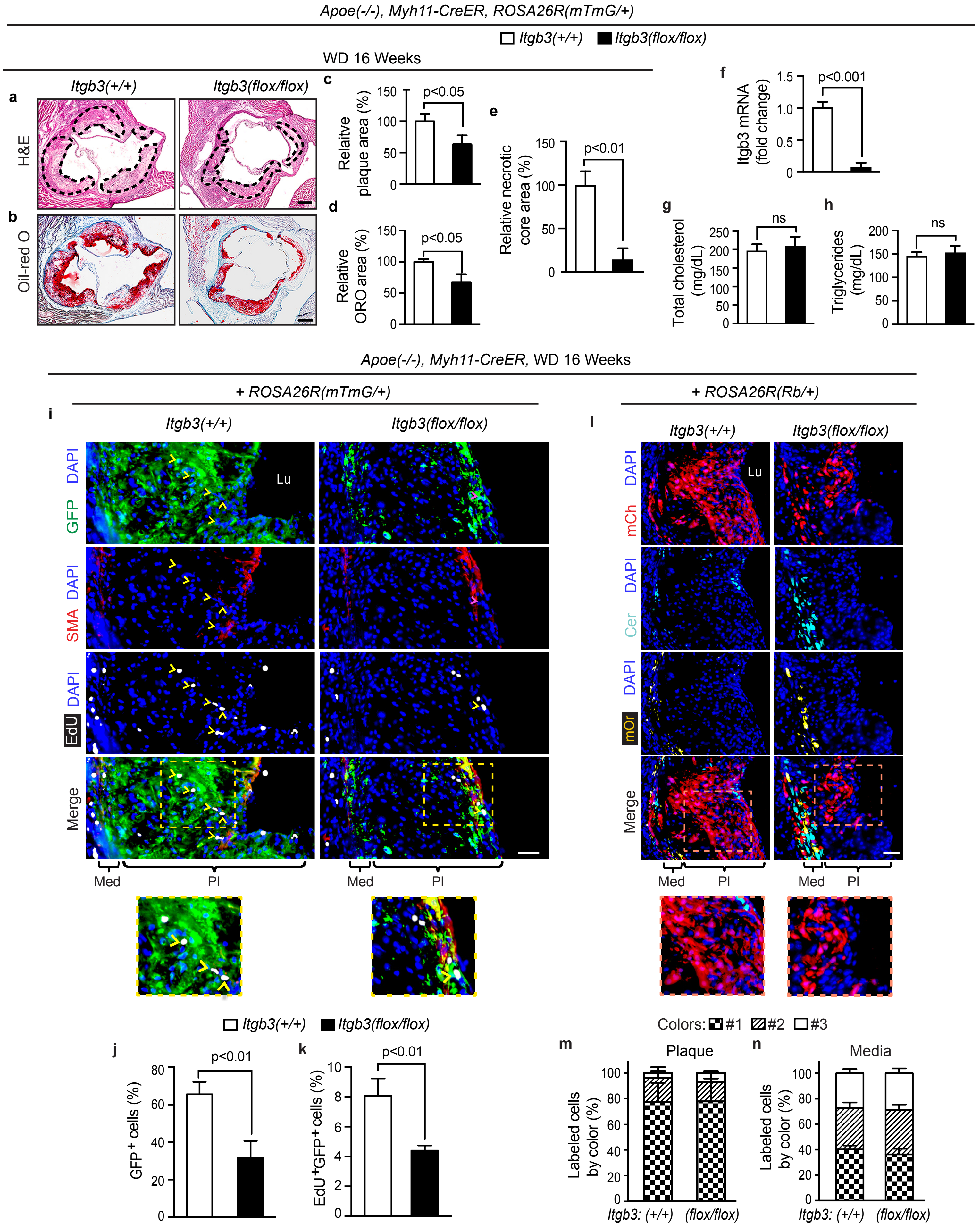
Apoe(-/-), Myh11-CreER, ROSA26R(mTmG/+)
Itgb3(+/+) Itgb3(flox/flox)
WD 16 Weeks
a
b
Itgb3(+/+)
Itgb3(flox/flox)
H&E
Oil-red O
c
Relative plaque area (%)
p<0.05
d
Relative ORO area (%)
p<0.05
e
Relative necrotic core area (%)
p<0.01
f
Itgb3 mRNA (fold change)
p<0.001
g
Total cholesterol (mg/dL)
ns
h
Triglycerides (mg/dL)
ns
Apoe(-/-), Myh11-CreER, WD 16 Weeks
+ ROSA26R(mTmG/+)
+ ROSA26R(Rb/+)
i
l
DAPI GFP
DAPI SMA
DAPI EdU
Merge
Lu
DAPI mCh
DAPI Cer
DAPI mOr
Med
Pl
j
GFP+ cells (%)
p<0.01
k
EdU+GFP+ cells (%)
p<0.01
Colors: #1 #2 #3
m
Plaque
n
Media
Labeled cells by color (%)
Itgb3: (+/+) (flox/flox)

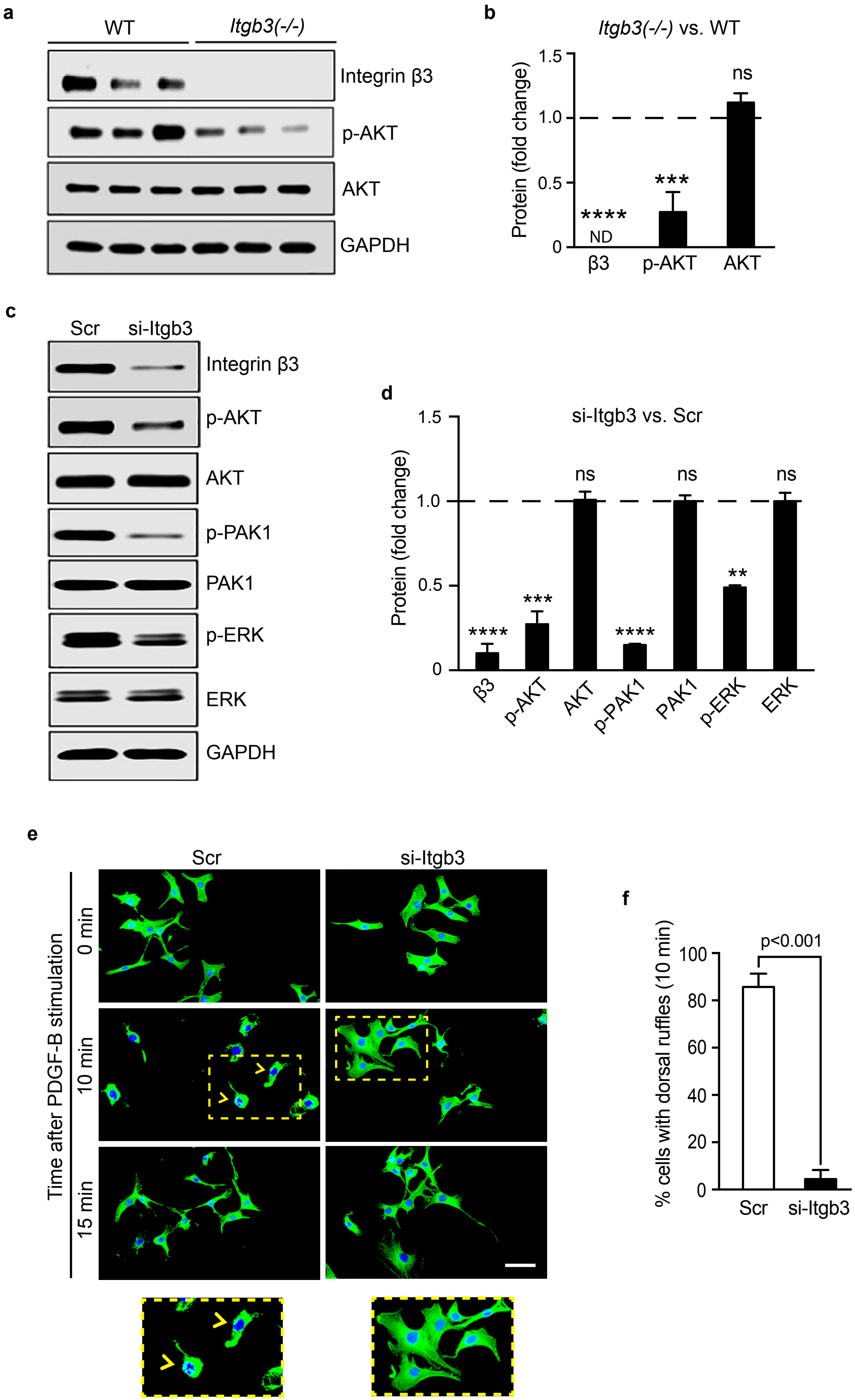
a
WT
Itgb3(-/-)
Integrin β3
p-AKT
AKT
GAPDH
b
Itgb3(-/-) vs. WT
Protein (fold change)
1.5
1.0
0.5
0
ns
***
****
ND
β3
p-AKT
AKT
c
Scr
si-Itgb3
Integrin β3
p-AKT
AKT
p-PAK1
PAK1
p-ERK
ERK
GAPDH
d
si-Itgb3 vs. Scr
Protein (fold change)
1.5
1.0
0.5
0
ns
ns
ns
****
***
****
**
β3
p-AKT
AKT
p-PAK1
PAK1
p-ERK
ERK
e
Scr
si-Itgb3
Time after PDGF-B stimulation
0 min
10 min
15 min
f
% cells with dorsal ruffles (10 min)
p<0.001
100
80
60
40
20
0
Scr
si-Itgb3

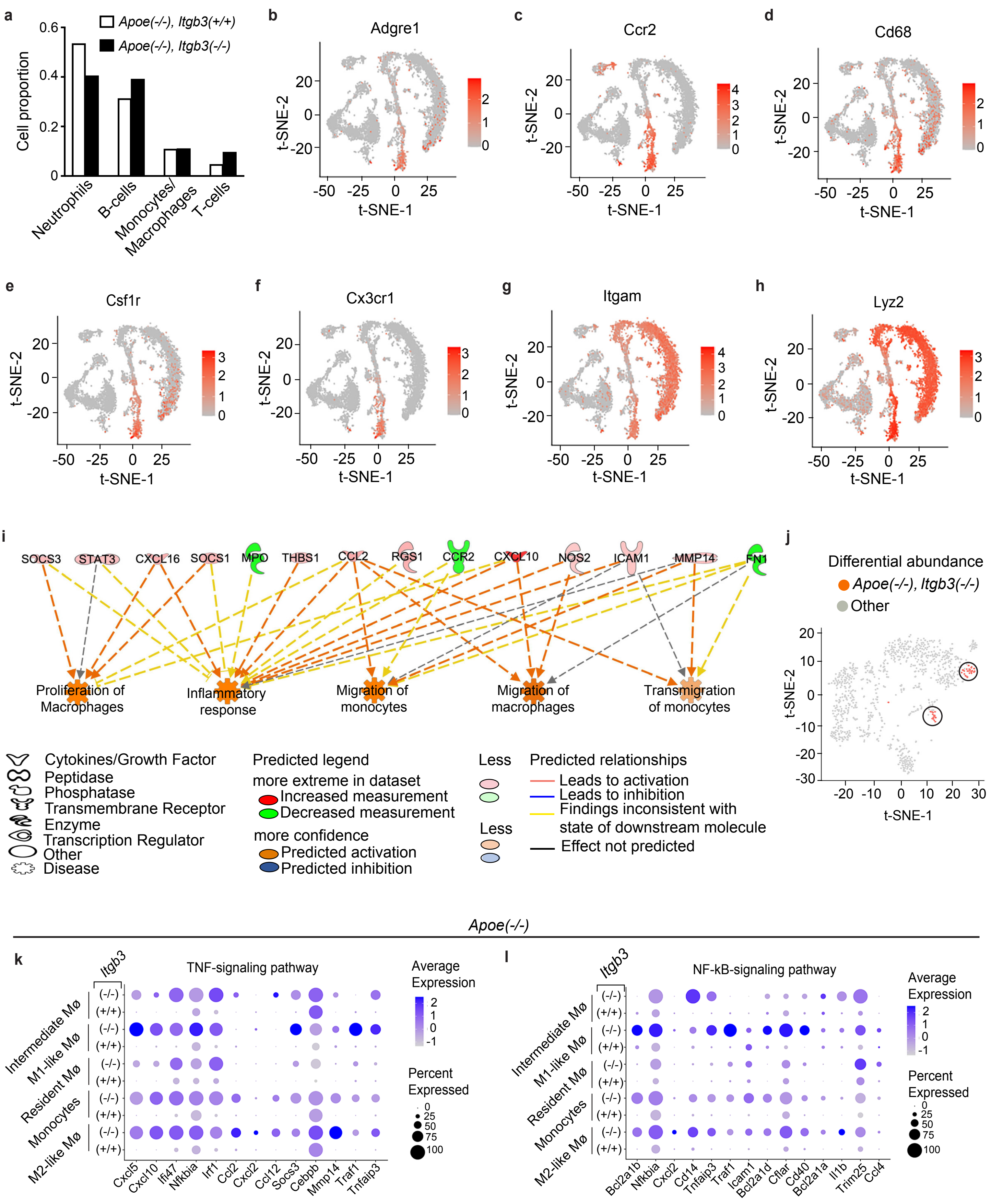
a
Apoe(-/-), Itgb3(+/+)
Apoe(-/-), Itgb3(-/-)
Cell proportion
Neutrophils
B-cells
Monocytes/ Macrophages
T-cells
b Adgre1
c Ccr2
d Cd68
e Csf1r
f Cx3cr1
g Itgam
h Lyz2
t-SNE-1
t-SNE-2
i
SOCS3 STAT3 CXCL16 SOCS1 MPO THBS1 CCL2 RGS1 CCR2 CXCL10 NOS2 ICAM1 MMP14 FN1
Proliferation of Macrophages
Inflammatory response
Migration of monocytes
Migration of macrophages
Transmigration of monocytes
Cytokines/Growth Factor
Peptidase
Phosphatase
Transmembrane Receptor
Enzyme
Transcription Regulator
Other
Disease
Predicted legend
more extreme in dataset
Increased measurement
Decreased measurement
more confidence
Predicted activation
Predicted inhibition
Less
Less
Predicted relationships
Leads to activation
Leads to inhibition
Findings inconsistent with state of downstream molecule
Effect not predicted
j
Differential abundance
Apoe(-/-), Itgb3(-/-)
Other
Apoe(-/-)
k
Itgb3
TNF-signaling pathway
Intermediate Mø
M1-like Mø
Resident Mø
Monocytes
M2-like Mø
(-/-)
(+/+)
Cxcl5 Cxcl10 Ifi47 Nfkbia Irf1 Ccl2 Cxcl2 Ccl12 Socs3 Cebpb Mmp14 Traf1 Tnfaip3
Average Expression
Percent Expressed
l
NF-kB-signaling pathway
Bcl2a1b Nfkbia Cxcl2 Cd14 Tnfaip3 Traf1 Icam1 Bcl2a1d Cflar Cd40 Bcl2a1a Il1b Trim25 Ccl4

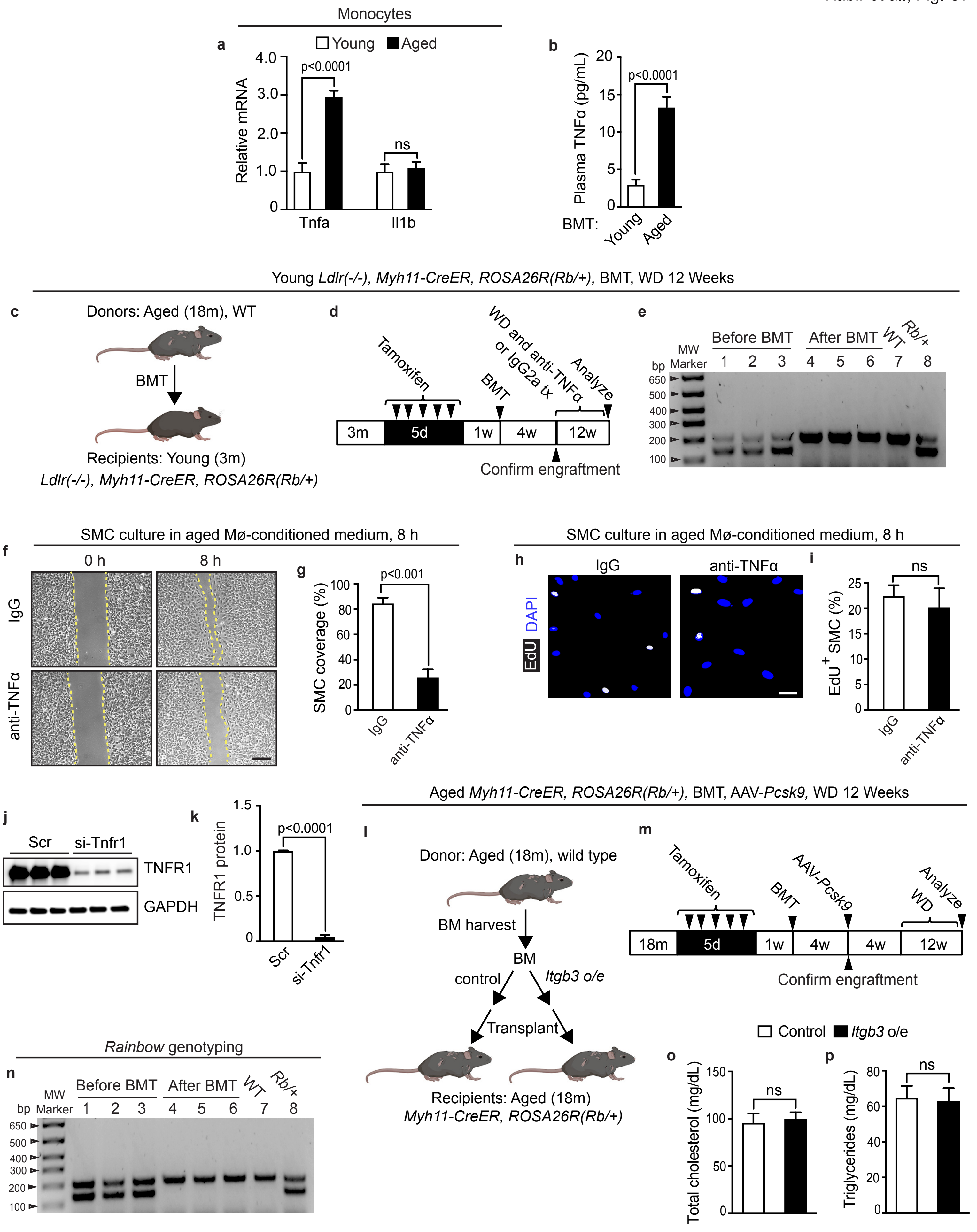

Monocytes
a
Young
Aged
Relative mRNA
p<0.0001
ns
Tnfa
Il1b
b
Plasma TNFα (pg/mL)
p<0.0001
BMT:
Young
Aged
Young Ldlr(-/-), Myh11-CreER, ROSA26R(Rb/+), BMT, WD 12 Weeks
c
Donors: Aged (18m), WT
BMT
Recipients: Young (3m)
Ldlr(-/-), Myh11-CreER, ROSA26R(Rb/+)
d
Tamoxifen
BMT
WD and anti-TNFα or IgG2a tx
Analyze
3m
5d
1w
4w
12w
Confirm engraftment
e
Before BMT
After BMT
WT
Rb/+
bp
MW Marker
SMC culture in aged Mø-conditioned medium, 8 h
f
0 h
8 h
IgG
anti-TNFα
g
SMC coverage (%)
p<0.001
IgG
anti-TNFα
SMC culture in aged Mø-conditioned medium, 8 h
h
IgG
anti-TNFα
EdU
DAPI
i
ns
EdU+ SMC (%)
IgG
anti-TNFα
j
Scr
si-Tnfr1
TNFR1
GAPDH
k
TNFR1 protein
p<0.0001
Scr
si-Tnfr1
Aged Myh11-CreER, ROSA26R(Rb/+), BMT, AAV-Pcsk9, WD 12 Weeks
l
Donor: Aged (18m), wild type
BM harvest
BM
control
Itgb3 o/e
Transplant
Recipients: Aged (18m)
Myh11-CreER, ROSA26R(Rb/+)
m
Tamoxifen
BMT
AAV-Pcsk9
Analyze
WD
18m
5d
1w
4w
4w
12w
Confirm engraftment
Rainbow genotyping
n
Before BMT
After BMT
WT
Rb/+
bp
MW Marker
Control
Itgb3 o/e
o
Total cholesterol (mg/dL)
ns
p
Triglycerides (mg/dL)
ns

### Supplemental Figure Legends

**Figure S1. Aged BMT promotes polyclonal expansion of SMC progenitors, and culturing for 8 hours with aged or *Tet2*$^{(-/-)}$ macrophage conditioned medium does not alter SMC proliferation. a-h,** Young (3 month old) *Apoe*$^{(-/-)}$, *Myh11-CreER*$^{T2}$, *ROSA26R*$^{(Rb/+)}$ mice were induced with tamoxifen, irradiated and transplanted with bone marrow (BM) from young or aged (18 month old) mice, injected with AAV-*Pcsk9* to reduce hepatic LDLR receptor level and fed a WD for 16 weeks. Representative histological analysis of transverse sections of the aortic sinus stained with H&E and Oil Red O are shown in **a, b**, respectively, with quantification of the lesion area **(c;** demarcated by dashed lines in **b)**, lipid content **(d)** and necrotic core area **(e)** in the plaques. The transverse aortic root sections were stained with DAPI and directly imaged for Rb colors (mCherry, mOrange and Cerulean) in **f**, and of the marked plaque **(g)** or medial cells **(h)**, the percent of cells of each color was quantified for each BM transplant (BMT) group. In a given plaque (or media), color 1 is the most dominant color of cells; color 2 is the second most common color; and color 3 is the least frequent color. n=5 mice and 12 plaques per group, 5-6 sections with a total of ~1000-1150 cells and spanning 200 μm per plaque. Lu, lumen; Med, tunica media; Pl; plaque. **i-l,** Murine aortic SMCs were cultured for 8 h in the presence of EdU in medium conditioned by BM-derived macrophages of either young vs. aged wild type (WT) mice or WT vs. *Tet2*$^{(-/-)}$ mice. In **i, k,** SMCs were stained for EdU and nuclei (DAPI), and in **j, l,** percent of DAPI$^{+}$ nuclei that also express EdU was quantified. n=3 mice per group. All data are averages $\pm$ SD. Student's *t*-test was used. ns, not significant. Scale bars, 100 μm **(a, b)**, 50 μm **(f)** and 25 μm **(i, k).**

**Figure S2. Aged BM induces polyclonal expansion of SMMHC$^+$ progenitors in the plaque.** Aged (18 month) *Myh11-CreER$^{T2}$*, *ROSA26R$^{(Rb/+)}$* mice were induced with tamoxifen, irradiated, transplanted with BM from young (3 month) or aged mice, injected with AAV-*Pcsk9* and fed a WD for 16 weeks. Transverse aortic root sections were analyzed. **a,** Schematic of experiments. **b-e,** Sections were stained with H&E **(**dashed lines demarcate lesion; **b)** and Oil-red O **(c)** to determine lesion area **(d)** and lipid content **(e)**, respectively. n=5-7 mice per BMT group, triplicate measurements from each mouse. **f-i,** Sections were stained with DAPI and directly imaged for Rb colors (mCh, mOr, Cer). In **f,** representative sections are shown with close-ups of boxed regions displayed on the right. In **g,** percent of DAPI$^+$ plaque cells that were marked by any of the Rb colors was quantified. Of the marked plaque **(h)** or underlying medial cells **(i)**, percent of cells of each color was quantified for each BMT group. In a given plaque (or media), color 1 is color with the greatest number of cells in plaque (or media), color 2 is second most common color and color 3 is least frequent color. n=6-8 mice and 14 plaques per BMT group, 5 sections with a total of ~1000-1200 cells and spanning 200 μm per plaque. Lu, lumen; Med, media; Pl, plaque. All data are averages $\pm$ SD, and Student's *t*-test was used. Scale bars, 100 μm **(b, c)** and 50 μm **(f)**.

**Figure S3. Deletion of *Itgb3* in CSF1R$^+$ cells exacerbates atherosclerosis and induces proliferation and accumulation of this cell lineage in the plaque. a,** Lysates of monocytes isolated from young or aged mice underwent bulk RNA-seq (n=3 mice per age group), and the transcriptome was subjected to gene set enrichment analysis. Activated pathways are shown with gene ratio indicating the number of activated genes relative to the total number of genes in the corresponding pathways and size of the dot representing number of activated genes.

**b-k,** *Apoe$^{(-/-)}$*, *Csf1r-Mer-iCre-Mer* mice also carrying *Itgb3$^{(flox/flox)}$* or wild type for *Itgb3* were induced with tamoxifen. In **b-g,** mice were then fed a WD for 16 weeks, and transverse aortic root sections were stained with H&E **(**dashed lines demarcate lesion and necrotic core in **b** and **f,** respectively) and Oil Red O **(c)**. Quantification of lesion area **(d)**, lipid content **(e)** and necrotic core area (**g**). n=5 mice and 10-12 plaques per genotype (triplicate measurements per plaque). In **h-k,** mice were analyzed after a 5 day rest period. BM-derived GFP$^{+}$ myeloid cells (Ly6C$^{+}$) were isolated by FACS and subjected to qRT-PCR for Itgb3 (**h**). Itgb3 mRNA is represented as relative to Gapdh and normalized to control. n=9 mice per group, and qRT-PCR was done in triplicate. Total cholesterol and triglycerides levels were assessed in blood plasma after 16 h of fasting, respectively (n=5-7 mice per group; **i, j**). Leukocyte counts were analyzed from peripheral blood of *Itgb3$^{(+/+)}$* and *Itgb3$^{(flox/flox)}$* mice (**k**). n=5-7 mice. **l-p,** *Apoe$^{(-/-)}$, Csf1r-Mer-iCre-Mer, ROSA26R$^{(mTmG/+)}$* mice also carrying *Itgb3$^{(+/+)}$* or *Itgb3$^{(flox/flox)}$* were induced with tamoxifen and rested. **l, n,** After 16 weeks of WD feeding, EdU was injected intraperitoneally 12 h prior to euthanasia. Transverse aortic root sections were stained for markers of fate (GFP), SMCs (α-smooth muscle actin [SMA]), nuclei (DAPI) and either proliferation (EdU) in **l** or CD68 in **n**. Boxed regions in **l** are shown as close-ups below with arrowheads indicating GFP$^{+}$EdU$^{+}$ cells. In **m,** the percentage of GFP$^{+}$ plaque cells that are EdU$^{+}$ is shown. In, **o, p,** percentage of plaque cells that are GFP$^{+}$ or CD68$^{+}$ was quantified, respectively. n=5-7 mice and 10 plaques per genotype, 4 sections with a total of ~1000-1250 cells and spanning 150 µm per plaque. All data are averages ± SD, and Student's *t*-test was used. ns, not significant. Lu, lumen; Med, media; Pl, plaque. Scale bars, 100 µm **(b, c, f)**, 50 µm **(l, n)**.

**Figure S4. SMC-specific *Itgb3* deletion attenuates atherosclerosis and SMC contribution to the plaque but does not alter plasma lipids.** *Apoe*$^{(-/-)}$, *Myh11-CreER*$^{T2}$ mice also carrying *Itgb3*$^{(flox/flox)}$ or *Itgb3*$^{(+/+)}$ and either *ROSA26R*$^{(mTmG/+)}$ **(a-k)** or *ROSA26R*$^{(Rb/+)}$ **(l-n)** were induced with tamoxifen. **a-e** and **i-n**, Mice were rested, subjected to WD for 16 weeks and aortic root transverse sections were analyzed. In **a-e,** sections were stained with H&E [dashed lines demarcating lesion; **(a)**] and Oil Red O **(b)**, and lesion area **(c)**, lipid content **(d)** and necrotic core area **(e)** were quantified. n=7-8 mice per group. In **i,** EdU was injected intraperitoneally 12 h prior to euthanasia. Sections were stained for SMA, GFP (fate marker) and nuclei (DAPI). Close-ups of boxed regions are shown below. Arrowheads indicate GFP$^{+}$EdU$^{+}$ cells. Percent of total plaque cells that are GFP$^{+}$ and percent of total plaque GFP$^{+}$ cells that are EdU$^{+}$ are shown in **j** and **k**, respectively. n=5 mice and 10 plaques per genotypes, 5-6 sections with a total of ~1200-1500 cells and spanning 200 μm per plaque. In **l**, sections were stained with DAPI and directly imaged for Rb colors with close-ups of boxed regions below. Of the marked plaque **(m)** or underlying medial cells **(n)**, the percent of cells of each color was quantified. In a given plaque (or media), color 1 is the most dominant color of cells in plaque (or media), color 2 is the second most prevalent and color 3 is least frequent. n=5 mice and 12 plaques per genotype, 6 sections with a total of ~1000-1300 cells and spanning 250 μm per plaque. Lu, lumen; Med, media; Pl, plaque. **f-h,** Mice were analyzed after a 5 day rest. In **f,** GFP$^{+}$ SMCs were isolated by FACS and subjected to qRT-PCR for Itgb3. Itgb3 mRNA is represented relative to Gapdh and normalized to *Itgb3*$^{(+/+)}$ group. n=6 mice per group, and qRT-PCR was done in triplicate. After 16 h fasting, plasma total cholesterol and triglycerides were measured in **g** and **h**, respectively; n=5-7 mice per group. All data are averages ± SD, and Student's *t*-test was used. Scale bars, 100 μm **(a, b)** and 50 μm **(i, l)**.

**Figure S5. Integrin β3 reduction in aortic SMCs attenuates phosphorylation of AKT and ERK and PDGF-B-induced dorsal ruffles**. **a, b,** Aortic lysates of wild type (WT) or *Itgb3*$^{(-/-)}$ mice were subjected to Western blot for integrin β3, GAPDH and phosphorylated (p-) and total AKT **(a)** with densitometry of each protein relative to GAPDH and normalized to WT **(b)**. n=3 mice per genotype. **c-f,** Mouse aortic SMCs were isolated and treated with Itgb3 siRNA or scrambled (Scr) RNA. Cell lysates were subjected to Western blot for integrin β3, GAPDH as well as phosphorylated and total AKT, PAK1 and ERK **(c)** with densitometry of each protein relative to GAPDH and normalized to Scr **(d)**. In **e**, following siRNA treatment, SMCs were serum starved overnight, treated with PDGF-B (10 ng/mL) for the indicated times and stained with FITC-phalloidin. Arrowheads indicate dorsal ruffles with boxed regions shown as close-ups below. Scale bar, 25 μm. Quantification of dorsal ruffles is shown in **(f)**. n=3 experiments, each experiment done in triplicate. All data are averages ± SD, and Student's *t*-test was used. ns, not significant.

**Figure S6. scRNA-seq analysis of *Apoe*$^{(-/-)}$, *Itgb3*$^{(-/-)}$ and *Apoe*$^{(-/-)}$, *Itgb3*$^{(+/+)}$ BM cells, focusing on monocyte/macrophage cluster**. BM was isolated from *Apoe*$^{(-/-)}$ mice carrying *Itgb3*$^{(-/-)}$ or *Itgb3*$^{(+/+)}$ and subjected to scRNA-seq. **a,** Proportion of leukocyte sub-types in BM cells from scRNA-seq data. **b-h,** t-SNE plots of pooled scRNA-seq data from BM of both genotypes showing the expression of monocyte/macrophage markers. **i**, IPA predicts the effects of differentially expressed genes on the inflammatory responses and monocyte/macrophage migration or proliferation. n=3 mice pooled per genotype. **j,** Differential abundance analysis of sub-clustered monocytes/macrophages show abundance of *Apoe*$^{(-/-)}$, *Itgb3*$^{(-/-)}$ cells in circled regions. **k, l,** Dot plot analysis of monocytes/macrophage sub-cluster showing differentially expressed genes in TNFα and NF-kB signaling pathways, respectively. Dot size represents the fraction of cells expressing the gene, and blue color represents average gene expression.

**Figure S7: Aged BM, integrin β3, TNF and SMC biology. a,** Lysates of $CD3^-$ $CD19^-$ $Cd11b^+$ $Ly6C^+$ monocytes isolated from the BM of young or aged mice were subjected to qRT-PCR to measure Tnfa and IL-1b transcript levels. n=3 mice per age group. **b,** Young *Ldlr*$^{(-/-)}$ recipients were transplanted with young or aged BM, and 4 weeks later, plasma TNFα levels were quantified by ELISA. n=5 mice per age group. **c-e,** Young *Ldlr*$^{(-/-)}$*, Myh11-CreER*$^{T2}$*, ROSA26R*$^{(Rb/+)}$ recipient mice were induced with tamoxifen, transplanted with aged BM and then treated with 12 weeks of WD and concomitant two times per week injections of anti-TNFα antibody (20 mg/kg) or isotype control IgG2a. In **e,** genomic DNA isolated from peripheral blood of recipient mice before (lanes 1-3) or after (lanes 4-6) aged BMT or of WT (lane 7) or *ROSA26R*$^{(Rb/+)}$ (lane 8) mice was amplified using primers for the *Rainbow (Rb)* allele. **f-i,** BM of young or aged wild type mice was harvested and differentiated into macrophages, and isolated mouse aortic SMCs were cultured with macrophage conditioned medium. Anti-TNFα blocking antibody or control IgG was added to conditioned medium, 1 h prior to incubation with SMCs. For migration assay **(f, g)**, confluent SMCs with central acellular area were cultured in conditioned medium for 0 or 8 h. Brightfield images **(f)** and quantification of the percent **(g)** of SMC coverage at 8 h of uncovered area of 0 h are shown, respectively. For proliferation assay **(h, i)**, SMCs were incubated with conditioned medium with EdU for 8 h. SMCs were stained for

EdU and DAPI, and percent of $EdU^+$ cells was quantified. n=3. **j, k**, Mouse aortic SMCs were isolated and treated with Tnfr1 siRNA or scrambled (Scr) RNA. Cell lysates were subjected to Western blot for TNFR1 and GAPDH with densitometry relative to GAPDH and normalized to Scr. n=3 mice. **l-n,** BM cells harvested from aged wild type donor mice were infected with empty vector (control) or *Itgb3* overexpressing (*Itgb3* o/e) lentivirus. After 1 day of infection, aged *Myh11-CreER$^{T2}$*, *ROSA26R$^{(Rb/+)}$* recipient mice were induced with tamoxifen, irradiated, and then transplanted with control and *Itgb3* o/e lentivirus treated BM cells followed by injection with AAV-*Pcsk9* and WD feeding for 12 weeks. In **n,** genomic DNA isolated from peripheral blood of recipient mice before (lanes 1-3) or after (lanes 4-6) BMT or of WT (lane 7) or *ROSA26R$^{(Rb/+)}$* (lane 8) mice was amplified using primers for the *Rainbow (Rb)* allele. **o, p,** after 4 weeks of BMT, fasting total cholesterol and triglycerides levels were assessed in blood plasma, respectively. n=4-6 mice per group. All data are averages ± SD. Student's *t*-test was used. Scale bars, 50 µm **(f)** and 25 µm **(h)**.

**Young**

| Sex | Age (years) | Race/Ethnicity | Relative ITGB3 mRNA level |
|---|---|---|---|
| F | 21 | Hispanic | 0.74 |
| F | 23 | Hispanic | 0.82 |
| F | 23 | Hispanic | 0.93 |
| F | 24 | Asian | 0.88 |
| F | 25 | Hispanic | 1.08 |
| F | 26 | White | 1.16 |
| F | 34 | Hispanic | 1.11 |
| M | 23 | Asian | 1.05 |
| M | 23 | Asian | 1.13 |
| M | 24 | White | 1.06 |
| M | 24 | Black | 0.90 |
| M | 25 | White | 1.12 |
| Avg: | 24.6 | | 1.00 |
| Stdev: | 3.1 | | 0.13 |

**Aged**

| Sex | Age (years) | Race/Ethnicity | Relative ITGB3 mRNA level |
|---|---|---|---|
| M | 50 | Black | 0.28 |
| M | 51 | White | 0.23 |
| M | 52 | Black | 0.23 |
| M | 53 | Asian | 0.50 |
| M | 54 | Asian | 0.26 |
| F | 50 | Hispanic | 0.11 |
| F | 50 | Asian | 0.38 |
| F | 50 | White | 0.36 |
| F | 59 | Asian | 0.19 |
| F | 59 | Hispanic | 0.31 |
| F | 59 | Hispanic | 0.44 |
| F | 73 | Hispanic | 0.26 |
| Avg: | 55.0 | | 0.29 |
| Stdev: | 6.8 | | 0.11 |

**Supplemental Table 1.** Human subjects - RT-qPCR for ITGB3 mRNA in isolated circulating monocytes. Values are normalized to young.

| Gene | Primers |
|---|---|
| *Ldlr* | Primer 1: AATCCATCTTGTTCAATGGCCGATC<br>Primer 2: CCATATGCATCCCCAGTCTT<br>Primer 3: GCGATGGATACACTCACTGC |
| *Tet2* | Primer 1: AGCTGATGGAAAATGCAAGC<br>Primer 2: TCTCAGAGCAAAGAGGACTGC<br>Primer 3: GCCACTTTAGAAGCCTATTGGA |
| *Itgb3* | Primer 1: ATGAAACCAGGAGGAAAGCA<br>Primer 2: GCCTGGGTGCTCGGTTTA<br>Primer 3: GCCAGAGGCCACTTGTGTAG |
| *Rainbow (Rb)* | Primer 1: CTCTGCTGCCTCCTGGCTTCT<br>Primer 2: CGAGGCGGATCACAAGCAATA<br>Primer 3: TCAATGGGCGGGGGTCGTT |

**Supplemental Table 2.** Primer sequences used for genotyping to assess bone marrow reconstitution.

| **Gene** | **Primers** |
|---|---|
| *Itgb3* | Forward: CCACACGAGGCGTGAACTC<br>Reverse: CTTCAGGTTACATCGGGGTGA |
| *Tet2* | Forward: AGAGAAGACAATCGAGAAGTCGG<br>Reverse: CCTTCCGTACTCCCAAACTCAT |
| *Gapdh* | Forward: AGGTCGGTGTGAACGGATTTG<br>Reverse: TGTAGACCATGTAGTTGAGGTCA |
| *Tnfa* | Forward: CCCTCACACTCAGATCATCTTCT<br>Reverse: GCTACGACGTGGGCTACAG |
| *Il-1b* | Forward: GCAACTGTTCCTGAACTCAACT<br>Reverse: ATCTTTTGGGGTCCGTCAACT |

**Supplemental Table 3.** Primer pair sequences used for quantitative reverse transcription polymerase chain reactions.

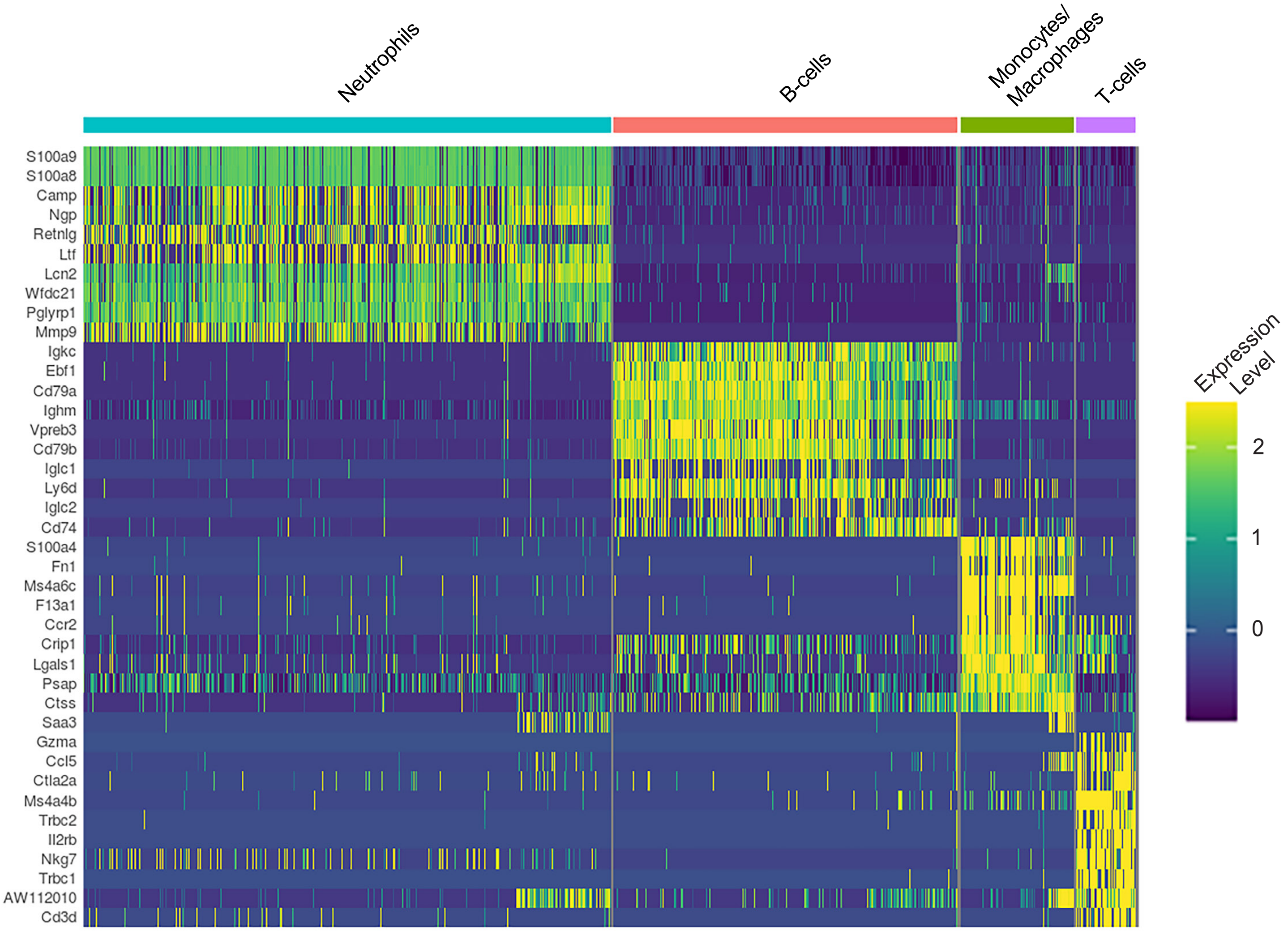


**Supplemental Table 4.** Cluster markers identified by scRNA-seq of BM-derived cells from *Apoe(-/-), Itgb3(+/+)* and *Apoe(-/-), Itgb3(-/-)* mice.

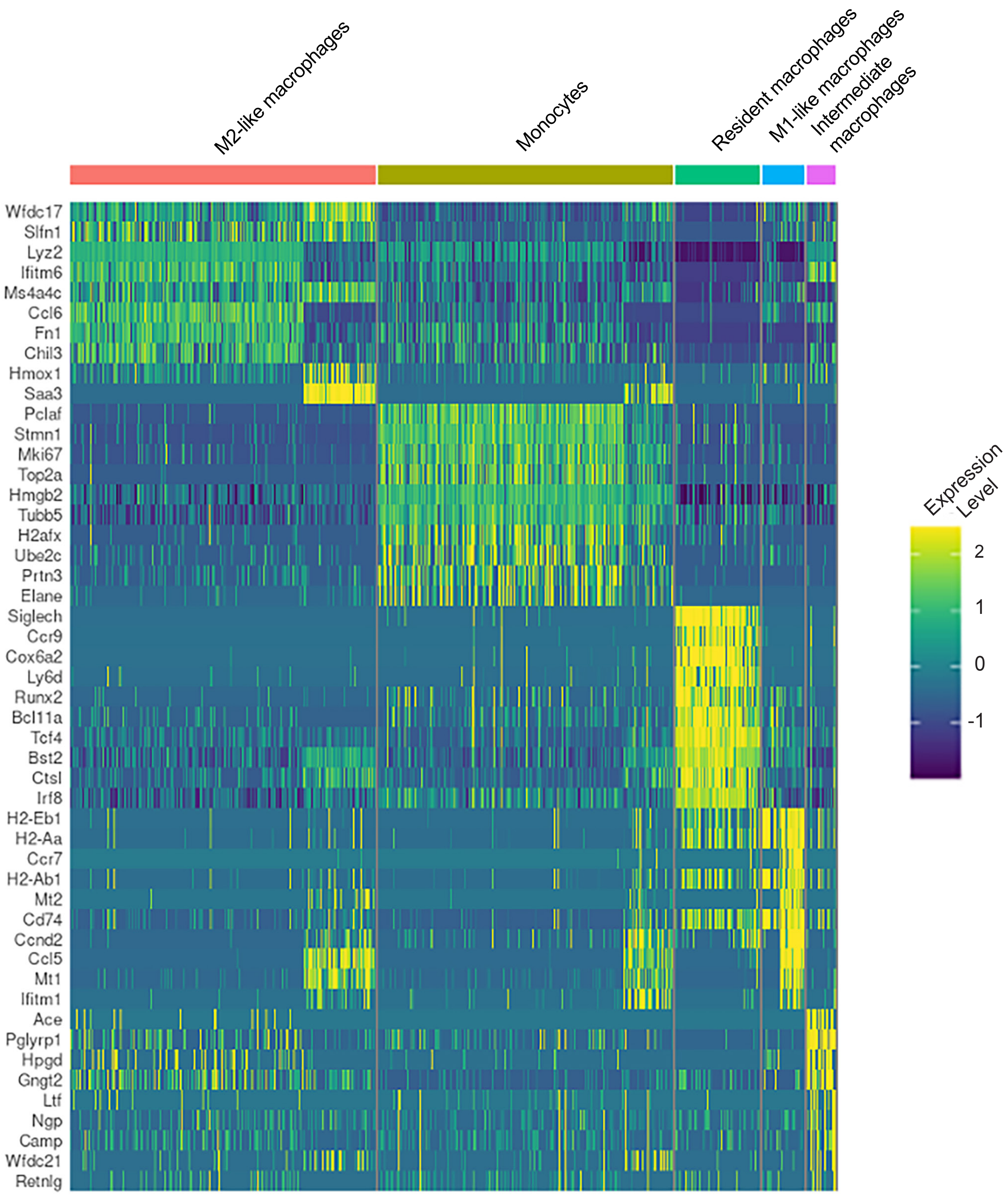


**Supplemental Table 5.** Markers of monocyte/macrophage sub-clusters identified by scRNA-seq of BM cells from *Apoe(-/-), Itgb3(+/+)* and *Apoe(-/-), Itgb3(-/-)* mice.

**Supplemental Methods**

**Plasma lipids and TNFα measurement**

Mice were fasted for 12-16 h and then blood was collected by retro-orbital venous plexus puncture. Plasma was separated by centrifugation. Total plasma cholesterol and triglycerides were measured (Thermo Fisher Scientific, Cholesterol E kit, NC9138103 and Infinity Triglycerides Reagents, TR22421) according to instructions from the manufacturer. Plasma TNFα was quantified by ELISA as per instructions of the manufacturer (Thermo Fisher Scientific, BMS607-2INST).

**Blood leukocyte analysis**

Blood was collected by retro-orbital puncture in heparinized micro-hematocrit capillary tubes. Erythrocytes were lysed with ACK lysis buffer (Thermo Fischer Scientific, A1049201; 155 mM ammonium chloride, 10 mM potassium bicarbonate, 0.01 mM EDTA, pH 7.4). Leukocytes were resuspended in 3% FBS in PBS, and then stained with a cocktail of antibodies (all 1:300, BioLegend): FITC anti-Ly6C (AL-21), PE anti-CD115 (AFS98), APC anti-Ly6G (1A8), Pacific blue anti-CD4 (GK1.5) and Alexa Fluor 647 anti-CD8 (YTS156.7.7). Monocytes were identified as $CD115^{hi}$ and subsets as $Ly6C^{hi}$ and $Ly6C^{lo}$; neutrophils were identified as $CD115^{lo}Ly6C^{hi}Ly6G^{hi}$. In addition. B and T cells were identified with $CD4^{+}$ and $CD8^{+}$ staining, respectively.

**Phalloidin staining for dorsal ruffle**

PDGF-BB (Millipore Sigma, SRP3229) at 10 ng/ml dissolved in DMEM medium containing 0.1% FBS was added to SMCs. Cells were fixed with 4% formaldehyde and permeabilized with PBS-T. F-actin for dorsal ruffle formation was stained using FITC-phalloidin according to manufacturer's protocol (Invitrogen, F432). Cells with dorsal ruffles were counted and represented as a percentage of total cells.

**siRNA-mediated knockdown**

Isolated murine aortic SMCs were transfected with Lipofectamine RNAiMAX (Invitrogen) containing siRNAs (Origene; 50 nM) targeting Itgb3, Tnfr1 or Scr RNA for 6 h. Cells were then washed in PBS and cultured in DMEM supplemented with 10% FBS for 72 h prior to analysis.
