## Supplementary material for "Age of the bone marrow dictates clonality of smooth muscle-derived cells in the atherosclerotic plaque": https://www.biorxiv.org/content/biorxiv/early/2022/01/21/2022.01.18.476756/DC2/embed/media-2.pdf?download=true

**Dataset S1:** Bulk-RNA sequencing of bone marrow-derived monocytes isolated from aged mice relative to young mice

| gene | baseMean | log2FoldChange | lfcSE | stat | pvalue | padj | Symbol |
|---|---|---|---|---|---|---|---|
| ENSMUSG00000109491.1 | 292.9893828 | -24.22098532 | 4.784915417 | -5.061946389 | 4.15E-07 | 5.59E-05 | Gm15396 |
| ENSMUSG00000091228.1 | 204.4615139 | -23.71434455 | 4.784993093 | -4.955983025 | 7.20E-07 | 9.19E-05 | Gm20390 |
| ENSMUSG00000112508.1 | 1576.100152 | -13.98093761 | 1.450685503 | -9.637469724 | 5.55E-22 | 3.43E-19 | Septin2 |
| ENSMUSG00000098650.1 | 805.438148 | -9.606132361 | 2.919125697 | -3.290756671 | 0.00099918 | 0.039275579 | Gm28048 |
| ENSMUSG00000096108.2 | 74.55802999 | -9.578774465 | 1.529602543 | -6.262263688 | 3.79E-10 | 8.03E-08 | Ighv11-2 |
| ENSMUSG00000109129.1 | 63.14509459 | -9.339273437 | 1.544158375 | -6.048131841 | 1.47E-09 | 2.75E-07 | Gm44973 |
| ENSMUSG00000075593.10 | 51.36384795 | -9.042167853 | 1.565706832 | -5.7751347 | 7.69E-09 | 1.36E-06 | Gal3st4 |
| ENSMUSG00000076655.2 | 44.94952316 | -8.851320447 | 1.584613523 | -5.58579131 | 2.33E-08 | 3.79E-06 | Ighv4-1 |
| ENSMUSG00000076576.2 | 25.39219995 | -8.028829413 | 1.691870035 | -4.745535561 | 2.08E-06 | 0.000231904 | Igkv6-32 |
| ENSMUSG00000090009.1 | 20.80401349 | -7.742119045 | 1.746877349 | -4.431976319 | 9.34E-06 | 0.000833553 | Gm16282 |
| ENSMUSG00000094094.2 | 20.56618158 | -7.724894726 | 1.745836472 | -4.424752748 | 9.66E-06 | 0.000846637 | Igkv5-45 |
| ENSMUSG00000031596.15 | 15.74016321 | -7.33935039 | 1.830448533 | -4.009591233 | 6.08E-05 | 0.004192326 | Slc7a2 |
| ENSMUSG00000072663.12 | 11.70829806 | -6.909299292 | 1.937235745 | -3.566576402 | 0.00036168 | 0.017688672 | Spef2 |
| ENSMUSG00000096326.2 | 10.66089511 | -6.781620096 | 2.043059337 | -3.319345636 | 0.00090229 | 0.036532752 | Ighv1-78 |
| ENSMUSG00000058626.16 | 10.1852313 | -6.713858573 | 2.032872289 | -3.30264651 | 0.00095777 | 0.038256603 | Capn11 |
| ENSMUSG00000036172.17 | 9.421913744 | -6.592692752 | 2.047482417 | -3.219902011 | 0.00128234 | 0.047626714 | Cd200r3 |
| ENSMUSG00000092544.8 | 23.85004384 | -6.478060267 | 1.769907369 | -3.660112602 | 0.0002521 | 0.013154703 | Gm20422 |
| ENSMUSG00000094345.2 | 360.02342 | -5.68570083 | 0.367254197 | -15.48164969 | 4.61E-54 | 1.71E-50 | Igkv14-126 |
| ENSMUSG00000037982.16 | 248.5527983 | -5.528452609 | 0.42712815 | -12.94331131 | 2.56E-38 | 3.17E-35 | Gm9725 |
| ENSMUSG00000096458.7 | 22.66470733 | -4.049102535 | 1.165221813 | -3.474962871 | 0.00051092 | 0.022736888 | Moap1 |
| ENSMUSG00000087221.3 | 45.61371452 | -3.280530149 | 0.701350726 | -4.677446002 | 2.90E-06 | 0.000311919 | BC037032 |
| ENSMUSG00000095351.2 | 73.17685787 | -2.91483916 | 0.614299161 | -4.744983139 | 2.09E-06 | 0.000231904 | Igkv3-2 |
| ENSMUSG00000052419.7 | 766.3763481 | -2.332949885 | 0.188114669 | -12.40174353 | 2.56E-35 | 2.53E-32 | 2610001J05Rik |
| ENSMUSG00000069273.2 | 50.95383169 | -1.975651923 | 0.598700335 | -3.299901149 | 0.00096719 | 0.03832292 | Hist1h3e |
| ENSMUSG00000019066.13 | 5674.947453 | -1.714684146 | 0.0962433 | -17.81614038 | 5.30E-71 | 3.92E-67 | Rab3d |
| ENSMUSG00000074221.12 | 214.7500469 | -1.504242092 | 0.408284634 | -3.684297593 | 0.00022933 | 0.012137511 | Zfp568 |
| ENSMUSG00000085754.7 | 268.6160733 | -1.474875168 | 0.377584073 | -3.906084167 | 9.38E-05 | 0.006017659 | Gm15886 |
| ENSMUSG00000029863.13 | 1225.980957 | -1.31772001 | 0.146821414 | -8.974985121 | 2.83E-19 | 1.50E-16 | Casp2 |
| ENSMUSG00000060012.8 | 318.5569964 | -1.285966388 | 0.247086656 | -5.20451574 | 1.95E-07 | 2.77E-05 | Kif13b |
| ENSMUSG00000064367.1 | 4193.204885 | -1.264855972 | 0.098242141 | -12.87488198 | 6.23E-38 | 7.11E-35 | mt-Nd5 |

| | | | | | | | |
|---|---|---|---|---|---|---|---|
| ENSMUSG00000064341.1 | 5785.244515 | -1.206934044 | 0.107089888 | -11.27028949 | 1.84E-29 | 1.43E-26 | mt-Nd1 |
| ENSMUSG00000020231.15 | 222.2813569 | -1.155521354 | 0.352862514 | -3.274707027 | 0.00105772 | 0.040712452 | Dip2a |
| ENSMUSG00000038991.17 | 4937.691126 | -1.141973569 | 0.097789583 | -11.67786524 | 1.65E-31 | 1.36E-28 | Txndc5 |
| ENSMUSG00000000561.14 | 861.9935301 | -1.12094279 | 0.249933477 | -4.484964569 | 7.29E-06 | 0.000663001 | Wdr77 |
| ENSMUSG00000004317.14 | 524.1491669 | -1.078904775 | 0.239230016 | -4.509905539 | 6.49E-06 | 0.000616095 | Clcn5 |
| ENSMUSG00000098178.1 | 31648.30044 | -1.050631105 | 0.080647332 | -13.027475 | 8.54E-39 | 1.15E-35 | Gm42418 |
| ENSMUSG00000106106.2 | 62456.07562 | -1.029068919 | 0.151870391 | -6.775968053 | 1.24E-11 | 3.33E-09 | CT010467.1 |
| ENSMUSG00000033826.9 | 236.0813443 | -1.012844893 | 0.280503145 | -3.610814747 | 0.00030524 | 0.015588469 | Dnah8 |
| ENSMUSG00000030987.5 | 1583.667312 | -0.978351834 | 0.134585658 | -7.269361774 | 3.61E-13 | 1.12E-10 | Stim1 |
| ENSMUSG00000020893.17 | 305.8968082 | -0.978225767 | 0.280743294 | -3.484413648 | 0.00049322 | 0.022151712 | Per1 |
| ENSMUSG00000026924.17 | 1677.480712 | -0.976795388 | 0.214972796 | -4.543809288 | 5.52E-06 | 0.0005458 | Sec16a |
| ENSMUSG00000068551.12 | 462.6830634 | -0.955496075 | 0.209094635 | -4.569682408 | 4.88E-06 | 0.000489091 | Zfp467 |
| ENSMUSG00000064345.1 | 1919.592054 | -0.948170779 | 0.145458738 | -6.518486216 | 7.10E-11 | 1.73E-08 | mt-Nd2 |
| ENSMUSG00000031431.13 | 4062.393824 | -0.946316418 | 0.182741158 | -5.178452563 | 2.24E-07 | 3.16E-05 | Tsc22d3 |
| ENSMUSG00000058773.2 | 778.01086 | -0.945883054 | 0.221857648 | -4.263468322 | 2.01E-05 | 0.001666335 | Hist1h1b |
| ENSMUSG00000030539.13 | 300.4493023 | -0.940844312 | 0.250409673 | -3.757220326 | 0.00017181 | 0.009830388 | Sema4b |
| ENSMUSG00000002320.15 | 1165.669489 | -0.923274251 | 0.145850371 | -6.330283869 | 2.45E-10 | 5.41E-08 | Tm9sf1 |
| ENSMUSG00000064368.1 | 1292.877194 | -0.910269968 | 0.139054207 | -6.546151936 | 5.90E-11 | 1.48E-08 | mt-Nd6 |
| ENSMUSG00000039512.11 | 249.6029958 | -0.901821583 | 0.271635895 | -3.319964705 | 0.00090029 | 0.036532752 | Uhrf1bp1 |
| ENSMUSG00000061482.7 | 1266.705978 | -0.893054449 | 0.228202554 | -3.913428815 | 9.10E-05 | 0.00586283 | Hist1h4d |
| ENSMUSG00000025153.9 | 1428.777441 | -0.852387636 | 0.176061859 | -4.841409945 | 1.29E-06 | 0.000150432 | Fasn |
| ENSMUSG00000064370.1 | 7370.248833 | -0.821571773 | 0.116998877 | -7.022048343 | 2.19E-12 | 6.35E-10 | mt-Cytb |
| ENSMUSG00000028124.15 | 988.6329254 | -0.804435186 | 0.236938094 | -3.395128118 | 0.00068597 | 0.029210673 | Gclm |
| ENSMUSG00000039202.12 | 894.4141333 | -0.782145951 | 0.192042685 | -4.072771385 | 4.65E-05 | 0.003341974 | Abhd2 |
| ENSMUSG00000007080.14 | 718.912469 | -0.777740965 | 0.20573406 | -3.78032187 | 0.00015663 | 0.00906655 | Pole |
| ENSMUSG00000031889.9 | 1098.184503 | -0.775816311 | 0.223778585 | -3.466892555 | 0.00052651 | 0.023290704 | D230025D16Rik |
| ENSMUSG00000027630.13 | 4921.047091 | -0.765891112 | 0.180592192 | -4.240997927 | 2.23E-05 | 0.001782511 | Tbl1xr1 |
| ENSMUSG00000064351.1 | 11504.68554 | -0.759868447 | 0.134721041 | -5.640310092 | 1.70E-08 | 2.79E-06 | mt-Co1 |
| ENSMUSG00000073678.4 | 727.6809763 | -0.744737897 | 0.219718475 | -3.389509672 | 0.00070018 | 0.02956105 | Pgap1 |
| ENSMUSG00000001020.8 | 6375.709857 | -0.730767104 | 0.116158072 | -6.291143563 | 3.15E-10 | 6.87E-08 | S100a4 |
| ENSMUSG00000040809.10 | 70957.8328 | -0.724418356 | 0.075660675 | -9.574569074 | 1.02E-21 | 6.06E-19 | Chil3 |
| ENSMUSG00000021144.14 | 605.7604932 | -0.707185224 | 0.185144212 | -3.819645328 | 0.00013364 | 0.007911944 | Mta1 |
| ENSMUSG00000032902.1 | 647.478533 | -0.695329579 | 0.182119346 | -3.817988559 | 0.00013454 | 0.007911944 | Slc16a1 |

| | | | | | | | |
|---|---|---|---|---|---|---|---|
| ENSMUSG00000029478.16 | 1802.217483 | -0.68612807 | 0.183930108 | -3.730373889 | 0.0001912 | 0.010773122 | Ncor2 |
| ENSMUSG00000020689.4 | 1075.048078 | -0.684245513 | 0.160852759 | -4.253862464 | 2.10E-05 | 0.001729825 | Itgb3 |
| ENSMUSG00000010205.11 | 2760.624463 | -0.683397611 | 0.151860846 | -4.500156745 | 6.79E-06 | 0.00064093 | Raver1 |
| ENSMUSG00000027575.19 | 733.3888788 | -0.677231719 | 0.18583434 | -3.644276503 | 0.00026815 | 0.01384544 | Arfgap1 |
| ENSMUSG00000020715.9 | 1558.760586 | -0.676650226 | 0.150548196 | -4.49457544 | 6.97E-06 | 0.000649696 | Ern1 |
| ENSMUSG00000078122.4 | 751.3110853 | -0.673984322 | 0.192349828 | -3.503950742 | 0.00045841 | 0.021096832 | F630028O10Rik |
| ENSMUSG00000064363.1 | 2159.489264 | -0.671269832 | 0.138111687 | -4.86034054 | 1.17E-06 | 0.000138924 | mt-Nd4 |
| ENSMUSG00000026923.15 | 3240.452672 | -0.655459781 | 0.119910388 | -5.46624687 | 4.60E-08 | 7.12E-06 | Notch1 |
| ENSMUSG00000052565.7 | 560.6910903 | -0.653627096 | 0.199239721 | -3.280606358 | 0.00103584 | 0.040375849 | Hist1h1d |
| ENSMUSG00000006050.11 | 1329.481867 | -0.651544711 | 0.137904044 | -4.724623665 | 2.31E-06 | 0.000251206 | Sra1 |
| ENSMUSG00000027878.11 | 12105.5921 | -0.649936745 | 0.118797273 | -5.470973629 | 4.48E-08 | 7.06E-06 | Notch2 |
| ENSMUSG00000031314.18 | 707.3824194 | -0.633183798 | 0.171104963 | -3.700557759 | 0.00021513 | 0.011677447 | Taf1 |
| ENSMUSG00000026469.14 | 1455.512887 | -0.631338586 | 0.169210493 | -3.731084125 | 0.00019066 | 0.010773122 | Xpr1 |
| ENSMUSG00000024085.13 | 1404.683848 | -0.619572341 | 0.180187361 | -3.438489455 | 0.00058497 | 0.02544216 | Man2a1 |
| ENSMUSG00000031010.17 | 2613.376106 | -0.617887282 | 0.12519083 | -4.935563419 | 7.99E-07 | 0.000101225 | Usp9x |
| ENSMUSG00000038914.15 | 1160.946453 | -0.617644644 | 0.156052563 | -3.957926938 | 7.56E-05 | 0.004957354 | Dido1 |
| ENSMUSG00000027510.17 | 854.8093896 | -0.61601233 | 0.161316518 | -3.818656246 | 0.00013418 | 0.007911944 | Rbm38 |
| ENSMUSG00000054843.8 | 1906.386747 | -0.61571118 | 0.122035205 | -5.045357054 | 4.53E-07 | 5.99E-05 | Atrnl1 |
| ENSMUSG00000060938.14 | 2108.859594 | -0.608821264 | 0.163479247 | -3.724150156 | 0.00019597 | 0.010917829 | Rpl26 |
| ENSMUSG00000045382.6 | 3050.665071 | -0.608098667 | 0.168615734 | -3.606417102 | 0.00031045 | 0.015701766 | Cxcr4 |
| ENSMUSG00000057278.8 | 1497.55208 | -0.607655505 | 0.13219663 | -4.59660361 | 4.29E-06 | 0.000435875 | Snrpg |
| ENSMUSG00000055670.14 | 1483.417014 | -0.606564325 | 0.152812328 | -3.969341552 | 7.21E-05 | 0.004876839 | Zzef1 |
| ENSMUSG00000032661.9 | 2624.974047 | -0.601319436 | 0.170831371 | -3.519959083 | 0.00043161 | 0.020113455 | Oas3 |
| ENSMUSG00000040423.10 | 664.0838269 | -0.597342419 | 0.181302485 | -3.29472825 | 0.00098517 | 0.038827711 | Rc3h1 |
| ENSMUSG00000032518.6 | 7771.673038 | -0.596598613 | 0.160058182 | -3.727385912 | 0.00019348 | 0.010841357 | Rpsa |
| ENSMUSG00000020653.12 | 899.6958095 | -0.595841519 | 0.175648859 | -3.39223107 | 0.00069326 | 0.029352607 | Klf11 |
| ENSMUSG00000037369.17 | 837.4233914 | -0.595044704 | 0.183256444 | -3.247060191 | 0.00116604 | 0.044080373 | Kdm6a |
| ENSMUSG00000019726.11 | 3338.637852 | -0.59348553 | 0.123512699 | -4.805056769 | 1.55E-06 | 0.000176355 | Lyst |
| ENSMUSG00000054008.8 | 2305.92856 | -0.592626295 | 0.139359001 | -4.252515369 | 2.11E-05 | 0.001730652 | Ndst1 |
| ENSMUSG00000066440.5 | 866.6462464 | -0.586265025 | 0.180515398 | -3.247728619 | 0.0011633 | 0.044080373 | Zfyve26 |
| ENSMUSG00000037487.6 | 1376.410412 | -0.58577008 | 0.176762141 | -3.313888803 | 0.00092008 | 0.036950349 | Ubr5 |
| ENSMUSG00000005893.14 | 1729.636017 | -0.577382769 | 0.137586821 | -4.196497628 | 2.71E-05 | 0.002092214 | Nr2c2 |
| ENSMUSG00000027009.18 | 6284.715085 | -0.56658276 | 0.147113547 | -3.851329614 | 0.00011748 | 0.00716424 | Itga4 |

| | | | | | | | |
|---|---|---|---|---|---|---|---|
| ENSMUSG00000034269.12 | 1065.621255 | -0.551652477 | 0.158107928 | -3.489088016 | 0.00048467 | 0.021964365 | Setd5 |
| ENSMUSG00000056602.11 | 1284.917066 | -0.550282026 | 0.155776308 | -3.532514231 | 0.00041163 | 0.019426485 | Fry |
| ENSMUSG00000030660.9 | 1190.234376 | -0.550262673 | 0.142835109 | -3.852432903 | 0.00011695 | 0.007161498 | Pik3c2a |
| ENSMUSG00000042284.10 | 2917.179908 | -0.549520672 | 0.130937001 | -4.19683259 | 2.71E-05 | 0.002092214 | Itga1 |
| ENSMUSG00000079017.3 | 3172.531713 | -0.547702358 | 0.117937847 | -4.6439915 | 3.42E-06 | 0.000356639 | Ifi27l2a |
| ENSMUSG00000074129.13 | 2190.660038 | -0.547597537 | 0.128983046 | -4.245500126 | 2.18E-05 | 0.001766185 | Rpl13a |
| ENSMUSG00000036781.13 | 2180.426397 | -0.545892715 | 0.134978263 | -4.044300936 | 5.25E-05 | 0.003703302 | Rps27l |
| ENSMUSG00000003226.7 | 1874.79543 | -0.543324711 | 0.14307412 | -3.797505163 | 0.00014616 | 0.008527327 | Ranbp2 |
| ENSMUSG00000037805.14 | 1521.570747 | -0.531296355 | 0.129287522 | -4.109417126 | 3.97E-05 | 0.002939045 | Rpl10a |
| ENSMUSG00000022533.13 | 3061.140497 | -0.531004799 | 0.108593739 | -4.889828851 | 1.01E-06 | 0.000122589 | Atp13a3 |
| ENSMUSG00000023143.10 | 1862.541687 | -0.528871626 | 0.134633171 | -3.928241624 | 8.56E-05 | 0.005586124 | Nagpa |
| ENSMUSG00000062328.7 | 1822.646142 | -0.527360059 | 0.133220465 | -3.958551404 | 7.54E-05 | 0.004957354 | Rpl17 |
| ENSMUSG00000024897.8 | 1067.665886 | -0.526602057 | 0.147477547 | -3.570726993 | 0.00035599 | 0.017468352 | Apba1 |
| ENSMUSG00000009470.17 | 2472.3563 | -0.525008476 | 0.147411433 | -3.561518025 | 0.00036872 | 0.0179148 | Tnpo1 |
| ENSMUSG00000074398.5 | 2744.869158 | -0.523241171 | 0.111588282 | -4.689033324 | 2.74E-06 | 0.000296919 | Gm15441 |
| ENSMUSG00000002748.7 | 2669.751071 | -0.521769513 | 0.110007278 | -4.743045404 | 2.11E-06 | 0.000231904 | Baz1b |
| ENSMUSG00000050931.7 | 3016.093209 | -0.510889258 | 0.115331027 | -4.429764239 | 9.43E-06 | 0.000837106 | Sgms2 |
| ENSMUSG00000024681.11 | 1618.82303 | -0.50852761 | 0.132600414 | -3.835037883 | 0.00012555 | 0.007562807 | Ms4a3 |
| ENSMUSG00000022314.9 | 3228.575079 | -0.499113852 | 0.125786465 | -3.967945609 | 7.25E-05 | 0.004883189 | Rad21 |
| ENSMUSG00000024590.8 | 3724.244194 | -0.495127034 | 0.118757958 | -4.169211404 | 3.06E-05 | 0.002334797 | Lmnb1 |
| ENSMUSG00000025362.6 | 2780.104115 | -0.490907197 | 0.130826342 | -3.752357443 | 0.00017518 | 0.009984555 | Rps26 |
| ENSMUSG00000035181.6 | 2154.040387 | -0.488145942 | 0.1287552 | -3.791271664 | 0.00014988 | 0.008709963 | Heatr5a |
| ENSMUSG00000025290.17 | 1600.163909 | -0.487653661 | 0.13246081 | -3.681493886 | 0.00023187 | 0.012228119 | Rps24 |
| ENSMUSG00000028234.6 | 1666.652236 | -0.483937357 | 0.131042645 | -3.692976124 | 0.00022165 | 0.011814946 | Rps20 |
| ENSMUSG00000045128.9 | 8442.070473 | -0.483700393 | 0.117125223 | -4.129771386 | 3.63E-05 | 0.002731541 | Rpl18a |
| ENSMUSG00000022587.14 | 29533.36405 | -0.48179772 | 0.140792274 | -3.422046584 | 0.00062152 | 0.026773997 | Ly6e |
| ENSMUSG00000025949.16 | 1089.021213 | -0.481280307 | 0.144674374 | -3.326645172 | 0.00087898 | 0.035982424 | Pikfyve |
| ENSMUSG00000062006.12 | 1739.02524 | -0.47864604 | 0.124683887 | -3.838876478 | 0.0001236 | 0.007475949 | Rpl34 |
| ENSMUSG00000073702.11 | 1821.657846 | -0.478340734 | 0.136164756 | -3.512955533 | 0.00044315 | 0.020586408 | Rpl31 |
| ENSMUSG00000037563.15 | 3459.382158 | -0.476999562 | 0.122248524 | -3.901884022 | 9.54E-05 | 0.006043704 | Rps16 |
| ENSMUSG00000052812.5 | 1582.195117 | -0.47420384 | 0.131483992 | -3.60655189 | 0.00031029 | 0.015701766 | Atad2b |
| ENSMUSG00000038393.14 | 18163.4248 | -0.472858881 | 0.083027676 | -5.695195931 | 1.23E-08 | 2.05E-06 | Txnip |
| ENSMUSG00000049517.8 | 4461.496031 | -0.46771147 | 0.097175909 | -4.813039285 | 1.49E-06 | 0.000170765 | Rps23 |

| | | | | | | | |
|---|---|---|---|---|---|---|---|
| ENSMUSG00000012848.15 | 4823.470398 | -0.467431739 | 0.102769294 | -4.548359946 | 5.41E-06 | 0.000537717 | Rps5 |
| ENSMUSG00000024491.15 | 1125.249539 | -0.456840942 | 0.142548476 | -3.204811133 | 0.00135151 | 0.04981089 | Rbm27 |
| ENSMUSG00000079477.9 | 9321.741995 | -0.453366762 | 0.085934502 | -5.275724572 | 1.32E-07 | 1.92E-05 | Rab7 |
| ENSMUSG00000009376.15 | 1707.352024 | -0.452849892 | 0.132268457 | -3.423717957 | 0.00061771 | 0.026687461 | Met |
| ENSMUSG00000035992.15 | 1226.411761 | -0.448418718 | 0.139013413 | -3.225722678 | 0.00125655 | 0.047022276 | Fnip1 |
| ENSMUSG00000079481.11 | 5304.366802 | -0.445401252 | 0.110120877 | -4.044657708 | 5.24E-05 | 0.003703302 | Nhsl2 |
| ENSMUSG00000024164.15 | 27756.05704 | -0.445321015 | 0.093564203 | -4.759523408 | 1.94E-06 | 0.000219514 | C3 |
| ENSMUSG00000057841.5 | 8950.444083 | -0.444437374 | 0.112171233 | -3.962133256 | 7.43E-05 | 0.004957354 | Rpl32 |
| ENSMUSG00000029322.12 | 13982.55369 | -0.435481573 | 0.09341567 | -4.661761486 | 3.14E-06 | 0.000331855 | Plac8 |
| ENSMUSG00000051344.13 | 2544.340593 | -0.433534993 | 0.130563044 | -3.320503096 | 0.00089855 | 0.036532752 | Plekhm3 |
| ENSMUSG00000061477.4 | 4811.962005 | -0.433172582 | 0.103076537 | -4.202436312 | 2.64E-05 | 0.00205951 | Rps7 |
| ENSMUSG00000026434.12 | 5422.246377 | -0.428137476 | 0.094899349 | -4.511490106 | 6.44E-06 | 0.000615455 | Nucks1 |
| ENSMUSG00000032849.13 | 2519.940397 | -0.424495792 | 0.118067338 | -3.595370224 | 0.00032393 | 0.016022831 | Abcc4 |
| ENSMUSG00000049313.8 | 26190.25235 | -0.424328923 | 0.097950914 | -4.332056789 | 1.48E-05 | 0.001250917 | Sorl1 |
| ENSMUSG00000008668.14 | 3355.120158 | -0.423353026 | 0.101813106 | -4.158138791 | 3.21E-05 | 0.002438304 | Rps18 |
| ENSMUSG00000026180.8 | 12299.14098 | -0.418440511 | 0.098452218 | -4.250188786 | 2.14E-05 | 0.001739119 | Cxcr2 |
| ENSMUSG00000056054.9 | 170211.1264 | -0.418305985 | 0.085221492 | -4.908456451 | 9.18E-07 | 0.000114313 | S100a8 |
| ENSMUSG00000020841.5 | 3474.008976 | -0.417040379 | 0.105195354 | -3.964437239 | 7.36E-05 | 0.004933138 | Cpd |
| ENSMUSG00000040314.2 | 3367.146975 | -0.416483576 | 0.128094518 | -3.251377041 | 0.00114847 | 0.043639083 | Ctsg |
| ENSMUSG00000027435.8 | 1928.528075 | -0.415606878 | 0.123710299 | -3.359517211 | 0.00078079 | 0.032410352 | Cd93 |
| ENSMUSG00000022186.14 | 2068.796662 | -0.413253948 | 0.117851242 | -3.506572716 | 0.00045392 | 0.020955146 | Oxct1 |
| ENSMUSG00000029202.12 | 2573.298376 | -0.411326492 | 0.123708507 | -3.324965303 | 0.0008843 | 0.036100202 | Pds5a |
| ENSMUSG00000041959.14 | 7158.319001 | -0.411156659 | 0.09843637 | -4.176877479 | 2.96E-05 | 0.002269212 | S100a10 |
| ENSMUSG00000068747.14 | 8197.916704 | -0.403465979 | 0.087365183 | -4.618155245 | 3.87E-06 | 0.000395684 | Sort1 |
| ENSMUSG00000008730.17 | 5754.812113 | -0.402808141 | 0.094978566 | -4.24104255 | 2.22E-05 | 0.001782511 | Hipk1 |
| ENSMUSG00000015027.11 | 2504.90254 | -0.398727606 | 0.116772126 | -3.414578626 | 0.00063881 | 0.027359794 | Galns |
| ENSMUSG00000029265.4 | 6391.056398 | -0.397436214 | 0.099916472 | -3.977684629 | 6.96E-05 | 0.004752295 | Dr1 |
| ENSMUSG00000020315.18 | 3601.380925 | -0.394507428 | 0.10689049 | -3.690762661 | 0.00022358 | 0.011875526 | Sptbn1 |
| ENSMUSG00000037361.8 | 2083.15506 | -0.392268239 | 0.122424886 | -3.204154412 | 0.0013546 | 0.04981089 | Sf3b6 |
| ENSMUSG00000031731.16 | 2653.886871 | -0.391662291 | 0.111970366 | -3.497910235 | 0.00046892 | 0.021360913 | Ap1g1 |
| ENSMUSG00000008682.13 | 3763.123042 | -0.389554089 | 0.109340413 | -3.562764 | 0.00036697 | 0.017888608 | Rpl10 |
| ENSMUSG00000012405.16 | 2385.248186 | -0.383890826 | 0.119293992 | -3.218023131 | 0.00129077 | 0.047819953 | Rpl15 |
| ENSMUSG00000066415.4 | 2337.793499 | -0.381128546 | 0.118523665 | -3.215632478 | 0.00130157 | 0.048099815 | Msl2 |

| | | | | | | | |
|---|---|---|---|---|---|---|---|
| ENSMUSG00000030432.12 | 2464.334563 | -0.374686963 | 0.116109101 | -3.227024921 | 0.00125085 | 0.046927295 | Rpl28 |
| ENSMUSG00000022462.6 | 7071.204433 | -0.373621073 | 0.092435558 | -4.041962645 | 5.30E-05 | 0.003722707 | Slc38a2 |
| ENSMUSG00000022283.14 | 11739.61494 | -0.370574645 | 0.102865835 | -3.602504611 | 0.00031517 | 0.015858884 | Pabpc1 |
| ENSMUSG00000008540.11 | 10567.41049 | -0.36734341 | 0.091147533 | -4.030206841 | 5.57E-05 | 0.003878528 | Mgst1 |
| ENSMUSG00000001525.10 | 8496.802475 | -0.361616037 | 0.091338153 | -3.959090768 | 7.52E-05 | 0.004957354 | Tubb5 |
| ENSMUSG00000028367.5 | 7178.213538 | -0.361076566 | 0.103937012 | -3.47399408 | 0.00051277 | 0.022750819 | Txn1 |
| ENSMUSG00000028081.6 | 8846.526348 | -0.354560255 | 0.100964416 | -3.511734832 | 0.00044519 | 0.020616561 | Rps3a1 |
| ENSMUSG00000102051.1 | 12316.38321 | -0.354429443 | 0.092699167 | -3.823437199 | 0.0001316 | 0.007863877 | I830127L07Rik |
| ENSMUSG00000027035.10 | 6362.930218 | -0.352421467 | 0.103866462 | -3.393024671 | 0.00069125 | 0.029351561 | Cers6 |
| ENSMUSG00000038607.12 | 6911.996843 | -0.350305496 | 0.100163273 | -3.497344743 | 0.00046991 | 0.021360913 | Gng10 |
| ENSMUSG00000058546.8 | 4123.76449 | -0.347340253 | 0.101866712 | -3.409752263 | 0.00065022 | 0.027768292 | Rpl23a |
| ENSMUSG00000055024.12 | 3892.651122 | -0.347091271 | 0.107673542 | -3.223552088 | 0.00126611 | 0.047260766 | Ep300 |
| ENSMUSG00000003746.16 | 3352.870134 | -0.34400945 | 0.102029271 | -3.37167408 | 0.00074713 | 0.031187861 | Man1a |
| ENSMUSG00000036550.16 | 3960.578013 | -0.342603612 | 0.098333288 | -3.484106117 | 0.00049378 | 0.022151712 | Cnot1 |
| ENSMUSG00000019122.8 | 19833.37865 | -0.334500521 | 0.093045608 | -3.595016761 | 0.00032437 | 0.016022831 | Ccl9 |
| ENSMUSG00000020125.7 | 23501.86783 | -0.332493228 | 0.077985894 | -4.263504737 | 2.01E-05 | 0.001666335 | Elane |
| ENSMUSG00000003970.8 | 5909.53872 | -0.330933444 | 0.102755986 | -3.22057583 | 0.00127933 | 0.047626714 | Rpl8 |
| ENSMUSG00000048578.11 | 8192.188778 | -0.323139026 | 0.089731324 | -3.601184218 | 0.00031677 | 0.015858884 | Mlec |
| ENSMUSG00000031232.16 | 3913.578297 | -0.322797255 | 0.098832927 | -3.266090196 | 0.00109044 | 0.041863103 | Magt1 |
| ENSMUSG00000047126.17 | 14834.87979 | -0.317241481 | 0.091292458 | -3.475002067 | 0.00051085 | 0.022736888 | Cltc |
| ENSMUSG00000026581.14 | 31330.40537 | -0.307651738 | 0.080329846 | -3.829855943 | 0.00012822 | 0.007692579 | Sell |
| ENSMUSG00000071866.12 | 23297.98781 | -0.288637383 | 0.083797237 | -3.444473745 | 0.00057217 | 0.025011858 | Ppia |
| ENSMUSG00000037926.15 | 9977.748496 | -0.279249249 | 0.085169016 | -3.278765714 | 0.00104262 | 0.040375849 | Ssh2 |
| ENSMUSG00000029622.16 | 24591.19259 | 0.269144725 | 0.078279175 | 3.438267272 | 0.00058545 | 0.02544216 | Arpc1b |
| ENSMUSG00000021939.8 | 45268.89057 | 0.283257638 | 0.079972184 | 3.541951998 | 0.00039718 | 0.018925328 | Ctsb |
| ENSMUSG00000002111.8 | 14902.76124 | 0.297945621 | 0.084574058 | 3.522896148 | 0.00042686 | 0.019954629 | Spi1 |
| ENSMUSG00000040659.3 | 12429.44712 | 0.306945896 | 0.082622968 | 3.715018989 | 0.00020319 | 0.011152035 | Efhd2 |
| ENSMUSG00000023913.17 | 5572.306541 | 0.311546442 | 0.095026488 | 3.278522123 | 0.00104352 | 0.040375849 | Pla2g7 |
| ENSMUSG00000027293.13 | 5591.283452 | 0.311833878 | 0.093425391 | 3.337785118 | 0.00084449 | 0.034762505 | Ehd4 |
| ENSMUSG00000035004.3 | 8916.629462 | 0.313269572 | 0.085367348 | 3.669665048 | 0.00024287 | 0.012762653 | Igsf6 |
| ENSMUSG00000021094.10 | 5341.004203 | 0.321164942 | 0.097626761 | 3.289722414 | 0.00100286 | 0.03931593 | Dhrs7 |
| ENSMUSG00000024621.15 | 38206.22682 | 0.3268328 | 0.079930539 | 4.088960274 | 4.33E-05 | 0.003163171 | Csf1r |
| ENSMUSG00000018008.7 | 9755.024669 | 0.34727373 | 0.093180991 | 3.726873099 | 0.00019387 | 0.010841357 | Cyth4 |

| | | | | | | | |
|---|---|---|---|---|---|---|---|
| ENSMUSG00000052212.6 | 31967.94307 | 0.356212661 | 0.081407722 | 4.37566184 | 1.21E-05 | 0.001043056 | Cd177 |
| ENSMUSG00000035673.10 | 4104.236588 | 0.359452318 | 0.098422936 | 3.652119433 | 0.00026009 | 0.0135235 | Sbno2 |
| ENSMUSG00000026687.14 | 3357.159052 | 0.360974171 | 0.108039115 | 3.341143361 | 0.00083434 | 0.0344404 | Aldh9a1 |
| ENSMUSG00000052298.12 | 5826.999919 | 0.361381297 | 0.097833398 | 3.693843855 | 0.00022089 | 0.011814946 | Cdc42se2 |
| ENSMUSG00000079478.8 | 2938.211295 | 0.363664135 | 0.106156086 | 3.425749271 | 0.00061311 | 0.026566137 | Sssca1 |
| ENSMUSG00000022148.15 | 4565.331144 | 0.367340401 | 0.096196345 | 3.81865239 | 0.00013418 | 0.007911944 | Fyb |
| ENSMUSG00000044229.9 | 2948.844434 | 0.36911483 | 0.111494731 | 3.310603353 | 0.00093095 | 0.037285832 | Nxpe4 |
| ENSMUSG00000033088.18 | 3335.673591 | 0.373441162 | 0.104907496 | 3.55971859 | 0.00037125 | 0.01797905 | Triobp |
| ENSMUSG00000043832.13 | 6452.341884 | 0.387888231 | 0.094673784 | 4.097102843 | 4.18E-05 | 0.003084365 | Clec4a3 |
| ENSMUSG00000047735.14 | 3126.840388 | 0.395172644 | 0.111415228 | 3.546845898 | 0.00038987 | 0.018758187 | Samd9l |
| ENSMUSG00000073411.11 | 84611.00007 | 0.398586844 | 0.088755539 | 4.490839107 | 7.09E-06 | 0.000653633 | H2-D1 |
| ENSMUSG00000017774.19 | 2888.941141 | 0.407320858 | 0.109447171 | 3.721620714 | 0.00019795 | 0.010986496 | Myo1c |
| ENSMUSG00000053560.4 | 7943.676972 | 0.407581993 | 0.092188324 | 4.421188892 | 9.82E-06 | 0.000855661 | Ier2 |
| ENSMUSG00000024659.14 | 26384.17047 | 0.414150583 | 0.094811186 | 4.368161613 | 1.25E-05 | 0.001073279 | Anxa1 |
| ENSMUSG00000027073.5 | 1595.57401 | 0.414736615 | 0.125719601 | 3.298901781 | 0.00097064 | 0.038357052 | Prg2 |
| ENSMUSG00000064267.13 | 2609.313179 | 0.418061694 | 0.115664241 | 3.614442022 | 0.000301 | 0.015434074 | Hvcn1 |
| ENSMUSG00000015656.17 | 16813.5744 | 0.425153563 | 0.083000799 | 5.12228278 | 3.02E-07 | 4.18E-05 | Hspa8 |
| ENSMUSG00000026833.18 | 4576.903544 | 0.433819301 | 0.107644196 | 4.030122543 | 5.57E-05 | 0.003878528 | Olfm1 |
| ENSMUSG00000017144.8 | 3472.829065 | 0.436590183 | 0.108752638 | 4.014525 | 5.96E-05 | 0.004124781 | Rnd3 |
| ENSMUSG00000019960.8 | 6579.503972 | 0.438458357 | 0.097484458 | 4.497725761 | 6.87E-06 | 0.000644197 | Dusp6 |
| ENSMUSG00000033446.7 | 1644.878647 | 0.443331159 | 0.127263677 | 3.483563972 | 0.00049479 | 0.022151712 | Lpar6 |
| ENSMUSG00000009687.14 | 12065.56315 | 0.443899259 | 0.089123648 | 4.980712422 | 6.34E-07 | 8.24E-05 | Fxyd5 |
| ENSMUSG00000056069.8 | 5969.093985 | 0.446805102 | 0.09763214 | 4.576414114 | 4.73E-06 | 0.000476843 | Fam105a |
| ENSMUSG00000022436.15 | 3231.348751 | 0.448668073 | 0.114463153 | 3.919759845 | 8.86E-05 | 0.005735875 | Sh3bp1 |
| ENSMUSG00000055436.18 | 2022.854467 | 0.449759741 | 0.124584643 | 3.610073693 | 0.00030611 | 0.015588469 | Srsf11 |
| ENSMUSG00000000326.12 | 1530.37587 | 0.450329739 | 0.133345084 | 3.377175411 | 0.00073234 | 0.030657049 | Comt |
| ENSMUSG00000038775.14 | 1378.666297 | 0.45138334 | 0.137441388 | 3.284187866 | 0.00102277 | 0.039990468 | Vill |
| ENSMUSG00000021963.17 | 1966.04295 | 0.454528212 | 0.131297811 | 3.461811027 | 0.00053655 | 0.023664256 | Sap18 |
| ENSMUSG00000025236.11 | 6301.80488 | 0.455408668 | 0.118372811 | 3.847240459 | 0.00011946 | 0.007254976 | Adpgk |
| ENSMUSG00000037197.11 | 1535.920201 | 0.461894561 | 0.128339467 | 3.59900638 | 0.00031944 | 0.015884941 | Rbm17 |
| ENSMUSG00000076617.9 | 17612.37876 | 0.465829826 | 0.079438101 | 5.864060482 | 4.52E-09 | 8.06E-07 | Ighm |
| ENSMUSG00000040613.14 | 2628.622737 | 0.470544929 | 0.145721033 | 3.22908039 | 0.00124189 | 0.046709556 | Apobec1 |
| ENSMUSG00000004266.15 | 15097.5119 | 0.486900471 | 0.080937204 | 6.015780716 | 1.79E-09 | 3.32E-07 | Ptpn6 |

| | | | | | | | |
|---|---|---|---|---|---|---|---|
| ENSMUSG00000024013.14 | 1027.987459 | 0.48971037 | 0.150422693 | 3.255561781 | 0.00113168 | 0.043222696 | Fgd2 |
| ENSMUSG00000030214.6 | 12979.37673 | 0.493974495 | 0.085925801 | 5.748849475 | 8.99E-09 | 1.55E-06 | Plbd1 |
| ENSMUSG00000080058.2 | 4300.010261 | 0.498605932 | 0.111687533 | 4.464293556 | 8.03E-06 | 0.00072589 | Gm11175 |
| ENSMUSG00000049550.17 | 1541.383843 | 0.501571328 | 0.135056852 | 3.713779205 | 0.00020419 | 0.011165485 | Clip1 |
| ENSMUSG00000074272.10 | 5093.61524 | 0.506921393 | 0.094369394 | 5.371671615 | 7.80E-08 | 1.19E-05 | Ceacam1 |
| ENSMUSG00000015837.15 | 5115.659513 | 0.507590747 | 0.146757264 | 3.458709523 | 0.00054277 | 0.023867381 | Sqstm1 |
| ENSMUSG00000027422.15 | 1809.702617 | 0.509960161 | 0.14559961 | 3.502483017 | 0.00046094 | 0.02114773 | Rrbp1 |
| ENSMUSG00000037012.18 | 2871.934427 | 0.514207674 | 0.105693755 | 4.865071474 | 1.14E-06 | 0.000136736 | Hk1 |
| ENSMUSG00000071076.6 | 6626.527876 | 0.516078647 | 0.115049798 | 4.485697975 | 7.27E-06 | 0.000663001 | Jund |
| ENSMUSG00000051212.7 | 1706.549456 | 0.519547432 | 0.132364633 | 3.925122759 | 8.67E-05 | 0.005634179 | Gpr183 |
| ENSMUSG00000015363.13 | 1470.718367 | 0.524023519 | 0.159888601 | 3.277428875 | 0.00104757 | 0.040426973 | Trabd |
| ENSMUSG00000031875.7 | 1804.600671 | 0.525478294 | 0.14597256 | 3.599842975 | 0.00031841 | 0.015884941 | Cmtm3 |
| ENSMUSG00000024381.15 | 1000.204506 | 0.532698207 | 0.152733455 | 3.487763746 | 0.00048708 | 0.022006136 | Bin1 |
| ENSMUSG00000101389.1 | 2240.582196 | 0.537235544 | 0.118365188 | 4.538796862 | 5.66E-06 | 0.000555233 | Ms4a4a |
| ENSMUSG00000026864.13 | 13587.32403 | 0.537632706 | 0.14236903 | 3.776331868 | 0.00015916 | 0.009177107 | Hspa5 |
| ENSMUSG00000022214.14 | 2467.522758 | 0.538089304 | 0.113464909 | 4.742341123 | 2.11E-06 | 0.000231904 | Dcaf11 |
| ENSMUSG00000025225.14 | 2130.423784 | 0.54911188 | 0.118021725 | 4.652633894 | 3.28E-06 | 0.000344433 | Nfkb2 |
| ENSMUSG00000053318.7 | 1129.629789 | 0.559299313 | 0.150503619 | 3.716185136 | 0.00020225 | 0.011141982 | Slamf8 |
| ENSMUSG00000034343.14 | 1650.076024 | 0.560208278 | 0.133147825 | 4.207415916 | 2.58E-05 | 0.002025322 | Ube2f |
| ENSMUSG00000024613.16 | 1739.401316 | 0.560843419 | 0.132745509 | 4.224952116 | 2.39E-05 | 0.001884813 | Tcof1 |
| ENSMUSG00000026150.14 | 2120.848432 | 0.561075472 | 0.148900617 | 3.768120519 | 0.00016448 | 0.009447474 | Mff |
| ENSMUSG00000028410.13 | 2617.089497 | 0.56123448 | 0.108680289 | 5.16408713 | 2.42E-07 | 3.38E-05 | Dnaja1 |
| ENSMUSG00000048924.14 | 1248.490801 | 0.561961322 | 0.167524953 | 3.354493244 | 0.00079511 | 0.032912459 | Ccdc125 |
| ENSMUSG00000025134.2 | 4252.159652 | 0.568319687 | 0.0989081 | 5.745936767 | 9.14E-09 | 1.56E-06 | Alyref |
| ENSMUSG00000108596.1 | 12218.99903 | 0.576648877 | 0.136398238 | 4.227685681 | 2.36E-05 | 0.001881115 | Itgam |
| ENSMUSG00000039497.8 | 2013.70601 | 0.578025919 | 0.128718286 | 4.490627834 | 7.10E-06 | 0.000653633 | Dse |
| ENSMUSG00000018381.15 | 2614.322035 | 0.579008054 | 0.13510029 | 4.285764699 | 1.82E-05 | 0.001524696 | Abi3 |
| ENSMUSG00000039208.15 | 3269.525705 | 0.580080765 | 0.10810868 | 5.365718699 | 8.06E-08 | 1.22E-05 | Metrnl |
| ENSMUSG00000021190.14 | 1591.000737 | 0.581224651 | 0.14889954 | 3.903468406 | 9.48E-05 | 0.006043704 | Lgmn |
| ENSMUSG00000040616.3 | 2179.871124 | 0.581512413 | 0.114379258 | 5.08407226 | 3.69E-07 | 5.02E-05 | Tmem51 |
| ENSMUSG00000042817.15 | 806.407104 | 0.582530574 | 0.164987276 | 3.530760614 | 0.00041437 | 0.019431957 | Flt3 |
| ENSMUSG00000032501.8 | 1409.0376 | 0.5829745 | 0.149410746 | 3.901824431 | 9.55E-05 | 0.006043704 | Trib1 |
| ENSMUSG00000025374.13 | 1040.060936 | 0.589442089 | 0.177032713 | 3.329565934 | 0.00086982 | 0.035705772 | Nabp2 |

| | | | | | | | |
|---|---|---|---|---|---|---|---|
| ENSMUSG00000028431.11 | 1303.430357 | 0.593217085 | 0.152888833 | 3.880055038 | 0.00010443 | 0.006475272 | Ikbkap |
| ENSMUSG00000049734.9 | 3533.895395 | 0.594079025 | 0.1453489 | 4.087261931 | 4.36E-05 | 0.003170788 | Trex1 |
| ENSMUSG00000050350.7 | 900.8017269 | 0.598977607 | 0.160023838 | 3.743052387 | 0.0001818 | 0.010322104 | Gpr18 |
| ENSMUSG00000040699.13 | 12149.5478 | 0.600894349 | 0.129969853 | 4.623336374 | 3.78E-06 | 0.000388604 | Limd2 |
| ENSMUSG00000002602.16 | 1564.607407 | 0.602510279 | 0.132988003 | 4.530561161 | 5.88E-06 | 0.000569778 | Axl |
| ENSMUSG00000002981.10 | 4805.524886 | 0.62304229 | 0.096263103 | 6.472285516 | 9.65E-11 | 2.31E-08 | Clptm1 |
| ENSMUSG00000027544.16 | 924.9672157 | 0.626865206 | 0.153219939 | 4.091276952 | 4.29E-05 | 0.003147236 | Nfatc2 |
| ENSMUSG00000029106.14 | 3864.911249 | 0.627442066 | 0.145132331 | 4.32324117 | 1.54E-05 | 0.001294587 | Add1 |
| ENSMUSG00000025144.17 | 698.4881157 | 0.638826236 | 0.192742008 | 3.314411024 | 0.00091836 | 0.036950349 | Cenpx |
| ENSMUSG00000022817.14 | 1935.769681 | 0.63919097 | 0.130533764 | 4.896748166 | 9.74E-07 | 0.000120325 | Itgb5 |
| ENSMUSG00000031613.9 | 4248.007643 | 0.645095647 | 0.100894695 | 6.393751891 | 1.62E-10 | 3.81E-08 | Hpgd |
| ENSMUSG00000095788.6 | 957.5844047 | 0.647683903 | 0.157102763 | 4.122676716 | 3.74E-05 | 0.002788765 | Sirpb1a |
| ENSMUSG00000031304.18 | 3267.635035 | 0.652988927 | 0.102489194 | 6.371295394 | 1.87E-10 | 4.26E-08 | Il2rg |
| ENSMUSG00000030629.15 | 1583.729385 | 0.655625092 | 0.185650585 | 3.531500276 | 0.00041321 | 0.019431957 | Zfand6 |
| ENSMUSG00000027650.12 | 773.3064447 | 0.656408134 | 0.200206716 | 3.278651916 | 0.00104304 | 0.040375849 | Tti1 |
| ENSMUSG00000025017.9 | 1854.902253 | 0.658782679 | 0.134714326 | 4.890219909 | 1.01E-06 | 0.000122589 | Pik3ap1 |
| ENSMUSG00000030103.11 | 663.9947643 | 0.669887987 | 0.205415582 | 3.261135209 | 0.00110967 | 0.042491515 | Bhlhe40 |
| ENSMUSG00000032484.8 | 95555.27452 | 0.674205706 | 0.076541411 | 8.808378287 | 1.27E-18 | 6.49E-16 | Ngp |
| ENSMUSG00000089824.10 | 1145.555558 | 0.680984987 | 0.176689933 | 3.854124436 | 0.00011615 | 0.007141683 | Rbm12 |
| ENSMUSG00000054263.11 | 495.8249464 | 0.69049659 | 0.200055184 | 3.451530608 | 0.00055742 | 0.024438922 | Lifr |
| ENSMUSG00000021756.12 | 2910.228461 | 0.701566005 | 0.1584824 | 4.426775493 | 9.57E-06 | 0.000843732 | Il6st |
| ENSMUSG00000097099.2 | 525.5767488 | 0.710357222 | 0.194699685 | 3.648476481 | 0.0002638 | 0.013668716 | Gm9917 |
| ENSMUSG00000018001.18 | 582.4188379 | 0.716608502 | 0.211912795 | 3.381619797 | 0.0007206 | 0.030327134 | Cyth3 |
| ENSMUSG00000074677.11 | 1525.243693 | 0.72110919 | 0.154406476 | 4.670200427 | 3.01E-06 | 0.0003208 | Sirpb1c |
| ENSMUSG00000038811.13 | 1097.381582 | 0.724682199 | 0.149300152 | 4.853861083 | 1.21E-06 | 0.000142404 | Gngt2 |
| ENSMUSG00000052837.6 | 9377.414587 | 0.725060184 | 0.096203625 | 7.536724151 | 4.82E-14 | 1.74E-11 | Junb |
| ENSMUSG00000008496.19 | 785.4909757 | 0.728343915 | 0.17956939 | 4.056058306 | 4.99E-05 | 0.003555691 | Pou2f2 |
| ENSMUSG00000010067.13 | 772.2645872 | 0.731477535 | 0.16856635 | 4.339404239 | 1.43E-05 | 0.001216772 | Rassf1 |
| ENSMUSG00000030034.11 | 527.5293635 | 0.733311068 | 0.202170546 | 3.627190422 | 0.00028652 | 0.014742949 | Ino80b |
| ENSMUSG00000001034.17 | 1087.366249 | 0.741185333 | 0.22334862 | 3.318513151 | 0.00090498 | 0.036541983 | Mapk7 |
| ENSMUSG00000010110.17 | 1305.90981 | 0.744453183 | 0.135884396 | 5.478577405 | 4.29E-08 | 6.83E-06 | Stx5a |
| ENSMUSG00000034168.7 | 581.4131867 | 0.747316758 | 0.207511529 | 3.60132645 | 0.0003166 | 0.015858884 | Irf2bpl |
| ENSMUSG00000013698.12 | 1228.730638 | 0.754944202 | 0.138124695 | 5.465671434 | 4.61E-08 | 7.12E-06 | Pea15a |

| | | | | | | | |
|---|---|---|---|---|---|---|---|
| ENSMUSG00000024521.7 | 754.6760497 | 0.761326703 | 0.195939779 | 3.885513739 | 0.00010211 | 0.006358081 | Pmaip1 |
| ENSMUSG00000053687.14 | 543.1352563 | 0.785331922 | 0.193365581 | 4.061384238 | 4.88E-05 | 0.003492316 | Dpep2 |
| ENSMUSG00000019850.11 | 612.6080522 | 0.79992513 | 0.246467182 | 3.245564469 | 0.00117218 | 0.044199843 | Tnfaip3 |
| ENSMUSG00000002625.9 | 415.7071868 | 0.804516285 | 0.224105195 | 3.589904668 | 0.0003308 | 0.016286077 | Akap8l |
| ENSMUSG00000032496.7 | 37539.71485 | 0.80826027 | 0.087753195 | 9.210607852 | 3.24E-20 | 1.78E-17 | Ltf |
| ENSMUSG00000029470.15 | 1002.857271 | 0.812914507 | 0.21895668 | 3.712672777 | 0.00020508 | 0.011173197 | P2rx4 |
| ENSMUSG00000003541.6 | 385.9153234 | 0.815301644 | 0.22240237 | 3.66588559 | 0.00024648 | 0.012906884 | Ier3 |
| ENSMUSG00000052684.4 | 3932.167526 | 0.837692817 | 0.134003266 | 6.251286578 | 4.07E-10 | 8.50E-08 | Jun |
| ENSMUSG00000079547.3 | 3171.689997 | 0.844464169 | 0.103334965 | 8.172104849 | 3.03E-16 | 1.36E-13 | H2-DMb1 |
| ENSMUSG00000024397.14 | 894.5169915 | 0.853606974 | 0.161073085 | 5.299500994 | 1.16E-07 | 1.72E-05 | Aif1 |
| ENSMUSG00000034401.16 | 567.3358665 | 0.904072471 | 0.194805569 | 4.640896435 | 3.47E-06 | 0.000359491 | Spata6 |
| ENSMUSG00000006611.15 | 1323.962888 | 0.914313321 | 0.146612135 | 6.236273141 | 4.48E-10 | 9.22E-08 | Hfe |
| ENSMUSG00000026365.15 | 2695.156364 | 0.918362961 | 0.13103785 | 7.008379342 | 2.41E-12 | 6.87E-10 | Cfh |
| ENSMUSG00000029810.15 | 3486.624767 | 0.920748832 | 0.112904493 | 8.15511243 | 3.49E-16 | 1.52E-13 | Tmem176b |
| ENSMUSG00000043953.12 | 249.4572427 | 0.949153238 | 0.277609322 | 3.41902509 | 0.00062846 | 0.026994605 | Ccrl2 |
| ENSMUSG00000032915.6 | 4464.462284 | 0.957833043 | 0.097707676 | 9.803048076 | 1.09E-22 | 7.04E-20 | Adgre4 |
| ENSMUSG00000074622.4 | 1829.443063 | 0.958559147 | 0.121437681 | 7.893424317 | 2.94E-15 | 1.18E-12 | Mafb |
| ENSMUSG00000023367.14 | 1629.832616 | 0.959159949 | 0.128229244 | 7.480040559 | 7.43E-14 | 2.62E-11 | Tmem176a |
| ENSMUSG00000026628.13 | 636.9248032 | 0.976465878 | 0.200436285 | 4.871702136 | 1.11E-06 | 0.0001333 | Atf3 |
| ENSMUSG00000030717.9 | 322.3905975 | 0.98446098 | 0.245975363 | 4.002274735 | 6.27E-05 | 0.004304123 | Nupr1 |
| ENSMUSG00000045087.8 | 624.0155869 | 0.995509338 | 0.188579776 | 5.278982507 | 1.30E-07 | 1.91E-05 | S1pr5 |
| ENSMUSG00000041235.12 | 898.2345933 | 0.999209936 | 0.156878858 | 6.369309083 | 1.90E-10 | 4.26E-08 | Chd7 |
| ENSMUSG00000038179.13 | 590.3487484 | 1.033737596 | 0.228935659 | 4.515406647 | 6.32E-06 | 0.000608112 | Slamf7 |
| ENSMUSG00000037348.15 | 1126.235668 | 1.039993685 | 0.142976135 | 7.273897054 | 3.49E-13 | 1.10E-10 | Paqr7 |
| ENSMUSG00000042558.16 | 558.2728445 | 1.089880385 | 0.190494856 | 5.721311378 | 1.06E-08 | 1.78E-06 | Adprhl2 |
| ENSMUSG00000040552.8 | 256.3498897 | 1.092432798 | 0.295676868 | 3.694684696 | 0.00022016 | 0.011814946 | C3ar1 |
| ENSMUSG00000021796.14 | 205.554136 | 1.09568112 | 0.309094836 | 3.544805647 | 0.0003929 | 0.018842799 | Bmpr1a |
| ENSMUSG00000003418.11 | 218.0761986 | 1.100378696 | 0.310552088 | 3.543298335 | 0.00039516 | 0.018889703 | St8sia6 |
| ENSMUSG00000004709.14 | 587.1423784 | 1.109409384 | 0.187092554 | 5.929735647 | 3.03E-09 | 5.55E-07 | Cd244 |
| ENSMUSG00000058427.10 | 342.3384173 | 1.109413514 | 0.249327348 | 4.449626255 | 8.60E-06 | 0.000772563 | Cxcl2 |
| ENSMUSG00000022122.14 | 200.5344424 | 1.118604242 | 0.315990262 | 3.539995935 | 0.00040013 | 0.019005044 | Ednrb |
| ENSMUSG00000024610.14 | 42257.04628 | 1.131497054 | 0.088621854 | 12.76769785 | 2.48E-37 | 2.63E-34 | Cd74 |
| ENSMUSG00000017754.13 | 1722.727806 | 1.133809059 | 0.12915222 | 8.778858484 | 1.65E-18 | 8.16E-16 | Pltp |

| | | | | | | | |
|---|---|---|---|---|---|---|---|
| ENSMUSG00000014846.12 | 658.8956977 | 1.211032374 | 0.267208664 | 4.53215984 | 5.84E-06 | 0.000569202 | Tppp3 |
| ENSMUSG00000027562.12 | 279.5564902 | 1.220479218 | 0.295625846 | 4.128459109 | 3.65E-05 | 0.0027333 | Car2 |
| ENSMUSG00000024210.2 | 170.3465325 | 1.228028974 | 0.371996836 | 3.301181237 | 0.00096279 | 0.03832292 | Ip6k3 |
| ENSMUSG00000022026.6 | 1156.615093 | 1.232184487 | 0.145111807 | 8.491276577 | 2.04E-17 | 9.77E-15 | Olfm4 |
| ENSMUSG00000036905.8 | 508.9099482 | 1.248741014 | 0.20424765 | 6.113857428 | 9.73E-10 | 1.92E-07 | C1qb |
| ENSMUSG00000051682.15 | 681.6036698 | 1.265447675 | 0.184510795 | 6.858393698 | 6.96E-12 | 1.95E-09 | Treml4 |
| ENSMUSG00000023034.6 | 2486.720461 | 1.27051697 | 0.125539376 | 10.12046585 | 4.48E-24 | 3.02E-21 | Nr4a1 |
| ENSMUSG00000040435.12 | 312.475046 | 1.275735838 | 0.264149628 | 4.829595432 | 1.37E-06 | 0.00015839 | Ppp1r15a |
| ENSMUSG00000024190.6 | 6201.091774 | 1.277242633 | 0.090746361 | 14.07486345 | 5.42E-45 | 8.93E-42 | Dusp1 |
| ENSMUSG00000076609.2 | 3605.272885 | 1.277420058 | 0.107665252 | 11.86473846 | 1.80E-32 | 1.57E-29 | Igkc |
| ENSMUSG00000022885.15 | 293.3280741 | 1.281601621 | 0.313986152 | 4.081713836 | 4.47E-05 | 0.003231613 | St6gal1 |
| ENSMUSG00000021127.7 | 272.800682 | 1.282038018 | 0.380737501 | 3.367249127 | 0.00075922 | 0.031603627 | Zfp36l1 |
| ENSMUSG00000103593.1 | 299.2947338 | 1.284985328 | 0.258318064 | 4.974430779 | 6.54E-07 | 8.43E-05 | Gm37352 |
| ENSMUSG00000036594.14 | 11089.41374 | 1.29412342 | 0.088395894 | 14.64008522 | 1.56E-48 | 3.30E-45 | H2-Aa |
| ENSMUSG00000042502.10 | 617.8957017 | 1.302500795 | 0.341626972 | 3.812640398 | 0.00013749 | 0.008053223 | Cd2bp2 |
| ENSMUSG00000044786.6 | 17518.92382 | 1.311005967 | 0.086853825 | 15.09439526 | 1.76E-51 | 5.23E-48 | Zfp36 |
| ENSMUSG00000051285.17 | 494.7311072 | 1.32872874 | 0.214699855 | 6.188773355 | 6.06E-10 | 1.21E-07 | Pcmtd1 |
| ENSMUSG00000020681.14 | 2929.877663 | 1.337301338 | 0.124064971 | 10.77904044 | 4.32E-27 | 3.05E-24 | Ace |
| ENSMUSG00000024985.18 | 137.1214701 | 1.356133822 | 0.364861897 | 3.716841452 | 0.00020173 | 0.011141982 | Tcf7l2 |
| ENSMUSG00000060600.15 | 401.9297644 | 1.356143803 | 0.222214863 | 6.10284923 | 1.04E-09 | 2.03E-07 | Eno3 |
| ENSMUSG00000022504.9 | 688.5553981 | 1.378436065 | 0.195955308 | 7.03444105 | 2.00E-12 | 5.93E-10 | Ciita |
| ENSMUSG00000038525.16 | 282.141162 | 1.434999105 | 0.282144815 | 5.086037484 | 3.66E-07 | 5.02E-05 | Armc10 |
| ENSMUSG00000060586.10 | 6261.088207 | 1.441219479 | 0.116546365 | 12.36606115 | 3.99E-35 | 3.69E-32 | H2-Eb1 |
| ENSMUSG00000058755.3 | 1266.487634 | 1.448814308 | 0.155748744 | 9.302253543 | 1.37E-20 | 7.84E-18 | Osm |
| ENSMUSG00000038615.17 | 1107.758321 | 1.459904047 | 0.198155487 | 7.367467174 | 1.74E-13 | 5.73E-11 | Nfe2l1 |
| ENSMUSG00000090877.3 | 107.7426511 | 1.510817999 | 0.408369944 | 3.699630743 | 0.00021591 | 0.011677447 | Hspa1b |
| ENSMUSG00000073421.5 | 14945.95709 | 1.517367442 | 0.087509781 | 17.33940394 | 2.37E-67 | 1.17E-63 | H2-Ab1 |
| ENSMUSG00000024742.15 | 1915.97264 | 1.521265931 | 0.242458185 | 6.274343472 | 3.51E-10 | 7.54E-08 | Fen1 |
| ENSMUSG00000053119.11 | 1787.909235 | 1.552919588 | 0.230501894 | 6.737122897 | 1.62E-11 | 4.28E-09 | Chmp3 |
| ENSMUSG00000018474.17 | 91.43418942 | 1.564515113 | 0.462747296 | 3.380927617 | 0.00072242 | 0.030327134 | Chd3 |
| ENSMUSG00000002985.16 | 8800.726017 | 1.568142049 | 0.482297316 | 3.251401153 | 0.00114838 | 0.043639083 | Apoe |
| ENSMUSG00000002944.15 | 473.3030718 | 1.583367226 | 0.205489003 | 7.705362342 | 1.30E-14 | 5.06E-12 | Cd36 |
| ENSMUSG00000051184.7 | 378.6158107 | 1.596999845 | 0.249995296 | 6.388119571 | 1.68E-10 | 3.89E-08 | Zfp524 |

| | | | | | | | |
|---|---|---|---|---|---|---|---|
| ENSMUSG00000056579.17 | 1941.838289 | 1.619176163 | 0.123646915 | 13.09516024 | 3.51E-39 | 5.20E-36 | Tug1 |
| ENSMUSG00000028063.15 | 114.2544952 | 1.694817431 | 0.435876691 | 3.888295624 | 0.00010095 | 0.006312183 | Lmna |
| ENSMUSG00000000982.5 | 404.7240873 | 1.736602111 | 0.255565979 | 6.795122408 | 1.08E-11 | 2.97E-09 | Ccl3 |
| ENSMUSG00000029816.10 | 369.0318858 | 1.806421106 | 0.241893229 | 7.467844845 | 8.15E-14 | 2.81E-11 | Gpnmb |
| ENSMUSG00000056973.6 | 208.7853207 | 1.8505406 | 0.303495813 | 6.097417243 | 1.08E-09 | 2.05E-07 | Ces1d |
| ENSMUSG00000036526.8 | 90.73897596 | 1.879547318 | 0.485705899 | 3.869723057 | 0.00010896 | 0.006727768 | Card11 |
| ENSMUSG00000072109.11 | 97.10132843 | 1.928087975 | 0.436241486 | 4.419772156 | 9.88E-06 | 0.000856252 | A530040E14Rik |
| ENSMUSG00000000957.10 | 59.68967884 | 1.955389608 | 0.59246832 | 3.300412095 | 0.00096543 | 0.03832292 | Mmp14 |
| ENSMUSG00000021250.13 | 8999.65289 | 2.068310066 | 0.100620143 | 20.5556264 | 6.85E-94 | 1.02E-89 | Fos |
| ENSMUSG00000038418.7 | 812.5888081 | 2.140575341 | 0.190418183 | 11.24144399 | 2.55E-29 | 1.89E-26 | Egr1 |
| ENSMUSG00000048498.7 | 129.9867805 | 2.311235952 | 0.443942809 | 5.206156975 | 1.93E-07 | 2.77E-05 | Cd300e |
| ENSMUSG00000026729.9 | 264.0715147 | 2.340176635 | 0.356551728 | 6.563357988 | 5.26E-11 | 1.34E-08 | 4930562F07Rik |
| ENSMUSG00000085009.1 | 80.51704454 | 2.44699649 | 0.496885535 | 4.924668393 | 8.45E-07 | 0.000106124 | Gm12977 |
| ENSMUSG00000026276.18 | 2407.802936 | 2.53544184 | 0.714038234 | 3.550848848 | 0.00038399 | 0.018535377 | 2-Sep |
| ENSMUSG00000091971.3 | 61.86242707 | 2.563534999 | 0.617989387 | 4.148186121 | 3.35E-05 | 0.002533746 | Hspa1a |
| ENSMUSG00000093894.1 | 58.12868869 | 3.475299478 | 0.653444965 | 5.318427207 | 1.05E-07 | 1.57E-05 | Ighv1-53 |
| ENSMUSG00000094652.2 | 25.2206787 | 3.752658443 | 0.964175828 | 3.89208932 | 9.94E-05 | 0.006240597 | Ighv1-42 |
| ENSMUSG00000029165.16 | 719.194785 | 3.757545121 | 0.254956737 | 14.73797147 | 3.68E-49 | 9.08E-46 | Agbl5 |
| ENSMUSG00000105852.4 | 67.11586905 | 4.287239407 | 0.64176432 | 6.68039539 | 2.38E-11 | 6.20E-09 | Gm42890 |
| ENSMUSG00000087612.1 | 329.5179489 | 6.77271051 | 0.47885134 | 14.14365993 | 2.04E-45 | 3.79E-42 | A230005M16Rik |
| ENSMUSG00000104953.1 | 104.5189407 | 6.825925454 | 0.834952295 | 8.175228088 | 2.95E-16 | 1.36E-13 | AC147560.1 |
| ENSMUSG00000076540.3 | 10.9927859 | 6.996958069 | 1.980389041 | 3.533123 | 0.00041068 | 0.019426485 | Igkv4-80 |
| ENSMUSG00000071042.11 | 12.40105342 | 7.170492833 | 1.939463513 | 3.697152735 | 0.00021803 | 0.011749105 | Rasgrp3 |
| ENSMUSG00000097325.2 | 14.26251126 | 7.373778345 | 1.857673019 | 3.96936289 | 7.21E-05 | 0.004876839 | Gm16897 |
| ENSMUSG00000073028.4 | 14.49104387 | 7.395504746 | 1.867078473 | 3.961003705 | 7.46E-05 | 0.004957354 | Igkv4-71 |
| ENSMUSG00000090706.1 | 14.78929582 | 7.428406507 | 2.123682776 | 3.497888946 | 0.00046896 | 0.021360913 | Gm17233 |
| ENSMUSG00000039021.15 | 15.33212946 | 7.480229879 | 1.917571903 | 3.900886254 | 9.58E-05 | 0.006043704 | Ttc16 |
| ENSMUSG00000035934.16 | 16.81786617 | 7.611949854 | 1.801715048 | 4.224835589 | 2.39E-05 | 0.001884813 | Pknox2 |
| ENSMUSG00000095589.2 | 41.39144039 | 8.910521126 | 1.595712543 | 5.584039033 | 2.35E-08 | 3.79E-06 | Ighv1-55 |
| ENSMUSG00000094075.1 | 48.75150332 | 9.147950132 | 1.587154031 | 5.763744383 | 8.23E-09 | 1.43E-06 | Ighv1-80 |
| ENSMUSG00000096459.1 | 51.80044591 | 9.233689913 | 1.572911808 | 5.870443507 | 4.35E-09 | 7.85E-07 | Ighv9-3 |
| ENSMUSG00000076672.7 | 67.0263336 | 9.60573333 | 1.541292807 | 6.232257289 | 4.60E-10 | 9.33E-08 | Ighv3-6 |
| ENSMUSG00000096422.2 | 83.94214232 | 9.930498553 | 1.521638547 | 6.526187559 | 6.75E-11 | 1.67E-08 | Igkv12-44 |

| | | | | | | | |
|---|---|---|---|---|---|---|---|
| ENSMUSG00000095682.1 | 136.3756146 | 10.63117033 | 1.491531752 | 7.12768623 | 1.02E-12 | 3.09E-10 | Igkv3-1 |
| ENSMUSG00000076534.6 | 158.157002 | 10.84460238 | 1.485273213 | 7.301419216 | 2.85E-13 | 9.17E-11 | Igkv12-89 |
| ENSMUSG00000076563.2 | 177.4278563 | 11.0101043 | 1.485318883 | 7.412619893 | 1.24E-13 | 4.17E-11 | Igkv5-48 |
| ENSMUSG00000076665.4 | 194.2257984 | 11.14138614 | 1.478247615 | 7.536887616 | 4.81E-14 | 1.74E-11 | Ighv7-1 |
| ENSMUSG00000096833.2 | 228.4364626 | 11.37471925 | 1.476703026 | 7.702780487 | 1.33E-14 | 5.06E-12 | Igkv4-55 |
| ENSMUSG00000076523.2 | 297.6230553 | 11.7572245 | 1.467863519 | 8.009753191 | 1.15E-15 | 4.73E-13 | Igkv15-103 |
| ENSMUSG00000095079.6 | 302.1654845 | 11.7783379 | 1.469012913 | 8.017858653 | 1.08E-15 | 4.56E-13 | Igha |
| ENSMUSG00000110423.1 | 119.5228741 | 23.87070576 | 4.785228797 | 4.988414719 | 6.09E-07 | 7.98E-05 | Gm45736 |
| ENSMUSG00000092345.1 | 166.5850058 | 24.20016289 | 4.785092367 | 5.05740768 | 4.25E-07 | 5.67E-05 | Gm20503 |
| ENSMUSG00000086324.8 | 5772.705813 | 29.17339327 | 4.784755956 | 6.097153866 | 1.08E-09 | 2.05E-07 | Gm15564 |
